# Structural innovations and neurogenic continuity define avian brain development and evolution

**DOI:** 10.1101/2025.05.30.654767

**Authors:** Hongcheng Shan, Yan Mei, Zhenhuan Feng, Xilin Gu, Liu Huang, Yuan Hui, Chunqiong Li, Yingying Zhou, Junjie He, Ran Wu, Zhengkun Zhuang, Mengzhu Wang, Duoyuan Chen, Yongjie Wu, Li Zhang, Guangshuai Jia

## Abstract

Understanding how neurogenic diversity and brain architecture emerge is crucial for deciphering the mechanisms of brain plasticity and cognitive evolution. Here, we present a comprehensive single-cell and spatial transcriptomic atlas of the budgerigar brain, a highly cognitive avian species. We uncover a striking dorsoventral symmetry in excitatory neuron distribution, precisely organized along the lamina mesopallialis intermedia (LMI), a key boundary structure. Cross-species comparisons further reveal the origins of this unique avian pallial organization, offering new insight into avian brain evolution. Our study also highlights distinct developmental trajectories and asynchronous maturation patterns between telencephalic and optic tectum excitatory neurons, underscoring their contribution to innate circuit assembly. Furthermore, we identify adult neural stem cells (NSCs) with evolutionarily conserved transcriptomic signatures across avian species, emphasizing their role in brain plasticity and adaptation. These findings refine fundamental models of avian brain development and elucidate conserved principles of cognitive plasticity.

## Main Text

Cognitive behaviors such as problem-solving, vocal learning, and social communication are often viewed as hallmarks of mammalian brain evolution, and these abilities are supported by the specialized circuitry of the neocortex (*1*). Avian species, particularly parrots (*Psittacidae*) and corvids (*Corvidae*), exhibit cognitive sophistication comparable to that of mammals, challenging the notion that a layered neocortex is required for advanced cognition (*2–4*). Their abilities are instead supported by distinct pallial structures, high neuronal densities, and a prefrontal cortex-like domain in the avian forebrain (*5–7*). In addition, birds retain robust adult neurogenesis across the brain (*8, 9*), unlike the spatially confined neurogenic niches observed in mammals (*10, 11*). The incorporation of newly generated neurons into pre-existing circuits is hypothesized to support the cognitive flexibility observed in birds (*12, 13*).

Understanding how such divergent architectures support similar functions requires a developmental and evolutionary framework grounded in cellular resolution (*14*). Historically, the avian telencephalon was thought to consist primarily of basal ganglia–like structures based on anatomical and histological observations. This view was overturned by the Avian Brain Nomenclature Consortium, which reclassified large regions of the avian brain as pallial and proposed partial homologies with mammalian cortical structures (*15*). More recently, the continuum model proposes that the avian pallium forms a continuous cellular gradient around the ventricle, rather than being divided into discrete dorsal and ventral regions (*16, 17*). Yet, the developmental and evolutionary mechanisms supporting this organization remain poorly defined.

Single-cell and spatial transcriptomics now offer the resolution needed to revisit long-standing questions in comparative neuroanatomy (*18, 19*). Recent studies in reptiles (*20*), amphibians (*21*), and chicken (*22*) suggest that key pallial subdivisions, including dorsal, medial, lateral, and ventral, are conserved across amniotes, but have diversified through species-specific trajectories of neurogenesis and circuit assembly (*22, 23*). However, a comprehensive, spatiotemporally resolved atlas in a cognitively advanced avian species has been lacking.

Here, we present the first single-cell resolution brain atlas of a cognitively advanced bird, the budgerigar (*Melopsittacus undulatus*), spanning embryonic to adult stages. Using single-nucleus RNA-seq, spatial transcriptomics (Stereo-seq), and in situ validation, we profiled 555,001 nuclei, identified 13 major cell classes, 49 subclasses, 121 neuronal supertypes, and mapped molecular domains across the pallium. Spatial mapping revealed striking dorsoventral symmetry in excitatory neuron organization, marked by graded expression of developmental regulators (*NTS, DACH2*) across hyperpallial and nidopallial territories. We also identified symmetrically distributed *GLRA3*⁺ neurons along the lamina mesopallialis intermedia (LMI), supporting the continuum model of a developmentally fused ventricular zone. This structural innovation enables the coexistence of mammalian neocortex-like input-output circuits along the dorsoventral axis of the avian brain, providing a neuroanatomical substrate for higher cognitive functions. Furthermore, we revealed that the optic tectum exhibits inside-out lamination and asynchronous neurogenesis, resembling the assembly of the mammalian neocortex. Finally, we mapped ongoing adult neurogenesis using 5-ethynyl-2′-deoxyuridine (EdU) labeling coupled with single-cell/nucleus sequencing, revealing conserved neural stem cell signatures and a uniquely expanded neurogenic niche in the avian brain. Together, these results provided a revised model of avian pallial organization and shed light on the evolutionary strategies that support plasticity and cognition across amniotes. To facilitate further exploration, we offered an open-access, high-resolution atlas of the budgerigar brain via an interactive web portal (www.budgiebrain.info; temporarily accessible during review at http://60.204.237.173).

## Cell classes and molecularly defined regions of the budgerigar brain

To chart the cellular taxonomy and molecular architecture of the budgerigar brain, we performed single-nucleus RNA sequencing (snRNA-seq) on four dissected brain regions (the telencephalon, mesencephalon, diencephalon, and cerebellum) across four developmental stages: embryonic day 14 (E14), postnatal day 1 (P1), juvenile, and adult (Fig. 1A, fig. S1, and Data S1). Following stringent quality control (see Materials and Methods), we obtained 555,001 high-quality nuclei, classified into 13 major cell classes including both neuronal and non-neuronal types (Fig. 1B). These cell classes included glutamatergic cells (*SLC17A6*), GABAergic cells (*GAD1, GAD2*), medium spiny neurons (*MEIS2*), Purkinje cells (*CA8*), astroependymal cells (*SPEF2*), astrocytes (*SLC4A4, SLC1A3*), astrocyte precursors (*TOP2A*), oligodendrocytes (*PLP1*), oligodendrocyte precursors (*VCAN*), microglia (*CSF1R*), vascular cells (*FLI1*), mural cells (*RGS5*), and blood cells (*F10*) (Fig. 1B, and fig. S2, A to C). The distribution of these classes varied across stages and brain regions, reflecting the spatiotemporal dynamics of brain organization and maturation (fig. S2, D and E).

**Fig. 1.**
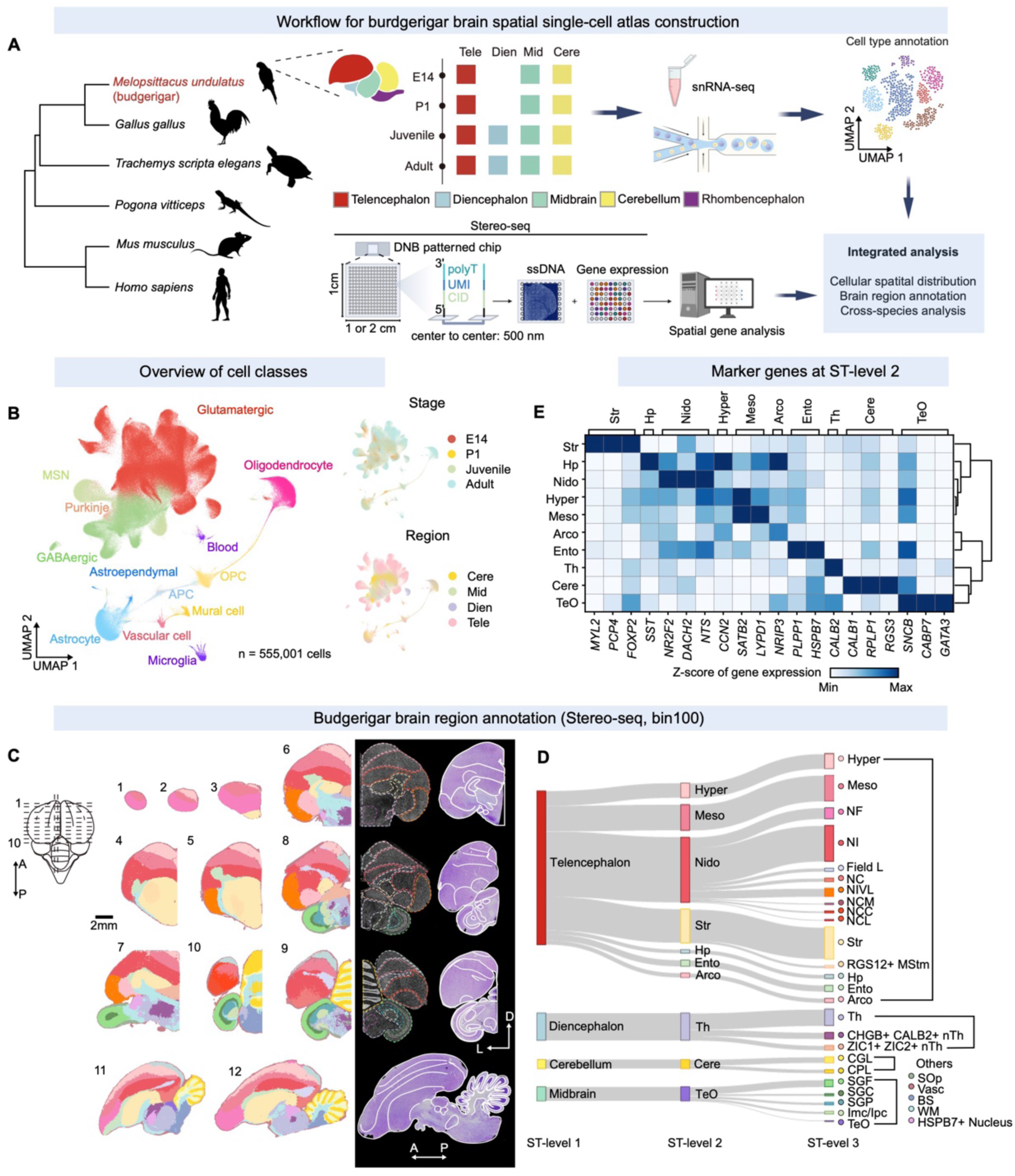
Cell types and spatial brain regions in the budgerigar brain. (**A**) Workflow for constructing the spatial single-cell atlas of the budgerigar (*Melopsittacus undulatus*) brain. snRNA-seq and spatial transcriptomics were performed across developmental stages including embryonic day 14 (E14), postnatal day 1 (P1), juvenile (2 months), and adult (5 months), and major brain regions including telencephalon (Tele), diencephalon (Dien), midbrain (Mid), and cerebellum (Cere). (**B**) UMAP of 555,001 nuclei, colored by cell class (left), developmental stage (top right), or brain region (bottom right). OPCs, oligodendrocyte progenitor cells; MSN, medium spiny neuron. (**C**) Spatial annotation of brain regions. (Left) Sampled brain sections (coronal: 1–10; sagittal: 11–12). (Middle) Regional labels based on ST-level 2 clusters (see **D**). (Right) Combined visualization of spatially resolved mRNA capture (ST data) and corresponding Nissl staining. (**D**) Hierarchical regional annotation based on ST data. ST-level 1 defines gross anatomical domains, ST-level 2 resolves major regional compartments, and ST-level 3 identifies fine subregions. Arrows indicate relationships across levels. (**E**) Expression of region-specific marker genes across ST-level 2 regions. Heatmap shows standardized Z-scores of gene expression.

To align spatial transcriptomic data with classical avian brain anatomy, we first generated a reference atlas of the budgerigar brain using Nissl staining (fig. S3, A to D), providing a cytoarchitectural scaffold for downstream molecular annotation. We then defined molecular regions based on spatial gene expression. Using spatial transcriptomics (ST) technique Stereo-seq, we profiled brain sections from six adult individuals, including ten coronal and two sagittal slices spanning the anterior–posterior axis (Fig. 1, A, C, and D, and fig. S4). For optimal gene detection and clustering, we applied a bin size of 100 (50 μm), followed by Spatial Leiden clustering (*24*) and Gaussian smoothing to enhance signal-to-noise ratios (fig. S5, A and B; see also Materials and Methods), enabling robust delineation of spatial domains (fig. S5C). This analysis revealed a hierarchical three-tiered spatial organization in the adult budgerigar brain: ST-level 1 defined broad anatomical territories; ST-level 2 resolved major regional divisions, especially within the telencephalon; and ST-level 3 captured fine-grained subregions (Fig. 1, C and D). ST-level 2 annotations were validated using canonical marker genes, which aligned with established anatomical domains (Fig. 1E).

To rigorously validate molecular subregions, we analyzed differentially expressed genes (DEGs) and cross-referenced them with published datasets (figs. S6, A to F, and S7, A to C). We further confirmed the spatial expression of five region-specific genes (*SST, NR2F2, NRIP3, GABRA4* and *PPP1R17*) via hybridization chain reaction RNA fluorescence *in situ* hybridization (HCR RNA-FISH) (figs. S6, G to K, S7D, and Data S2). Comparison with Nissl staining showed strong correspondence between molecular and anatomical boundaries, in line with the established nomenclature for the budgerigar brain (*25–27*) (figs. S8, A to C), including major telencephalic domains such as hyperpallium, arcopallium, nidopallium, mesopallium, and entopallium, as well as the laminar structure of the optic tectum. In contrast, molecular subregions within the diencephalon did not consistently map to anatomically defined areas, likely due to convergent gene expression across functionally distinct nuclei. Consequently, the diencephalon was not further analyzed.

## Spatial organization of telencephalic gene expression along medial–lateral and dorsal– ventral axes

The vertebrate telencephalon underlies higher cognitive functions and its organization depends on precise spatial patterning of gene expression (*28*). To identify genes driving spatial segmentation beyond canonical regional boundaries, we focused on spatially restricted genes that are key for cross-species comparisons and tracing the developmental origins of cell types. Inspired by the strategies applied in the goldfish telencephalon atlas (*29*), we systematically mapped the topographical distribution of such genes along the medial–lateral (X) and dorsal–ventral (Y) axes in the budgerigar telencephalon. Spatial specificity was quantified by calculating the spread of each gene’s X and Y coordinates across bin100 spots: genes with narrow spread localized to discrete areas, while broader or scattered expression yielded higher spread values. To capture trends extending beyond single sections, we integrated these spatial scores across coronal planes, identifying consistent anterior–posterior (Z-axis) patterns shared across genes (Fig. 2, A to D). This analysis revealed that axially patterned genes rarely aligned along a single axis and no gene showed exclusive dorsal–ventral (Y) distribution without concurrent medial–lateral (X) enrichment. Instead, most genes exhibited coaxial patterns spanning two or more axes, aligning with both anatomical features and molecularly defined domains. These patterns reflected distinct cellular distributions. For instance, the excitatory marker *SLC17A6* (also known as *VGLUT2*) localized to dorsal pallium-derived regions, while the inhibitory marker *GAD2* (also known as *GAD65*) was enriched ventrally in subpallial territories (Fig. 2B). Likewise, *MBP*, a marker of oligodendrocytes and white matter, clustered medially in arcopallium and entopallium, consistent with Nissl staining (Fig. 2D and fig. S3).

**Fig. 2.**
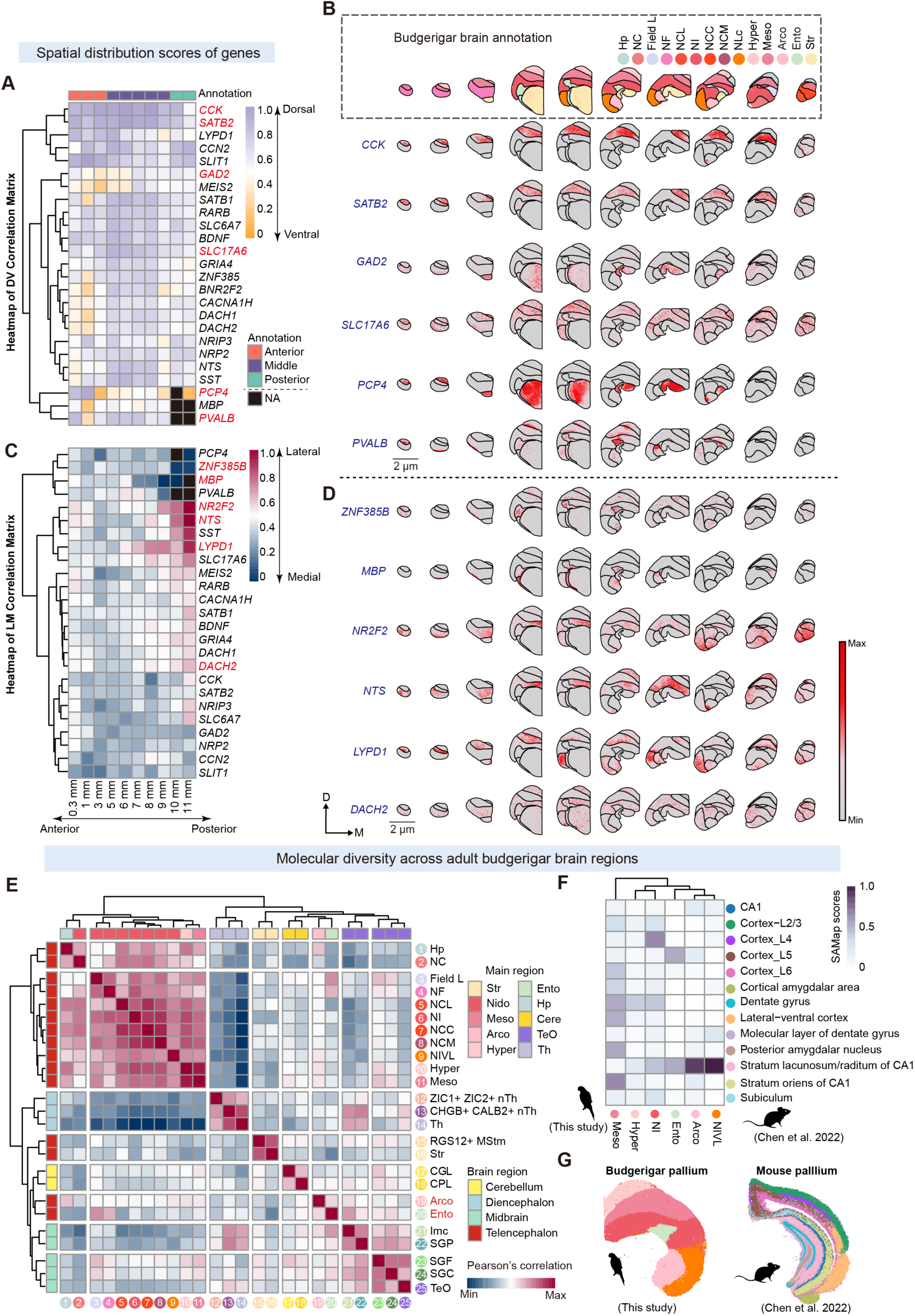
Spatial molecular diversity across adult budgerigar brain regions. (**A-D**) Spatially patterned expression of telencephalic marker genes along dorsal–ventral (**A, B**) and lateral–medial (**C, D**) axes. Left: heatmaps show normalized spatial distribution scores across ten coronal sections (anterior: 0.1, 1, 3 mm; middle: 5-9 mm; posterior: 10-11 mm), where values approaching 1 indicate dorsal (**A**) or lateral (**C**), while values near 0 reflect ventral (**A**) or medial (**C**) localization. Genes with strong axial bias are highlighted in red. Right: spatial expression of selected axially biased genes across anatomical annotation. (**E**) Pearson correlation comparing transcriptomic profiles across ST-level 1 subdivisions. Major brain regions (ST-level 2) are annotated by color. (**F**) Heatmap shows cross-species comparison of molecular architecture between the budgerigar and mouse pallium using SAMap. (**G**) Reference brain maps of the budgerigar (this study) and the mouse (*39*). Colors indicate ST-defined brain regions in (**F**).

Coaxial arrangements varied across genes, even among canonical regional markers. *CCK* and *SATB2*, associated with mammalian cortical neurons (*30, 31*), were confined to dorsal-medial mesopallium, suggesting the presence of intratelencephalic (IT)-like neurons with possible functional homology to mammalian cortex (*32, 33*) (Fig. 2B). The prototoxin *LYPD1*, which modulates nAChR-mediated synaptic transmission (*34*), showed spatially split expression: dorsomedial in anterior telencephalon and ventrolateral in the posterior (Fig. 2D). Similarly, *PCP4* and *PVALB*, typically expressed in deep layers of the mammalian neocortex (*35, 36*), exhibited medial enrichment spanning the ventral striatum and dorsal entopallium (Fig. 2B).

Notably, *ZNF385B*, despite resembling *PVALB* in expression, was nearly absent in field L (Fig. 2D), a posterior region involved in vocal control (*25*). This divergence challenges the traditional view that field L and entopallium comprise a unified intercalated pallial domain (*16, 17*). In parallel, neuropeptides *NTS* and *SST* showed similar Z-axis patterns, with strong expression in posterior structures, including the dorsal hippocampus (Hp) and lateral nidopallium caudolaterale (NCL) (Fig. 2D). Although both *NR2F2* and *DACH2* label nidopallium/hyperpallium domains, their Z-axis distributions diverged: *NR2F2* was absent from the nucleus interstitialis ventralis lateralis (NIVL) region in mid-telencephalon but enriched in posterior NCL, whereas *DACH2* spanned the entire nidopallium, including posterior field L, with additional sparse striatal expression (Fig. 2D).

## Conserved and divergent molecular architecture of the avian pallium and mammalian cortex

Having identified distinct axial gene expression patterns across the telencephalon, we next examined whether these patterns correspond to functional specialization within molecularly defined subregions. Using ST data across 25 telencephalic subregions, we analyzed gene expression profiles and observed marked transcriptional and functional divergence between the arcopallium and entopallium, two major pallial domains (Fig. 2E). This difference suggests that the arcopallium and entopallium subregions possess distinct cellular architectures and spatial patterns compared to other pallial regions. In line with this, NeuN staining revealed reduced neuronal density in both the arcopallium and entopallium relative to other pallial regions (fig. S9A).

Gene Ontology (GO) enrichment analysis of region-specific upregulated genes further distinguished arcopallium and entopallium, revealing unique functional signatures (Fig. S9, B and C). The entopallium showed additional enrichment for pathways related to cell migration and tissue remodeling (Fig. S9B), implying a neuroplasticity mechanism in thalamorecipient neurons (*37*) with parallels to mammalian cortical sensory integration circuits (*23*). These molecular features may support dynamic network reorganization and enhanced synaptic reconfiguration in response to visual input. Together, the results reveal substantial molecular heterogeneity within anatomically defined subdivisions, offering a basis for their specialized functions.

To assess cross-species relationships between avian and mammalian pallial structures, we performed comparative transcriptomic analysis using SAMap (*38*) to integrate our budgerigar ST data with published mouse pallial ST datasets (*39*). Within the avian pallium, the nidopallium intermediale (NI) showed greater transcriptomic similarity to the hyperpallium/medial regions (Fig. 2E). However, in cross-species comparison, the NI aligned more strongly with mouse cortical layer IV (L4) and exhibited a broader repertoire of functional pathways than hyperpallium/medial regions, indicating greater cellular diversity in NI (Fig. 2F, and fig. S9, D and E). The mesopallium showed transcriptomic similarity to multiple mouse brain areas (Fig. 2E), suggesting it comprises heterogeneous cell types with diverse developmental origins.

Strikingly, both the avian NIVL and arcopallium correlated more closely with the stratum lacunosum-moleculare (SLM) of the CA1 region in mammals than with lateral ventral cortex or amygdala (Fig. 2F), despite their similar topographical positions (Fig. 2G). This unexpected correspondence suggests that substantial morphological divergence occurred early in avian and mammalian evolution, indicating that regional homology cannot be inferred solely from conserved developmental positioning (*18*).

Although previous studies identified entopallium as a primary thalamorecipient zone functionally analogous to mammalian L4 (*32, 40*), our data showed minimal transcriptomic similarity between entopallium and L4 (Fig. 2F), suggesting they are unlikely to be direct homologs (*22, 23*). Instead, entopallium aligned more closely with cortical layer V (L5) (Fig. 2F). This correspondence may arise from conserved distributions of inhibitory neuron subtypes, particularly parvalbumin-positive (PV) interneurons (fig. S10A), and their shared transcriptional programs (*22, 41*), as well as similar oligodendrocyte distributions (fig. S10B), rather than from excitatory neuron identity.

## A systematic neuron taxonomy atlas of the adult budgerigar brain

To extend our analysis beyond spatial domains, we integrated snRNA-seq data from the adult budgerigar brain to resolve neuronal diversity at high resolution. We applied a hierarchical classification framework consisting of four levels: class, subclass, supertypes, and clusters (fig. S11; see also Materials and Methods). This classification scheme, used in previous single-cell profiling studies of the mouse (*42*) and chicken (*22*) brains, facilitates cross-species comparisons and enhances neuronal subtype resolution.

We applied this framework to snRNA-seq data across four major brain regions. In the telencephalon, 192,439 cells were assigned to 8 non-neuronal and 18 neuronal subclasses (10 excitatory, 8 inhibitory), the latter further resolved into 23 and 17 supertypes and 55 and 33 clusters, respectively (figs. S12 and S13). Subclass composition (fig. S14A), marker gene expression (fig. S14B), and batch integration quality (fig. S14C) were quantified, culminating in a structured taxonomy table (fig. S14D, and Data S3). In the midbrain, 37,373 cells yielded 8 non-neuronal and 14 neuronal subclasses (6 excitatory, 8 inhibitory), the latter grouped into 23 and 19 supertypes and 44 and 32 clusters, respectively (figs. S15, and S16). Subclass profiles and annotations were summarized in fig. S17 and Data S3. In the diencephalon, 38,652 cells were grouped into 8 non-neuronal populations, a distinct dopaminergic neuronal group, and 11 neuronal subclasses (6 excitatory, 5 inhibitory), subdivided into 12 and 15 supertypes and 36 and 31 clusters, respectively (figs. S18 and S19). Corresponding subclass summaries and marker validations are shown in fig. S20, A to D, and Data S3. In the cerebellum, 41,561 cells were assigned to 8 non-neuronal and 6 neuronal subclasses (3 excitatory, 3 inhibitory), resolved into 4 and 8 supertypes and 17 clusters each (fig. S21). Subclass identities, markers, and integration quality were assessed in fig. S22.

Together, these analyses yielded a region-resolved single-cell taxonomy, forming a comprehensive molecular atlas of the adult budgerigar brain (Fig. 3A).

**Fig. 3.**
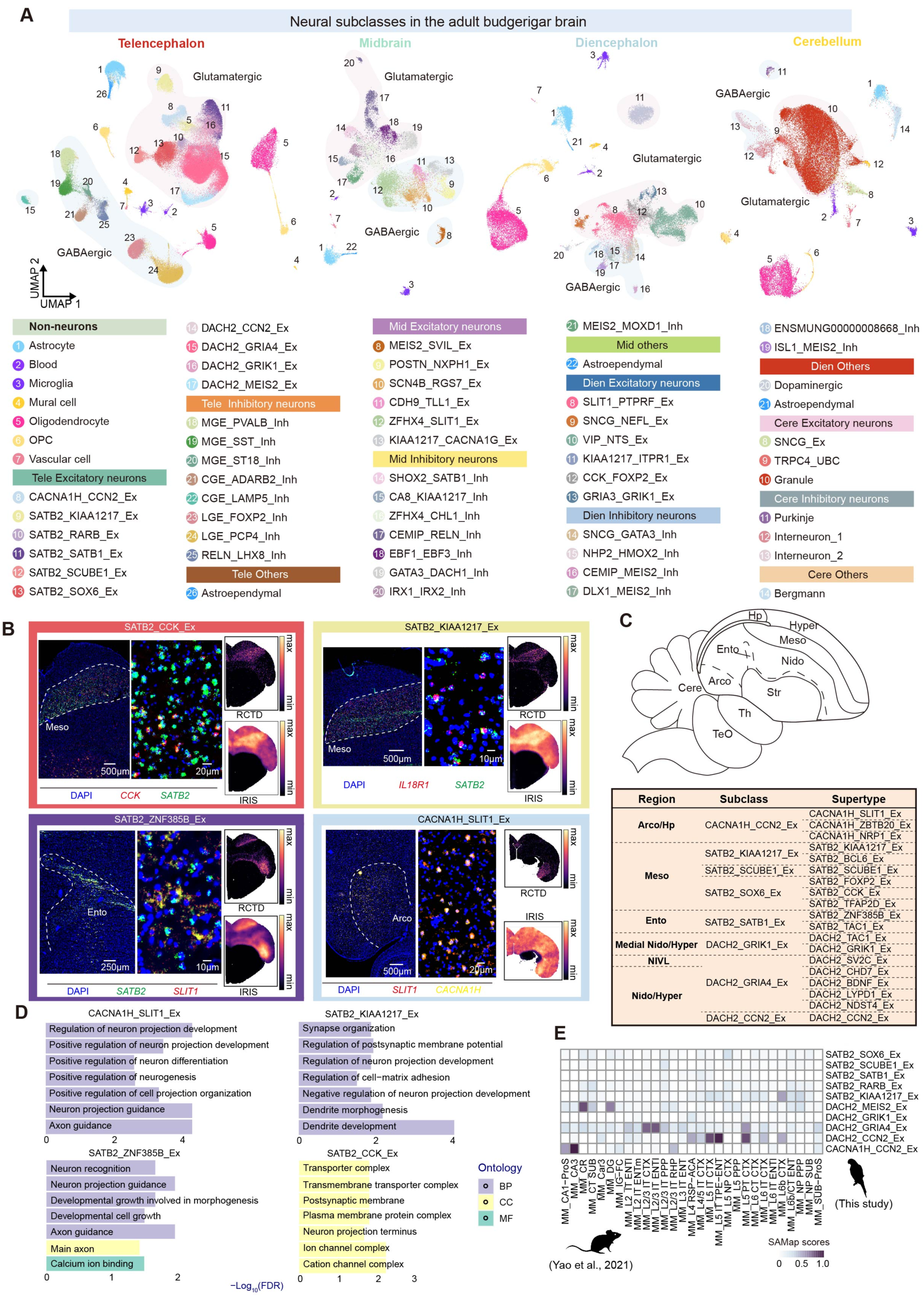
Cellular taxonomy and spatial organization of adult budgerigar brain neurons. (**A**) UMAP of major neuronal (Glutamatergic, GABAergic) and non-neuronal subclasses across four anatomical territories. Each point represents a single cell, colored by subclass identity. (**B**) Spatial localization of four excitatory neuron supertypes: SATB2_CCK_Ex, SATB2_KIAA1217_Ex, SATB2_ZNF385B_Ex, and CACNA1H_SLIT1_Ex. Spatial distributions were estimated using RCTD and IRIS deconvolution of Stereo-seq and validated by HCR RNA-FISH with subclass-specific markers. (**C**) Schematic showing excitatory neuronal subclasses and the spatial distribution of their corresponding supertypes. (**D**) Gene Ontology (GO) enrichment analysis of upregulated genes (log_2_FC > 0.5) for four representative supertypes. (**E**) Heatmap showing transcriptomic similarity of major excitatory neuron subtypes between budgerigar and mouse based on SAMap analysis.

## Spatial distribution and evolutionary relationships of neuronal cell types in the adult pallium

### Inhibitory neurons

With the neuron taxonomy atlas established, we examined the functional and evolutionary relationships of neuronal supertypes, beginning with inhibitory neurons in the adult budgerigar pallium. Prior studies have demonstrated that telencephalic inhibitory neurons are highly conserved across amniotes (*20, 22, 41, 43, 44*), originating from three distinct ganglionic eminences: lateral (LGE), caudal (CGE), and medial (MGE). Our comparative SAMap analysis of the mammalian cortex (mouse (*42*) and human (*45*)) and budgerigar telencephalon supports this view (fig. S23). Major GABAergic subclasses derived from the MGE, CGE, and LGE are evolutionarily conserved, but display divergent spatial distributions. For example, CGE-derived *LAMP5⁺* neurons in birds show transcriptional similarity to human *LAMP5⁺/LHX6⁺*chandelier cells (fig. S23C), yet are broadly dispersed across the pallium (fig. S24A). Although MGE-derived *PVALB*⁺ neurons in the avian brain show strong transcriptomic similarity to mammalian PV⁺ interneurons (fig. S23, A and B), their anatomical distribution is markedly different. In birds, these cells are predominantly found in the entopallium and interstitial hyperpallium apicale (IHA) (fig. S24, A and C), whereas in mammals, PV⁺ interneurons are enriched in cortical L4 and L5, forming a well-defined laminar pattern (*36*). This spatial divergence underscores functional analogies between the avian entopallium and mammalian L5 circuits, suggesting that inhibitory microcircuitry is conserved even in the absence of homologous excitatory architecture.

CGE-derived *ADARB2*⁺ inhibitory neurons (CGE_ADARB2_Inh) show strong conservation with mammalian CGE-derived interneurons (fig. S23C), despite lacking expression of *VIP*, a canonical CGE marker in mammals (*44*). Similarly, LGE-derived *PCP4⁺* neurons, confined to the striatum (fig. S24, A and B) and expressing either *DRD1* or *DRD2* receptors, align transcriptionally with human medium spiny neurons (MSNs) (fig. S23D), implying the early emergence of D1-MSN and D2-MSN subtypes in the amniotic ancestor (*41*). Although *PCP4* is typically enriched in mammalian telencephalic L4/5 and sparse in the striatum (*35*), its robust striatal expression in birds reflects regulatory divergence in the spatial deployment of conserved subtypes. These examples illustrate that evolutionarily homologous cell types can undergo significant divergence in spatial distribution and effector gene expression, while maintaining conserved core molecular identities.

### Excitatory neurons

Unlike inhibitory neurons, excitatory populations remain poorly characterized across amniotes, especially between avian and mammalian species in terms of transcriptional identity, spatial patterning, and developmental trajectories (*22, 23*). To address this, we integrated ST and snRNA-seq datasets using deconvolution algorithms [RCTD (*46*), IRIS (*47*)] and validated results via HCR RNA-FISH (fig. S4). This revealed distinct excitatory neuron distributions across major pallial domains including hyperpallium, nidopallium, mesopallium, entopallium, and arcopallium (Fig. 3B and fig. S25), and enabled mapping of supertype-specific spatial profiles (Fig. 3C), offering structural insights into avian neural circuitry.

To assess conservation across avian species, we transferred budgerigar supertype labels to zebra finch (*43*) and chicken single-cell/nucleus RNA-seq datasets (*22*) (fig. S26, A to D). Cross-species integration using a gene-specific index (GSI)-based mapping (see Materials and Methods) identified conserved transcriptional signatures. This analysis yielded a unified classification of telencephalic excitatory neurons at the supertype level, defining eight major subclasses (fig. S26E). These subclasses co-express *DACH2, SATB2*, and *CACNA1H*, but are distinguished by subclass-specific markers (fig. S26, A to E). Notably, the SATB2_SATB1_Ex subclass, which includes SATB2_TAC1_Ex and SATB2_ZNF385B_Ex, uniquely co-expresses multiple defining factors.

### Conserved glutamatergic cell types in the hippocampus and amygdala

Recent studies have highlighted structural parallels between the avian hippocampus and arcopallium (*17, 22*). We identified three CACNA1H_CCN2_Ex supertypes, namely CACNA1H_SLIT1_Ex, CACNA1H_ZBTB20_Ex and CACNA1H_NRP1_Ex, which exhibit medial–lateral gradients in both regions and recapitulate the Ex_CACNA1H subclasses previously described in the chicken (*22*) (fig. S26E and fig. S27A). This topographic organization implies that these supertypes arise from shared developmental trajectories, with transcriptomic diversification occurring prior to their anatomical compartmentalization. Cross-amniote comparisons revealed that CACNA1H_CCN2_Ex types are conserved homologs of hippocampal cell types (figs. S28 and S29). In particular, the *ZBTB20*⁺ lineage shows triadic conservation across chicken (Ex_CACNA1H_CPA6/PROX1 (*22*), EXC_GLU-6a (*41*)), turtle (DMC), and mammals (deep CA neurons), and enriched for neuron projection functions (fig. S27B), pointing to a deeply conserved pyramidal neuron lineage. The CACNA1H_SLIT1_Ex supertype, primarily localized to the arcopallium and secondarily to the hippocampus, is enriched for neurodevelopmental functions, including axon guidance and projection development (Fig. 3D). It aligns with zebra finch RA excitatory neurons, a key sensorimotor population (*43*) (fig. S26E), and is homologous to chicken EXC_GLU-6b (*41*) and Ex_CACNA1H_KIT (*22*). The latter shows molecular similarity to murine CA1 pyramidal neurons, despite the absence of direct mouse homologs in our dataset (fig. S28A). While this population correlates weakly with human CA1–3 neurons, it resembles CA4 populations (fig. S29B), suggesting evolutionary divergence from a medial pallium precursor with conserved functional modules.

The avian excitatory subtype CACNA1H_NRP1_Ex shows strong cross-species homology to Ex_CACNA1H_LHX9 in the chicken (fig. S26E), a population previously reported to exhibit transcriptional similarity to *Slc17a7⁺* neurons in the murine amygdala, CA3 pyramidal cells, the lizard medial cortex (MC), and amygdalar clusters (*22*). Our alignment further reveals conserved correspondence with human CA4 and amygdala excitatory neurons (fig. S29B), underscoring CACNA1H_NRP1_Ex as an amygdalar pyramidal-like lineage. These results position this population as a core arcopallial subtype with deep evolutionary ties to amygdalar circuits across amniotes.

### Evolutionary and cellular complexity of the mesopallium

The evolutionary origin of the mesopallium remains under debate. While several studies support its homology to the mammalian claustrum (*22, 48*), the potential relationship to IT neurons is unresolved (*23, 33, 49*). In agreement with previous studies in chickens, we identified three major excitatory subclasses, namely SATB2_KIAA1217_Ex, SATB2_SCUBE1_Ex, and SATB2_SOX6_Ex, which correspond to the chicken subclasses EXC_GLU_4, EXC_GLU_5, and EXC_GLU_7 (*41*) (fig. S26E). Although all express *SATB2*, a conserved cortical transcription factor, they exhibit pronounced molecular heterogeneity (Fig. 3A), indicating functional specialization. SATB2_SCUBE1_Ex (homologous to chicken Ex_SATB2_ZNF385B (*22*) and EXC_GLU_5 (*41*)) displays strong evolutionary conservation, aligning with mammalian claustral neurons (*22*) and reptilian pallial thickening (PT) neurons (fig. S30A). This supports its origin in the lateral pallium and suggests retention of a *SCUBE1*⁺ *SATB2*⁺ lineage across amniotes, diversified for associative circuitry. SATB2_KIAA1217_Ex (including SATB2_BCL6_Ex) corresponds to chicken EXC_GLU-7 (*41*) and Ex_KIAA1217 (*22*), and exhibits notable UMAP heterogeneity (Fig. 3A and fig. S12F). It is enriched for genes involved in dendritic growth, cell adhesion, and projection development (Fig. 3D). Although it resembles murine layer VIb (L6b) neurons (*22, 41*) (Fig. 3E), it lacks their characteristic long-range projections (*40*). Cross-species comparisons indicate stronger similarity to reptilian dorsal cortex (DC) and PT neurons (fig. S30, A and B), as well as human deep-layer near-projecting neurons (fig. S29, A and B). These data suggest SATB2_KIAA1217_Ex derives from an ancestral L6b-like lineage that underwent functional divergence. Within the *SATB2*⁺ group, SATB2_SOX6_Ex is distinguished by limited involvement in complex biological processes (Fig. 3D). This subclass consists of SATB2_CCK_Ex, SATB2_FOXP2_Ex, and SATB2_TFAP2D_Ex, which correspond to chicken EXC_GLU-4 (*41*) (fig. S26E) and partially to mammalian IT and CT neurons (fig. S28A). In addition, we observed that the neuronal population displayed higher similarity to entorhinal IT neurons, as well as to reptilian posterior lateral cortex (pLC) and anterior DVR (aDVR) (annotated as amDVR in the Zaremba *et al.* dataset) (*22*) (fig. S30, A and B). These findings suggest that SATB2_SOX6_Ex likely originated from the entorhinal IT neurons but evolved gene regulatory modules convergent with neocortical IT neurons (*22*).

### Entopallium, IHA, and Field L

The SATB2_ZNF385B_Ex supertype, part of the SATB2_SATB1_Ex subclass, is distributed across both the entopallium and IHA (Fig. 3B), providing cellular-level support for proposed circuit homologies (*40*) and clarifying regional relationships (*16, 17*). Functionally, this population is enriched for axon guidance and projection genes (Fig. 3D), resembling mammalian L4 neurons (*37, 40*), though lacking direct molecular homology (*22*). Its co-expression of *DACH2* and *SATB2* mirrors chicken Ex_DACH2_RORB (*22*) (fig. S26E), which aligns with piriform (PIR) input neurons (Fig. 3B). These data suggest that SATB2_ZNF385B_Ex arose from ventral pallial PIR input neurons and later acquired L4-like features, reflecting functional reprogramming in birds.

### Shared cell types in hyperpallium and nidopallium

The avian DVR and Wulst (including the hyperpallium apicale (HA), intercalated hyperpallium apicale (IHA), hyperpallium intercalatum (HI), and hyperpallium densocellulare (HD) exhibit marked dorsoventral symmetry across multiple scales, including neural circuitry, gene expression patterns, and cell-type composition (*17, 22, 40*). To understand the molecular logic underlying this symmetry, we focused on excitatory neuron populations that span the hyperpallium and nidopallium, the two principal domains defining the dorsoventral axis. Spatial mapping with RCTD, validated by HCR RNA-FISH, identified three *DACH2*⁺ excitatory subclasses, DACH2_CCN2_Ex, DACH2_GRIA4_Ex, and DACH2_GRIK1_Ex, with distinct, symmetric localization patterns along this axis (figs. S25 and S31). Among these, DACH2_CCN2_Ex stood out for its strong alignment with excitatory clusters in the chicken brain, including EXC_GLU_2 (*41*) and Ex_DACH2_ZMAT4 (*22*) (fig. S26E), and for its transcriptomic similarity to mammalian IT neurons located in cortical L4 and L5 (Fig. 3E and fig. S28).

Interestingly, both our data and Zaremba *et al.* dataset (*22*) suggest that DACH2_CCN2_Ex also partially overlaps with the molecular signature of L5 pyramidal tract (PT) neurons. Cross-species comparisons mapped DACH2_CCN2_Ex to aDC and pDC of turtles (fig. S30, A to B), PT-like neurons in the mammalian posterior dorsal cortex (fig. S28), and to pyramidal neurons in the human CA1 to CA3 regions (fig. S29, A to B). Notably, aDC and pDC in turtles correspond to IT and PT neuron types in the mouse, respectively (fig. S30C), suggesting that IT and PT modules may have been developmentally integrated during sauropsid evolution.

Together, these findings identify DACH2_CCN2_Ex as a conserved pyramidal-like population that likely supports both local hyperpallial processing (*50*) and long-range projection pathways (*40, 51*), providing a cellular correlate for the dorsoventral symmetry observed in the avian pallium. By contrast, DACH2_GRIA4_Ex that presents in both nidopallial and hyperpallial regions, shared transcriptomic profiles with upper-layer IT neurons of the mammalian entorhinal cortex (Fig. 3E) and lateral cortex (LC) domains in turtles (fig. S30, A and B). This subclass consists of six distinct supertypes (fig. S12F), indicating substantial internal heterogeneity. A third major subclass, DACH2_BDNF_Ex, defined by robust *BDNF* expression and corresponding to EXC_GLU-1b-2 (*41*)and Ex_DACH2_NR4A3 (*22*) (fig. S26E), aligned closely with mammalian L2/3 IT neurons (fig. S28) but paradoxically mapped to ventral DVR clusters in turtles (fig. S30B). Single-cell studies in mice have shown that L2/3 IT neurons from entorhinal, piriform, and neocortical regions cluster more tightly with one another than with other IT subtypes (*42*), suggesting that DACH2_BDNF_Ex and the other two supertypes (DACH2_CDH7_Ex, DACH2_NDST4_Ex) may represent a conserved ventrally originated IT-like lineage in amniotes. Additional supertypes, such as DACH2_LYPD1_Ex, DACH2_ZNF381_Ex, and DACH2_SV2C_Ex, were predominantly localized to the lateral nidopallium, including NIVL, NLc, and NCL (fig. S25). These populations correlated with turtle LC excitatory clusters and displayed specialized functional signatures (fig. S30B). Notably, DACH2_ZNF381_Ex was enriched in anterior NLc and showed high conservation with HVC_GLUT-2 neurons in zebra finch (*43*) (fig. S26E), suggesting a role in auditory circuit formation. Both DACH2_LYPD1_Ex and DACH2_ZNF381_Ex were situated near the arcopallium and partially overlapped with amygdala-associated excitatory types, as confirmed in chicken datasets (*41*) (fig. S29B). These findings suggest that amygdala-like neurons extend into defined nidopallial territories, supporting partial but not complete homology between the arcopallium and mammalian amygdala (*41*).

Through fine-scale spatial transcriptomics and cross-species integration of single-cell datasets, we uncovered a conserved, dorsoventrally symmetric architecture in the adult avian pallium. Some subtypes displayed clear transcriptional and anatomical homology to mammalian cortical and hippocampal neurons; others aligned more closely with turtle pallial structures, highlighting both divergence and deep conservation among amniotes. These results raise two central questions: (i) What molecular cues and lineage dynamics establish dorsoventral symmetry in the mature avian brain? (ii) How do embryonic programs orchestrate this spatial patterning over developmental time? Addressing these will provide critical insights into the developmental and evolutionary logic of pallial organization.

## Multimodal analysis redefines pallial organizational models underlying dorsoventral symmetry

The organization of the avian telencephalon has long been controversial, with three major hypotheses proposed. The “distinction model” (including tripartite, tetrapartite, and hexapartite versions) asserts that the Wulst lies dorsally to a vestigial ventricle and is structurally distinct from ventral pallial regions, including the mesopallium, nidopallium, and arcopallium, collectively termed the DVR (*48, 52*) (Fig. 4A). This model, however, does not reconcile the prominent dorsoventral symmetry in cell-type architecture. In contrast, the “continuum model” envisions a continuous cellular gradient that wraps around the embryonic ventricle, which fuses before hatching and creates an apparent dorsal–ventral separation (*16, 17*) (Fig. 4A). Yet, recent lineage tracing by Zaremba *et al.* indicates that the hyperpallium and nidopallium arise from distinct progenitor pools, suggesting convergent evolution rather than developmental continuity (*22*) (Fig. 4A).

**Fig. 4.**
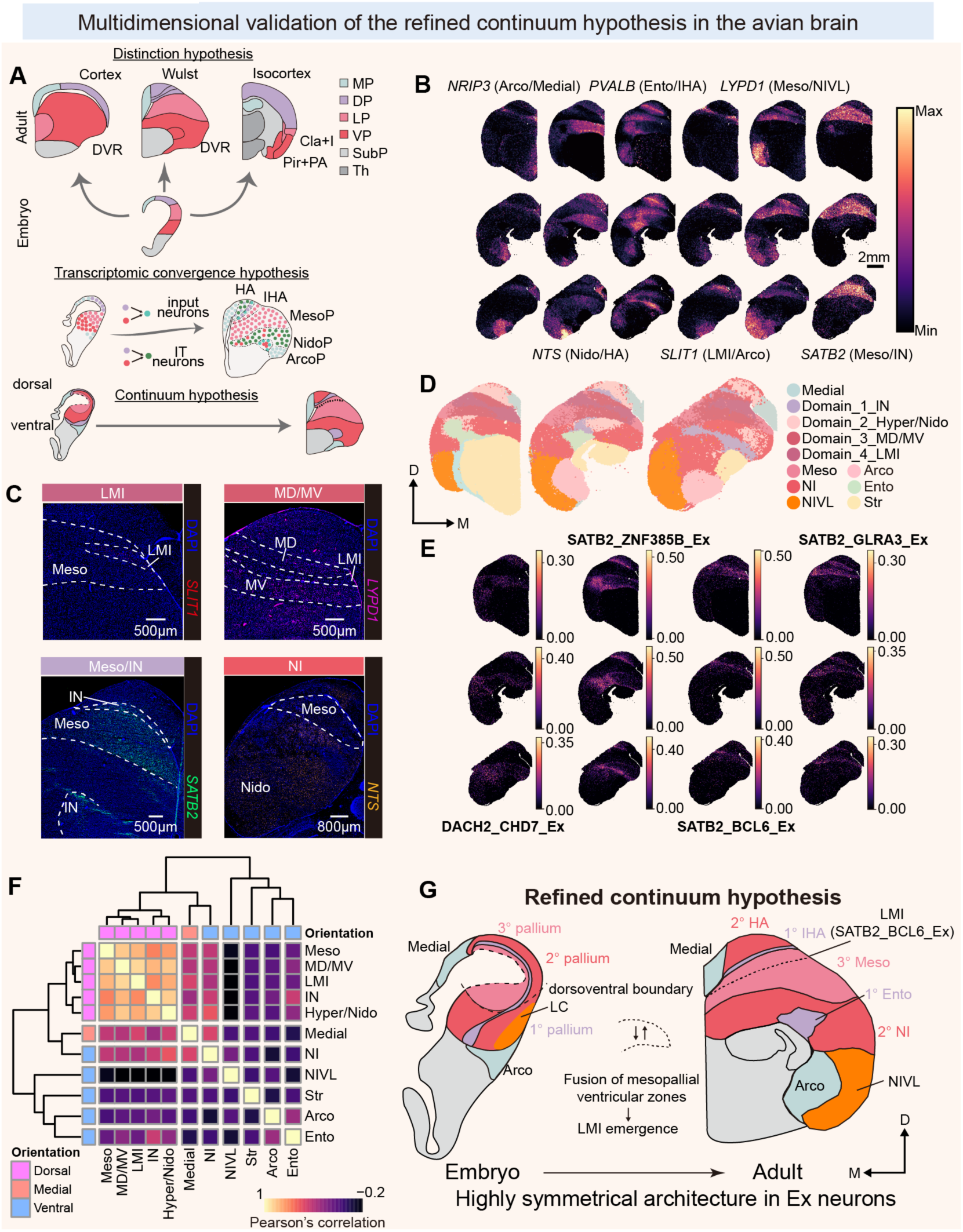
Multiscale anatomical and molecular characterization of dorsoventral symmetry in the avian pallium. (**A**) Schematic comparison of three proposed models of pallial organization: classical discrete-domain model (top), transcriptional convergence model (middle), and continuum model (bottom). (**B**) Spatial transcriptomic maps showing dorsoventrally symmetrical expression of telencephalic marker genes (*NRIP3, PVALB, LYPD1, NTS, SLIT1, SATB2*). (**C**) HCR RNA-FISH validation of gene expression patterns for *SATB2, SLIT1, NTS,* and *LYPD1* across dorsal (hyperpallium apicale, HA; interstitial HA, IHA) and ventral (nidopallium, NidoP) pallial regions. (**D**) Spatial Leiden clustering of transcriptomic data of three domains: intercalated pallium (IN), hyperpallium–nidopallium (Hyper-Nido), and dorsal/ventral mesopallium (MD/MV). (**E**) Spatial distribution of representative excitatory neuron supertypes SATB2_BCL6_Ex and SATB2_GLRA3_Ex. (**F**) Pearson correlation of global gene expression profiles across brain regions. (**G**) Schematic of the refined continuum model illustrating the lamina mesopallialis intermedia (LMI) as the central boundary between dorsal and ventral mesopallial domains, with terminology based on Jarvis *et al.* (*16, 17*).

To evaluate these models, we conducted analysis from six perspectives: (1) gene expression, (2) HCR RNA-FISH, (3) spatial transcriptomic clustering, (4) cell-type distribution, (5) functional enrichment, and (6) developmental lineage. Marker gene mapping (e.g., *NRIP3, NTS, PVALB, SLIT1, LYPD1,* and *SATB2*) revealed specific, regionally restricted expression in at least two brain domains (Figs. 2, B to D, and 4, B). In the mesopallium, *SLIT1* was highly enriched in the LMI, while *LYPD1*, the mammalian homolog of which is *Lynx1* that stabilizes mature cortical circuits, was confined to the dorsal and ventral mesopallium and absent from LMI. *SATB2* was broadly expressed across mesopallial regions. HCR RNA-FISH confirmed the spatial transcriptomic findings (Fig. 4C).

To assess dorsoventral symmetry relative to LMI, we reanalyzed spatial transcriptomic data using Spatial Leiden clustering (fig. S32). The IHA closely resembled Field L in transcriptional profile but diverged from the entopallium (Fig. 2C). Hyperpallium and nidopallium showed overlapping clusters, whereas NIVL displayed a distinct molecular signature. The dorsal and ventral mesopallium (MD and MV) remained transcriptionally indistinguishable, consistent with prior findings (*17*). In contrast, LMI exhibited a unique transcriptomic identity (Fig. 4D).

Analysis of excitatory neurons revealed region-specific enrichment of SATB2_ZNF385B in entopallium, IHA, and Field L. CACNA1H_CCN2 displayed dorsoventral symmetry across the hippocampus and arcopallium. Hyperpallium and nidopallium harbored symmetric distributions of excitatory subtypes. SATB2_GLRA3_Ex, linked to postsynaptic functions (fig. S33A), was symmetrically distributed in MD and MV (Fig. 4E). In contrast, SATB2_BCL6_Ex, which defined the LMI (Fig. 4E), was enriched in pathways related to presynaptic and projection terminal processes (fig. S33B). These patterns suggest that LMI arises developmentally via synaptic coupling between MD and MV, potentially facilitating ventricle fusion.

While convergent evolution has been proposed to explain similarities between dorsal and ventral pallial regions (*22*), this explanation does not account for the striking dorsoventral symmetry organized around the LMI. Pearson correlation of gene expression revealed that the hyperpallium exhibited high correlation with all regions of the mesopallium, whereas the highly symmetric regions of excitatory neurons did not show identical overall gene expression patterns (Fig. 4F). This suggests that the dorsal-ventral continuum was primarily applicable to excitatory neurons, while the overall correlations indicate that the tripartite division (DP, MP, VP) of the avian brain may provide a more accurate framework for understanding dorsal-ventral organization. Integrating these results, we propose an updated continuum model, identifying the LMI as a distinct pallial layer defined by *SLIT1* enrichment, *LYPD1* exclusion, and the selective presence of SATB2_BCL6_Ex neurons. Moreover, NIVL excitatory neurons (e.g., DACH2_SV2C_Ex) lacked overlap with hyperpallial subtypes (Fig. 4G), underscoring regional discontinuities that reshape current models of avian pallial evolution.

## Spatial distribution and developmental origin of telencephalic neurons

To trace the developmental origins of telencephalic excitatory neurons, we classified cells from three stages (E14, P1, and juvenile) into developmental (Dev) subtypes (fig. S34, A and B; see also Materials and Methods). We then inferred terminal fates by mapping adult supertypes onto developmental trajectories via label transfer (*53*) and aligning them with pseudotime (Fig. 5A). Mapping marker genes from adult region-specific supertypes (fig. S34, C and D, and Data S4) allowed identification of 15 distinct Dev subtypes. These analyses indicated that telencephalic excitatory neurons derive from a shared progenitor population (Progenitor_Ex), homologous to mammalian neocortical intermediate progenitors (fig. S35), and subsequently diversify through distinct trajectories (Fig. 5B). Pseudotime analysis supported these lineage assignments (Fig. 5C). Label transfer further revealed that SATB2_KIAA1217_Ex, DACH2_MEIS2_Ex, CACNA1H_SLIT1_Ex, and SATB2_SCUBE1_Ex neurons complete maturation during early development (Fig. 5D). Notably, SATB2_SCUBE1_Ex, a claustrum-homologous subtype, arose independently of Progenitor_Ex (Fig. 5B and C), indicating a distinct lineage and asynchronous maturation relative to other excitatory neurons. Interestingly, nidopallium and hyperpallium did not exhibit distinct developmental origins. Instead, two lineages, Nido/Hyper_GRIK1/CCN2_Ex and Nido/Hyper_GRIA4_Ex, were shared in both regions, indicating common ancestry and arguing against convergent evolution of dorsal–ventral transcriptional modules (*22*).

**Fig. 5.**
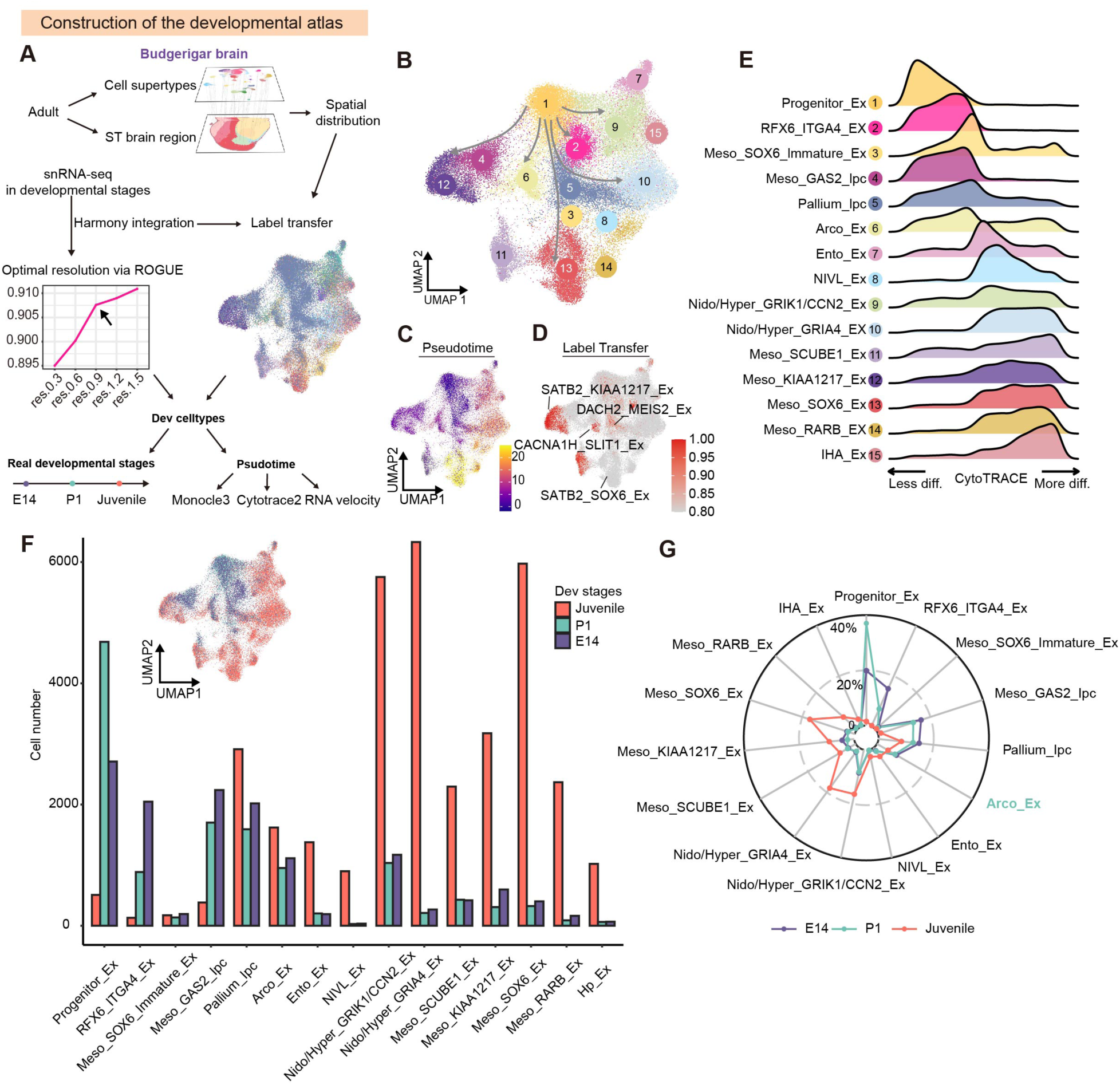
Developmental trajectory mapping of telencephalic excitatory neurons in the budgerigar. (**A**) Workflow for building the developmental atlas using snRNA-seq, label transfer, and spatial integration. (**B**) UMAP of integrated excitatory neurons across stages reveals distinct developmental trajectories. (**C**) Pseudotime inference using Monocle3, with higher scores indicating greater maturity. (**D**) Label transfer from adult supertypes predicts developmental states of excitatory cells. (**E**) CytoTRACE2 analysis ranks cells by differentiation potential. (**F)** Cell type abundance across E14, P1, and juvenile brains shows progressive emergence of subtype identities. **(G**) Radar plot quantifying subtype proportions at different developmental timepoints.

Although both were classified as mesopallial neurons, distinct mesopallial subclasses displayed divergent lineages (fig. S36, A and B). Integrated developmental and evolutionary analyses revealed ontogenetic divergence underlying mesopallial heterogeneity. Meso_SOX6_Ex neurons originated from pallial intermediate precursor cells (Pallium_Ipc) and Meso_SOX6_Immature_Ex progenitors, paralleling the conserved neurogenesis programs of immature mammalian neocortical neurons. In contrast, Meso_KIAA1217_Ex neurons arose from the avian-specific Meso_GAS2_Ipc lineage. The Meso_KIAA1217_Ex lineage exemplifies an evolutionary adaptation: while its progenitors (Meso_GAS2_Ipc) transcriptionally resemble mammalian upper-layer corticopontine neurons (UL_CPN), the mature *KIAA1217^+^* neurons exhibit L6b-like features. This phenotypic transition suggests that during avian evolution, the Meso_GAS2_Ipc lineage incorporated IT-associated transcriptional programs, enabling its L6b-homologous descendants to co-express upper-layer markers (e.g., *SATB2*) while retaining deep-layer projection characteristics. The apparent absence of a mammalian counterpart to Meso_SOX6_Ex argues against a direct homology with neocortical IT neurons (*49*) and instead points to a lineage that may have emerged independently within the archosaurian mesopallium. RNA velocity analysis revealed distinct trajectories and gene upregulation patterns between Meso_SOX6_Ex and Meso_KIAA1217_Ex (fig. S36, C and D). Although differentiation paths diverged, both lineages displayed consistent downregulation of *POSTN* and upregulation of *SATB2* (fig. S36, E and F), suggesting convergence of regulatory mechanisms despite lineage divergence.

In the mammalian cortex, excitatory neurogenesis is temporally graded and proceeds in a layer-specific sequence, giving rise to birthdate-dependent neuronal diversity (*54*). Unexpectedly, the avian pallium also undergoes asynchronous neuronal maturation, despite lacking the laminar organization characteristic of the mammalian cortex (*23*). Our analysis identified CACNA1H_SLIT1_Ex, an excitatory subtype enriched in the arcopallium, as one of the earliest-maturing populations. To assess whether the Arco_Ex follows an accelerated developmental trajectory, we integrated pseudotime inference with actual stage annotations. Differentiation scores computed by Cytotrace2 (*55*) revealed that both stem-like and terminal populations including Progenitor_Ex, RFX6_ITGA4_Ex (Cajal–Retzius cells, fig. S35), Meso_SOX6_Immature_Ex, Meso_GAS2_Ipc, Pallium_Ipc, and Arco_Ex, exhibited low differentiation levels (Fig. 5E), consistent with a compressed maturation timeline. Comparisons with murine datasets further demonstrated that Arco_Ex had the lowest transcriptional correlation with migrating neuronal populations (fig. S35), reflecting reduced neurodevelopmental gene expression and supporting its early maturation. Quantification of cell-type composition across developmental stages revealed a greater proportion of embryonic and neonatal cells within arcopallium compared to other pallial regions (Fig. 5, F to G), reinforcing the notion of accelerated maturation, which is in agreement with a recent study (*23*).

Together, these findings define core principles of telencephalic development: (i) excitatory neurons follow multiple, distinct neurogenic trajectories; (ii) a dorsoventral continuum of identities emerges during development; and (iii) neuronal populations exhibit marked asynchrony in maturation.

## Birthdate dependent inside-out development in the avian optic tectum

A hallmark of mammalian neocortical development is its birthdate-dependent, inside-out laminar organization (*54, 56*). Whether the avian optic tectum (TeO), which exhibits comparable lamination, follows a similar pattern remains unresolved. The TeO consists of 15 layers (*57*) (Fig. 6A), with neuronal enrichment predominantly in layers 2–13, corresponding to the stratum griseum superficiale (SGF) and stratum griseum centrale (SGC), which mediate early visual processing (*58*). RCTD-based deconvolution of ST data resolved layer-specific enrichment of excitatory neurons, with pronounced localization in the SGC and SGF (*57*) (Fig. 6B and fig. S38A).

**Fig. 6.**
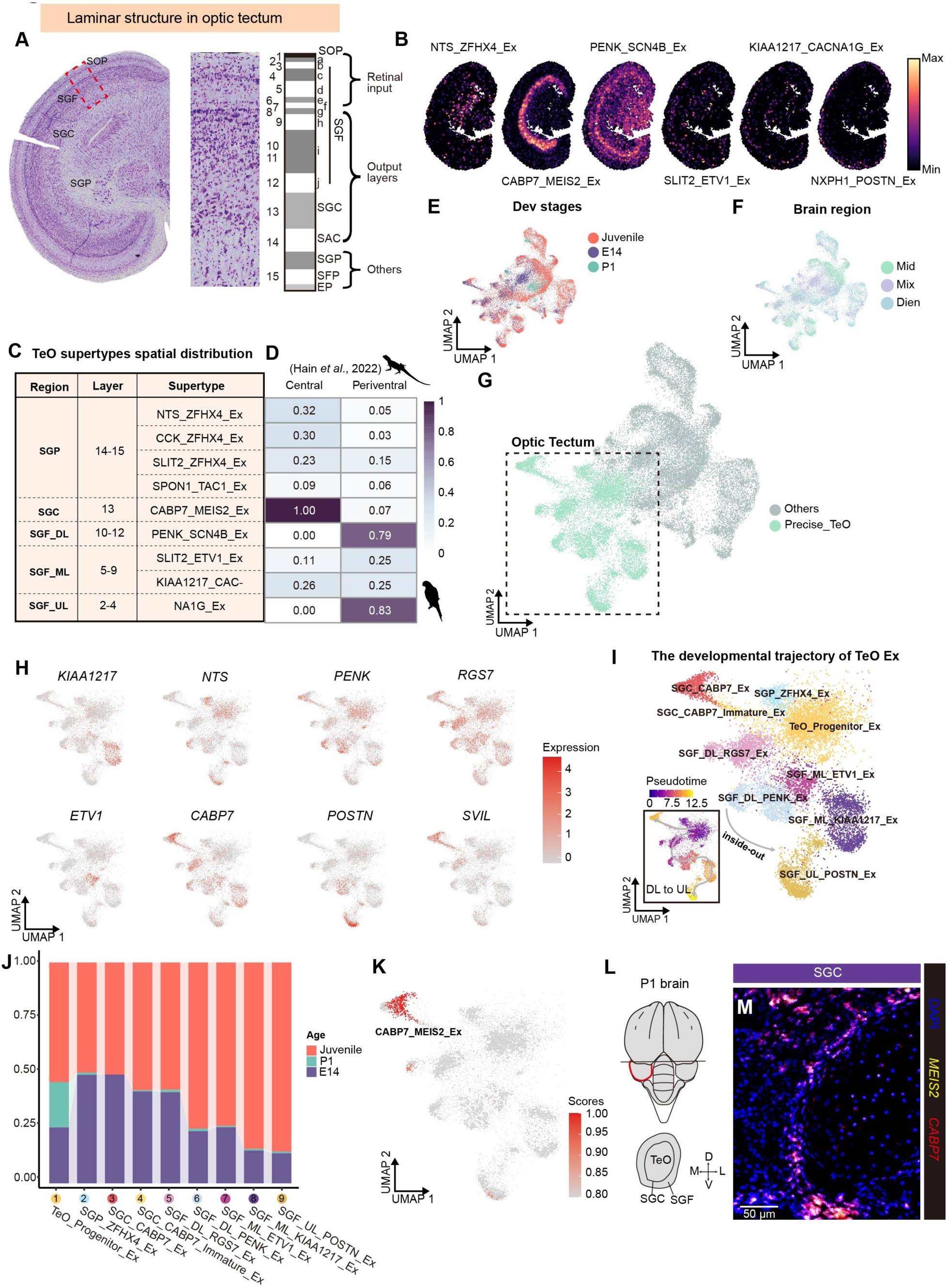
Birthdate-dependent inside-out manner of excitatory neurons in the budgerigar optic tectum. (**A**) Nissl staining of the budgerigar TeO. TeO, optic tectum. (**B**) Spatial distribution patterns of adult TeO excitatory supertypes across laminar zones. (**C**) Anatomical layer assignments of each supertype. (**D**) Cross-species comparison of laminar organization between the budgerigar and the lizard (*59*). (**E, F**) Integration of developmental datasets colored by stage (**E**) and region (**F**). (**G**) UMAP showing TeO-specific excitatory clusters. (**H**) Marker gene expression across TeO supertypes. (**I**) Pseudotime analysis showing an inside-out differentiation trajectory from deep (SGP) to upper (SGF_UL) layers. (**J**) Proportions of each excitatory subtype across developmental stages. (**K**) Predicted developmental state of CABP7_MEIS2_Ex cells. (**L**) Schematic of P1 TeO sectioning. (**M**) HCR RNA-FISH detecting *CABP7⁺MEIS2⁺* neuronal supertypes in the SGC at P1, with section levels corresponding to (**L**).

Among these, the CABP7_MEIS2_Ex supertype, part of the MEIS2_SVIL_Ex subclass, was spatially confined to the SGC. This pattern was validated by HCR RNA-FISH (Fig. 6C and figs. S38, B to E) and mirrored by a homologous domain in the lizard midbrain (*59*), suggesting deep conservation across sauropsids (Fig. 6D). In contrast, SGF supertypes showed graded distributions across laminae: PENK_SCN4B_Ex was enriched in deep SGF layers (DL; layers 10–12), with strong transcriptional similarity to the lizard periventricular region. SLIT2_ETV1_Ex and KIAA1217_CACNA1G_Ex localized to the middle SGF layers (ML; layers 5–9), while NXPH1_POSTN_Ex was enriched in upper SGF layers (UL; layers 2–4), showing high conservation with lizard homologs (Fig. 6D). These laminar patterns underscore conserved structural and cellular features in the optic tectum of sauropsids.

To chart the developmental trajectory of TeO excitatory neurons, we integrated embryonic-to-juvenile snRNA-seq data and applied label transfer from adult references to resolve TeO-specific subsets, despite partial diencephalic contamination in embryonic samples (Fig. 6, E to G, and fig. S39; see also Materials and Methods). To define developmental trajectories, we examined expression of layer-specific markers across TeO strata: SGP (*NTS*), SGC (*CABP7, SVIL*), and SGF (*RGS7, PENK, ETV1, KIAA1217,* and *POSTN*) (Fig. 6H). Despite originating from a shared progenitor pool, these regions diverged into distinct trajectories (Fig. 6I). Pseudotime analysis revealed a progressive maturation gradient in SGF (Fig. 6I), proceeding from deep to superficial layers, whereas SGC followed a more uniform and rapid developmental course.

Developmental-stage composition analysis showed a strikingly low proportion of SGF cell types at E14, in contrast to the SGC_CABP7_Ex population, which comprised approximately 50% of total cells, suggesting that SGC may have reached maturation by the E14 developmental time point (*60*) (Fig. 6J). Label transfer corroborated this conclusion, identifying SGC neurons as the earliest to mature within the TeO (Fig. 6K). This early differentiation is reflected in the adult TeO, where these neurons are enriched for genes involved in synaptic transmission and integrative processing (fig. S40), implicating them in early sensory computation (*60*). HCR RNA-FISH and P1-stage transcriptomic data further validated this interpretation (Fig. 6, L and M).

In summary, our analysis suggests that the SGC, located deeper than the SGF, undergoes earlier maturation, while the SGF displays a deep-to-superficial differentiation gradient. These findings reveal that TeO excitatory neurons develop in a birthdate-dependent inside-out manner, paralleling the laminar assembly of the mammalian neocortex.

## Molecular identity and conservation of adult neural stem cells in avian species

Mapping adult telencephalic excitatory supertypes onto developmental data revealed that DACH2_MEIS2_Ex closely resembled Pallium_Ipc (Fig. 5D). To further explore this relationship, we calculated DEGs for DACH2_MEIS2_Ex in adults and Pallium_Ipc during embryogenesis. These DEG sets exhibited substantial overlap (Fig. 7A), suggesting a shared cellular identity. Label transfer from developmental to adult excitatory neurons confirmed this connection: the majority of DACH2_MEIS2_Ex cells were assigned to Pallium_Ipc with high confidence scores (Fig. 7B). This finding implies that the adult avian brain may retain neural stem cells (NSCs) (*8, 9, 13, 61*) originating from early developmental stages (*62*). GO analysis showed that DACH2_MEIS2_Ex is significantly enriched for neurogenesis-associated and mitochondrial gene programs (Fig. 7C), underscoring its potential role in adult neurogenesis. To further assess developmental continuity, we used MetaNeighbor (*63*) to quantify transcriptional similarity between adult and embryonic populations. Most developmental types showed strong correspondence with their adult counterparts (Fig. 7D), with DACH2_MEIS2_Ex again exhibiting high similarity to Pallium_Ipc, reinforcing the inference of lineage continuity.

**Fig. 7.**
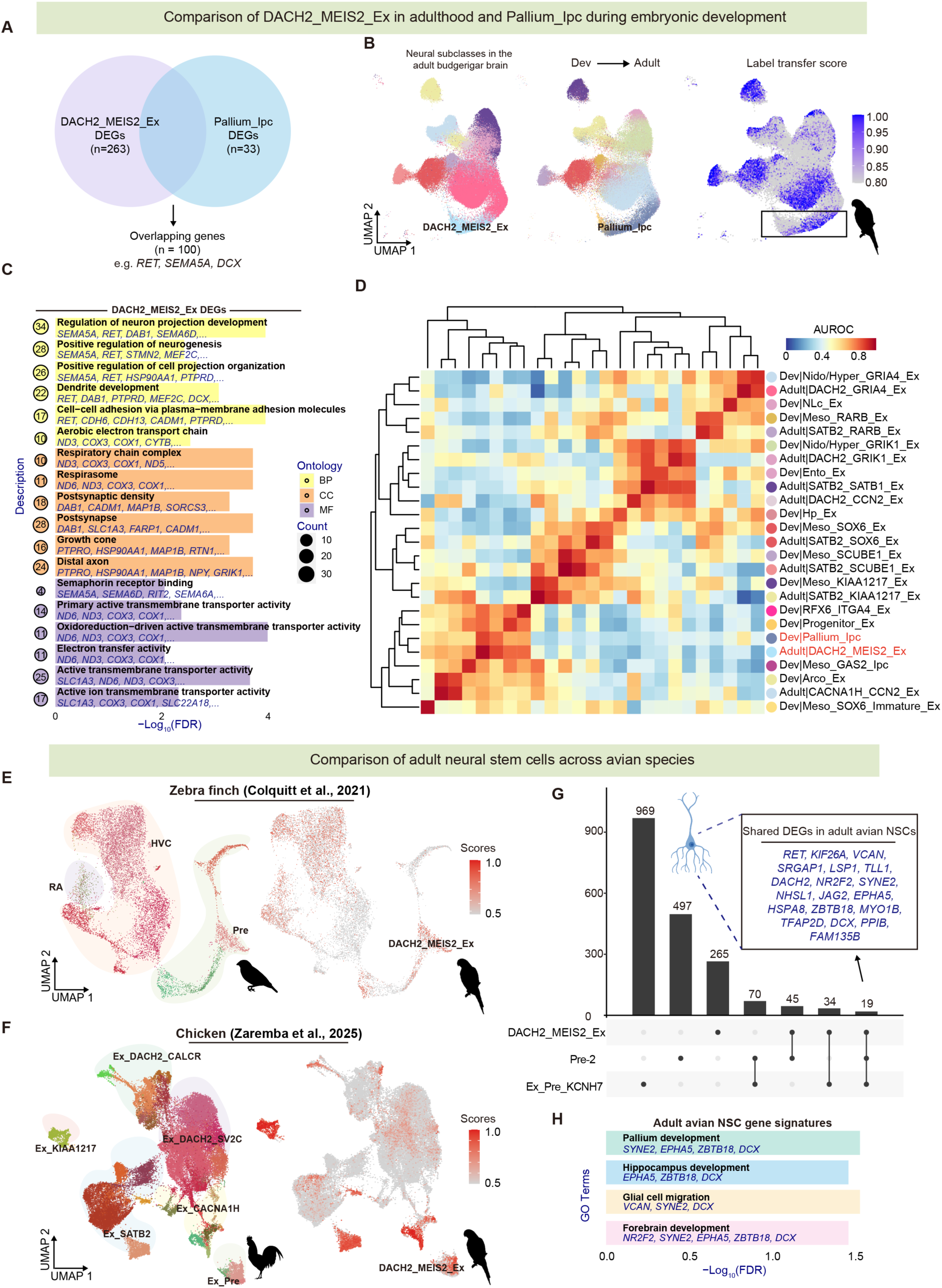
DACH2_MEIS2_Ex marks a putative adult neural stem cell population in the avian pallium. (**A**) Overlap of differentially expressed genes (DEGs) between adult DACH2_MEIS2_Ex and embryonic Pallium_Ipc populations. (**B**) Label transfer mapping DACH2_MEIS2_Ex cells onto embryonic populations. (**C**) GO enrichment analysis of DACH2_MEIS2_Ex DEGs showing enrichment for stemness– and migration-related functions. (**D)** MetaNeighbor analysis displaying transcriptional similarity between DACH2_MEIS2_Ex and embryonic pallial IPC-like cells. (**E, F**) Label transfer mapping DACH2_MEIS2_Ex-like cells in the adult zebra finch (**E**) and chicken (**F**) pallium. (**G**) UpSet plot showing conserved expression of adult neural stem cell (NSC) genes across avian species, including canonical markers (e.g. *RET, DCX, NR2F2, EPHA5*). (**H**) GO enrichment analysis of conserved adult NSC DEGs showing associations with pallial and hippocampal development, glial migration, and cytoskeletal organization.

To assess molecular conservation across species, we compared adult cell types in budgerigar, zebra finch, and chicken (fig. S28). DACH2_MEIS2_Ex corresponded to Pre-2 in zebra finch (*43*) and to Ex_Pre_KCNH7 in chicken (*22*) (Fig. 7, E and F), indicating conservation of transcriptomic identity across avian lineages. We next identified conserved molecular markers by intersecting DEGs from NSCs in all three species (Data S5). Nineteen genes were commonly expressed (Fig. 7G), enriched for functions in pallial, hippocampal, and forebrain development, as well as glial cell migration (Fig. 7H). Collectively, our results revealed a conserved transcriptional program and gene set in adult avian NSCs.

## Spatial organization and cellular diversity of adult neurogenesis in the telencephalon

To profile adult telencephalic neural stem cell lineages, we combined EdU labeling (1, 7, 14, and 28 days post-injection) with Smart-seq3xpress single-cell transcriptomics (*64*) on FACS-isolated EdU⁺ pallial cells from adult budgerigars (Fig. 8A). This approach enabled temporal tracking of newborn neurons and non-neuronal progeny across distinct neurogenic stages (fig. S38, A and B). To determine spatial localization, we performed NeuN immunostaining on EdU-labeled brains and identified NeuN⁺EdU⁺ cells as recently generated neurons derived from proliferating stem cells (Fig. 8B). Quantitative analysis showed that adult neurogenesis constitutes ∼0.48% of total brain cells, a proportion similar to that of non-neuronal lineages (0.71%), indicating widespread proliferative activity. Neurogenesis was concentrated in the hyperpallium (26.16% ± 3.83%), nidopallium (34.23% ± 4.58%), mesopallium (16.88% ± 5.77%), and subpallial striatum (16.49% ± 5.55%) (Fig. 8B). In contrast, arcopallium (3.21% ± 1.96%) and entopallium (3.03% ± 1.83%) exhibited markedly lower neurogenic activity, consistent with their roles in fixed sensory-motor pathways (Fig. 8B). This distribution reflects a gradient of neurogenic plasticity: the nidopallium, particularly the NLc region homologous to the HVC in songbirds (*27*), exhibits substantial postnatal neuronal integration (*13, 61*), whereas output regions such as the arcopallium show limited incorporation of new neurons.

**Fig. 8.**
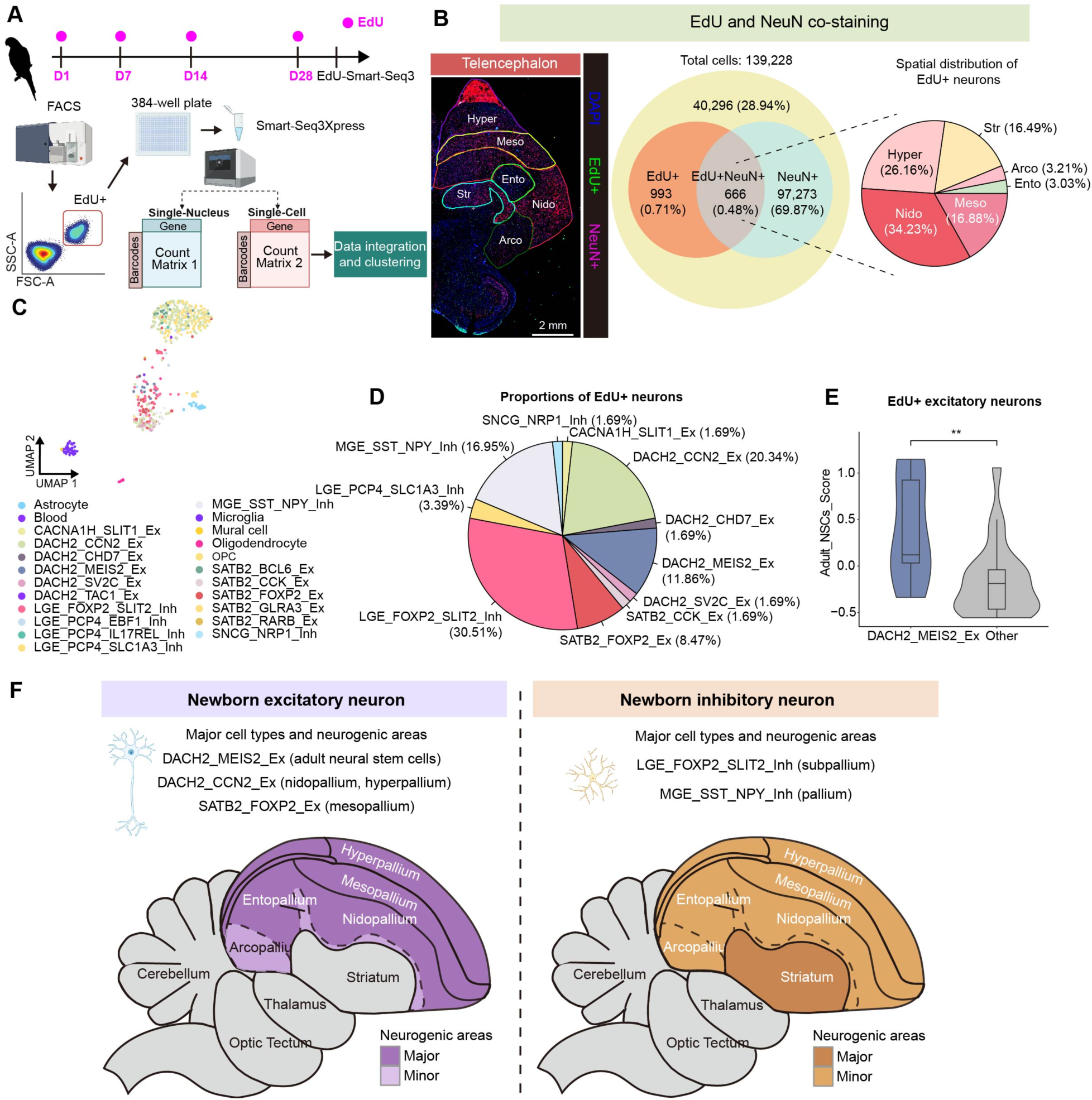
EdU pulse-chase identifies adult-born neuronal subtypes and region-specific neurogenesis in the budgerigar brain. (**A**) Schematic of lineage tracing in the adult budgerigar brain using EdU labeling and Smart-seq3Xpress. EdU was administered at 1, 7, 14, and 28 days before tissue collection. EdU⁺ cells or nuclei were isolated by FACS and subjected to Smart-seq3Xpress for single-cell/nucleus sequencing. (**B**) EdU labeling and NeuN co-staining in the budgerigar pallium. Representative image from two biological replicates are shown; only one hemisphere is shown due to symmetry. Quantification was based on two biological replicates, each with three technical replicates. (**C**) UMAP showing smart-seq clusters of EdU^+^ brain cells, including non-neurons and neurons. (**D**) Proportions of excitatory and inhibitory EdU⁺ neuronal supertypes. (**E**) NSC scores computed using Seurat’s AddModuleScore function for excitatory neuron supertypes. NSC scores computed for excitatory neuron supertypes using Seurat’s AddModuleScore function. Statistical comparisons were performed using the Wilcoxon rank-sum test (p < 0.01). (**F**) Anatomical summary of neurogenic regions across cell types: excitatory neurogenesis is restricted to the pallial region, whereas inhibitory neurogenesis occurs in both pallial and subpallial domains.

Transcriptomic profiling of EdU⁺ cells resulted in 23 cell types that were annotated by label transfer from adult datasets (Fig. 8C). Among newborn neurons, excitatory and inhibitory subtypes were approximately equally represented (Fig. 8D). Inhibitory neurogenesis was dominated by two supertypes: LGE_FOXP2_SLIT2_Inh (30.51%), enriched for learning-related and excitatory interaction modules, and MGE_SST_SPY_Inh (16.95%), associated with axonogenesis and projection development (fig. S39, A and B). These findings suggest that inhibitory neurons are functionally specialized—some modulate network plasticity, while others contribute to structural circuit formation. Excitatory neurogenesis was more heterogeneous, comprising projection neurons (DACH2_CCN2_Ex, 20.34%), stem-like cells (DACH2_MEIS2_Ex, 11.86%), and intratelencephalic neurons (SATB2_FOXP2_Ex, 8.47%). Scoring with our NSC-associated gene module (Fig. 7H) confirmed that DACH2_MEIS2_Ex retained a robust stem cell signature (p < 0.01 vs. other excitatory types; Fig. 8E), consistent with asymmetric division modes generating both self-renewing and differentiating progeny, as observed in mammalian cortical NSCs (*65*).

To assess evolutionary conservation, we compared gene expression modules between avian and mammalian NSCs. DACH2_MEIS2_Ex neurons exhibited strong transcriptional similarity to dentate gyrus (DG) neuroblasts in both mice and humans (Fig. 3E and figs. S28 to S29). Avian NSC gene scores were significantly higher in mammalian DG neuroblasts than in mature granule cells (p < 0.0001; fig. S40A), including canonical markers such as *Dcx* and novel candidates like *Epha5* and *Zbtb18* (fig. S40B). *Epha5* and *Zbtb18* are implicated in cortical development (*66–68*), suggesting their involvement in a core stemness program conserved across amniotes. Our comprehensive mapping of adult avian neurogenesis (Fig. 8F) reveals distinct regional and cellular signatures: excitatory neurogenesis predominantly occurs in the hyperpallium/nidopallium (DACH2_CCN2_Ex) and mesopallium (SATB2_FOXP2_Ex, a member of the SATB2_SOX6_Ex subclass), while being markedly reduced in the arcopallium and entopallium. The maintenance of this neurogenic system appears facilitated by asymmetric division mechanisms preserving the DACH2_MEIS2_Ex stem cell pool (Fig. 7D). In contrast, inhibitory neurogenesis primarily arises from the pallial MGE_SST_SPY_Inh cluster, which shows strong enrichment for neurodevelopmental functions. However, we did not detect definitive inhibitory neural stem cells, highlighting a key question for future investigation. Together, these findings indicate that avian and mammalian NSCs share a common evolutionary origin but have diverged in regulatory architecture and output potential. In mammals, adult neurogenesis is largely restricted to the hippocampal dentate gyrus, the subventricular zone (SVZ), and the olfactory bulb, and generates limited neuronal diversity (*10, 69*). In contrast, the avian telencephalon supports widespread neurogenesis across pallial regions (*69*), producing diverse excitatory and inhibitory lineages. This broader potential may underlie the enhanced neurogenic flexibility and cognitive capabilities observed in birds (*12*).

## Discussion

By integrating spatial and single-cell transcriptomics, we constructed a high-resolution molecular atlas of the adult budgerigar forebrain. This atlas delineates hierarchical molecularly defined regions and cell-type taxonomies across major pallial subdivisions. To place these results in an evolutionary context, we aligned our data with existing single-cell datasets from chicken (*22, 41*) and zebra finch (*43*), enabling phylogenetic tracing across the avian lineage. Furthermore, cross-species comparisons with reptiles (*20, 59*) and mammals (*39, 42, 45, 70, 71*) revealed conserved and divergent trajectories in neurogenesis, lineage allocation, and regional gene expression across amniotes. Based on these integrative analyses, we propose a unified developmental framework for pallial evolution (fig. S44).

Our study uncovers a deeply conserved scaffold of excitatory neuron types shared across mammals, reptiles, and birds, including amygdaloid (Amy), hippocampal (MP), and lateral pallial (LP) populations (*21*) (Fig. 3E and figs. S26, S28 to S30), alongside a core L6b-like module that is retained across species despite vast architectural divergence (*22, 41*). Strikingly, this evolutionary continuity is punctuated by a species-specific innovation in birds: the SATB2_KIAA1217_Ex population, which emerges through developmental co-option of an intratelencephalic transcriptional module, indicating a modular repurposing strategy embedded within avian pallial evolution.

While core cell types remain conserved across lineages, our data reveal two major developmental innovations that likely arose in amniotes ∼320 million years ago. First, we provide evidence for the emergence of a novel precursor population within the dorsal pallium, exhibiting transcriptional profiles and cytoarchitectonic characteristics analogous to PT neurons. This conclusion is supported by our identification of PT-like transcriptional modules in turtles, mammalian L5 PT neurons, and avian DACH2_CCN2 cells (Fig. 3E and figs. S26, S28 to S30). The appearance of this population introduced novel projection pathways absent in earlier vertebrates, while fundamentally challenging the long-held paradigm that PT neurons represent a mammalian-specific neocortical innovation (*72*). Second, our data suggest that IT neurons may derive from two distinct developmental origins – dorsal (dIT) and ventral (vIT) progenitor domains. The dorsal lineage gave rise to species-specific adaptations, including SATB2_SOX6_Ex neurons in avian and L4/5 IT neurons in mammals (fig. S44). Conversely, the ventral lineage underwent substantially enriched within ventral progenitor zones. These evolutionary modifications facilitated: (i) expanded intracortical connectivity (*49*), (ii) and signaled a shift from nuclear to more distributed circuit architectures.

The mammalian-reptilian divergence represents a second critical evolutionary transition, during which reptiles retained the ancestral four-region pallial organization, with ventrally derived IT (vIT) neurons originating predominantly from the VP and migrating radially to form nuclear structures. In mammals, the dorsal expansion of vIT neurogenesis involved substantial tangential migration from VP-derived progenitors into DP territories (*73*), where they underwent radial dispersal. This dual-phase migration pattern (ventral-to-dorsal tangential movement followed by local radial distribution) fundamentally distinguishes mammalian pallial development from the strictly radial migration observed in reptilian VP and subsequent establishment of PT-dependent radial scaffolds. These innovations collectively enabled: (i) the emergence of laminated neocortical and proto-entorhinal circuits, (ii) the evolutionary specialization of vIT neurons into mammalian L2/3 IT populations, and (iii) ultimately the developmental foundation for advanced cognitive circuitry.

In the archosaurian ancestor, which gave rise to both birds and crocodilians, we infer the emergence of a major evolutionary innovation. This transition involves a fundamental shift in pallial organization. Instead of forming discrete regional compartments, the developing pallium adopted a dorsoventral (DV) gradient architecture. We refer to this arrangement as the “Refined Continuum Model” (*17*). In this model, gene expression domains extend continuously across the ventricular closure boundary, giving rise to IT-like mesopallial neurons that share molecular signatures and circuit logic with mammalian entorhinal structures. Spatial transcriptomic analyses of the avian pallium support this model, revealing continuous molecular gradients rather than discrete lineage-restricted domains (Fig. 4). Critically, these patterns mirror observations in crocodilian brains (*32*), underscoring a shared organizational framework within crown archosaurs. Together, these data suggest that the archosaurian pallium was reorganized along a dorsoventral developmental continuum. We propose that this graded architecture liberated novel modes of microcircuit integration, enabling emergent input-output transformations that were evolutionarily constrained by the ancestral compartmentalized plan.

We further propose that avian telencephalon evolution elaborated this ancestral archosaurian framework through two key innovations: (i) fusion of dorsal and ventral mesopallial progenitor zones, forming the bilaterally symmetric LMI that bridges developmental compartment boundaries; and (ii) restriction of excitatory neuron migration to radial trajectories, eliminating ancestral tangential dispersion patterns (*74*). This latter modification established continuous cellular distributions prior to pallial ventricular zone regionalization, thereby enhancing spatial precision and synaptic specificity in microcircuit organization. Strikingly, the conserved molecular identities of output neurons (arcopallium/hippocampus) and input neurons (entopallium/IHA) directly corroborate this developmental model, demonstrating how lineage-specific modifications built upon shared archosaurian circuitry.

Together, these innovations produced a bilaterally symmetrical pallium optimized for efficient computation through tightly coordinated input–output circuits.

Developmental profiling revealed a conserved inside-out neurogenic gradient across both pallial and optic tectum excitatory neurons, mirroring the canonical mammalian cortical maturation pattern. Notably, output-projecting neurons in the inside part-including L5/6, arcopallium, SGC populations that form local microcircuits before projecting to thalamic and brainstem targets (*40, 54, 57*) – exhibited significantly earlier maturation than neurons in other regions (Fig. S45A). These early-born neurons likely serve as “pioneer” cells, providing both structural scaffolding and functional guidance to later-developing neurons, akin to a lead runner setting pace and direction. Intermediate progenitors (e.g., Pallium_Ipc) generated late-maturing excitatory neurons distributed across the nidopallium, mesopallium, and hyperpallium. Combinatorial EdU labeling and Smart-seq3xpress revealed a fundamental divergence in neurogenic potential: whereas mammalian adult neurogenesis is restricted to discrete niches producing limited neuronal subtypes, avian NSCs exhibit expanded developmental plasticity, generating diverse terminal populations throughout multiple pallial domains. Notably, this neurogenic capacity was pronounced in associative regions (hyperpallium/nidopallium/mesopallium) but virtually absent from primary input (entopallium) and output (arcopallium) nuclei. These findings support a model wherein avian NSCs establish experience-modulated microcircuits within the telencephalon – a regulatory mechanism phylogenetically convergent with mammalian olfactory learning pathways. Under baseline conditions, they appear quiescent, but upon environmental challenge, can be mobilized as a “reserve” pool to sustain neural plasticity.

The avian song system exemplifies this developmental logic. In the following, the brain region names are standardized to songbird nomenclature (*27*) (fig. S45B). The robust nucleus of the arcopallium (RA)—a central output node receiving pallial inputs from HVC and lateral magnocellular nucleus of the anterior nidopallium (LMAN)—exhibits precocious maturation with minimal adult neurogenesis. This constrained developmental program enables rapid establishment of innate circuitry resistant to experiential modification, ensuring transmission stability. Conversely, the pallial nuclei HVC and LMAN display delayed maturation coupled with persistent, hormonally regulated neurogenesis (*75*), peaking during breeding seasons to enhance song complexity (*12*). This dual mechanism reveals a functional segregation: hardwired stability in descending motor pathways (RA → nXIIts) versus experience-dependent plasticity in intra-pallial circuit (HVC/LMAN), reflecting evolutionary adaptations for both innate behavior and learned vocal optimization.

Therefore, we proposed the “Pioneer and Reserve Hypothesis” (fig. S45), which argues that amniote brain development employs evolutionarily conserved, complementary strategies: (i) early-maturing pioneer neurons establish fundamental circuitry during critical developmental periods, and (ii) reserved neural stem cells maintain lifelong plasticity through delayed neurogenesis and synaptic refinement. While this dual mechanism appears conserved across brain regions and species, providing both developmental robustness and adult flexibility, key questions remain unresolved. First, the integration mechanisms of newborn neurons into pre-existing pallial circuits require elucidation. Second, the potential conservation of gene regulatory networks (GRNs) governing reserve NSC maintenance across amniotic lineages warrants investigation. Resolving these questions could provide fundamental insights into both the evolution of cortical complexity and the pathogenesis of neurodevelopmental disorders.

## Acknowledgments

We thank Drs. Longqi Liu, Shiping Liu, Muyang Lu, and Zhengyang Wang for their insightful discussions throughout the project. Some data processing and analysis were carried out using the STOmics Cloud platform (https://cloud.stomics.tech). Some figures were created by www.BioRender.com.

## Author contributions

G.J. conceived and supervised the study. H.S. performed most of the analyses with support from Z.F. Y.M. carried out the animal experiments and sequencing. X.G. and L.H. conducted experiments, performed sequencing, and generated data. Y.H., C.L., Y.Z., J.H., R.W., Z.Z., M.W., and D.C. assisted with experiments, data generation, and discussions. Z.F. developed the website for data sharing. Y.W., L.Z., and G.J. acquired funding and co-supervised the project. H.S. and G.J. interpreted the data and wrote the manuscript.

## Data and materials availability

Raw and processed sequencing data have been deposited in the National Genomics Data Center, China (NGDC), under accession number PRJCA038662 *(under controlled access until publication).* Processed datasets with an interactive interface are also available at www.budgiebrain.info (*temporarily accessible during review at* http://60.204.237.173). All custom scripts used for data processing and analysis are available at https://github.com/JiaGSLab/budgiebrain.

## Supplementary Materials

Materials and Methods

Figs. S1 to S45

References (*1–75*)

Data S1 to S5

