## Supplemental Material for "Structural innovations and neurogenic continuity define avian brain development and evolution"

Science
AAAS

Supplementary Materials for

### Structural innovations and neurogenic continuity define avian brain development and evolution

Hongcheng Shan, Yan Mei, Zhenhuan Feng, Xilin Gu, Liu Huang, Yuan Hui, Chunqiong Li, Yingying Zhou, Junjie He, Ran Wu, Zhengkun Zhuang, Mengzhu Wang, Duoyuan Chen, Yongjie Wu, Li Zhang, Guangshuai Jia

**The PDF file includes:**

Materials and Methods
Figs. S1 to S45
References

**Other Supplementary Materials for this manuscript include the following:**

Data S1 to S5

#### **Materials and Methods**

Animals

Adult budgerigars (*Melopsittacus undulatus*) were purchased from certified commercial vendors. Birds were maintained in indoor aviaries with unrestricted access to food and water, under a controlled 12-hour light/dark cycle. Brain tissue was obtained from healthy individuals, with ages detailed in the main text and sample identifiers provided in Data S1. Immediately following euthanasia, brains were either dissected in ice-cold phosphate-buffered saline (PBS) and rapidly frozen in liquid nitrogen, or embedded in optimal cutting temperature (OCT) compound and preserved on dry ice for cryosectioning. For anatomical regionalization, brains were microdissected into four major regions: telencephalon, diencephalon, midbrain, and cerebellum. For developmental studies, fertilized budgerigar eggs were obtained from the same supplier and incubated at 37.5 °C under humidified conditions. Embryonic day 0 (E0) was designated as the day eggs were placed in the incubator. Embryos were harvested at defined stages of development. After dissection in chilled PBS, brains were either snap-frozen in liquid nitrogen or embedded in OCT medium for subsequent histological or molecular profiling.

All procedures were conducted in accordance with institutional and regional guidelines for animal experimentation and were approved by the Ethics Committee of Guangdong Provincial People's Hospital.

Nissl staining and anatomical regions

The experimental procedures were conducted following established protocols for Nissl staining and anatomical delineation in avian brain studies (*1, 2*). Adult budgerigars were deeply anesthetized via intraperitoneal injection of 0.96% sodium pentobarbital (5 mL, adjusted as needed) and transcardially perfused with PBS through the left ventricle until hepatic clearance was observed (liver color transition from red to pale yellow). Brains were rapidly extracted, post-fixed in 4% paraformaldehyde overnight, and cryoprotected in 25% sucrose solution at 4°C for 2–3 days, followed by immersion in 30% sucrose until tissue sedimentation. Dehydrated brains were embedded in OCT compound, frozen at −80°C, and sectioned coronally or sagittally at 20 μm thickness using a cryostat. Sections were mounted on slides, air-dried overnight, and subjected to Nissl staining using the Cresyl Violet method (Solarbio Nissl Stain Kit G1430). Briefly, slides were rinsed in PBS, stained at 56°C for 1 h, differentiated in 30 s, dehydrated through graded alcohols (50%, 75%, 95%, 100%), cleared in xylene, and coverslipped with resin.

Anatomical boundaries of brain regions and nuclei were defined based on cytoarchitectonic features visualized by Nissl staining, supplemented by reference to the revised avian brain nomenclature (*3*) and cross-verified with the budgerigar brain atlas (http://www.brauthlab.umd.edu/atlas.htm). Coronal and sagittal sections were systematically collected using a predefined coordinate system, with the midline sagittal plane as the reference for orientation. Serial sections were aligned anterior-posteriorly, and positional data were recorded relative to the initial reference plane to ensure spatial consistency across the atlas.

Single-nucleus RNA sequencing

Single-nucleus RNA sequencing (snRNA-seq) was performed on microdissected brain regions of budgerigar using two different platforms. For the majority of samples, snRNA-seq was performed using the DNBelab C4 platform (BGI). The single-nucleus suspensions were used for droplet generation, emulsion breakage, bead collection, reverse transcription and cDNA amplification to generate barcoded libraries according to the manufacturer' s protocol. Final libraries were sequenced by the DIPSEQ T7 platform (MGI) with the following sequencing strategy: 30-bp read length for read 1, and 100-bp read length for read 2. The remaining samples were processed using the GEXSCOPE® Single Nucleus Seq technology (Singleron Biotechnologies), following the manufacturer's protocol. Single-nucleus suspensions were loaded onto microfluidic chips for library preparation, and sequencing was performed on an Illumina NovaSeq 6000 platform with 150 bp paired-end reads.

<u>Stereo-seq</u>

Stereo-seq library preparation and sequencing were performed as previously described (*4*). Briefly, brain tissues were cryopreserved, embedded in OCT, and sectioned for spatial transcriptomics analysis. RNA quality was assessed prior to mounting each section onto a STOmics chip. Nucleic acid dye staining was used to evaluate tissue morphology and integrity. Sections were permeabilized at 37 °C for 12 minutes, followed by reverse transcription, second-strand synthesis, and cDNA denaturation. Indexed libraries were then constructed according to the manufacturer's protocol and sequenced on the DIPSEQ T7 platform (MGI).

<u>snRNA-seq data processing</u>

*1. Data quality control and pre-processing*

Raw sequencing data from the BGI and Singleron platforms were initially processed using the respective manufacturer-provided pipelines to generate gene expression matrices. Reads were aligned to the *Melopsittacus undulatus* 6.3 reference genome (release 100). The resulting expression matrices were subsequently analyzed using Seurat (v5.0) (*5*). Potential doublets were identified and removed with DoubletFinder (v2.0.4) (*6*) to mitigate artifacts introduced by droplet-based library construction. Nuclei with fewer than 100 genes detected (nFeature_RNA < 100), total UMI counts outside the range of 300–10,000 (nCount_RNA), or mitochondrial gene content exceeding 5% were excluded. Mitochondrial gene expression was quantified using a curated list of 13 mitochondrial genes (*ND1*, *ND2, ND3*, *ND4, ND4L, ND5, ND6, CYTB, COX1, COX2*, *COX3*, *ATP6, ATP8*). Data were log-normalized, and the top 2,000 highly variable genes (HVGs) were selected using the variance-stabilizing transformation (vst) method.

*2. Cell integration and annotation*

Batch effects arising from platform-specific technical variation (BGI vs. Singleron) were corrected using Seurat's integration pipeline. Briefly, datasets were split by sequencing run (Sample_ID), normalized independently, and anchored via reciprocal PCA (RPCA) to identify cross-dataset correspondences. Integrated data were scaled, and principal component analysis (PCA) was performed on HVGs (dims = 1:30). A shared nearest neighbor (SNN) graph was constructed, and cells were clustered using the Leiden algorithm at a resolution of 0.4 to balance cluster granularity and biological interpretability. Uniform Manifold Approximation and Projection (UMAP) was applied for 2D visualization. Clusters were annotated using canonical marker genes (*7*) derived from avian and mammalian brain atlases: glutamatergic cells (*SLC17A6*), GABAergic cells (*GAD1, GAD2*), medium spiny neurons (*MEIS2*), Purkinje cells (*CA8*), astroependymal cells (*SPEF2*), astrocytes (*SLC4A4, SLC1A3*), astrocyte precursors (*TOP2A*), oligodendrocytes (*PLP1*), oligodendrocyte precursors (*VCAN*), microglia (*CSF1R*), vascular cells (*FLI1*), mural cells (*RGS5*), and blood cells (*F10*).

<u>Annotation of single-cell subclasses and supertypes</u>

To systematically characterize neuronal subpopulations in the adult budgerigar brain, we employed a hierarchical classification framework comprising four levels: class, subclass, supertypes, and clusters, which is consistent with prior single-cell profiling studies in the mouse and chicken brains. The analysis involved four sequential stages: (i) neuronal class re-clustering: RPCA integration was utilized to annotate whole-brain cells at low resolution, followed by the extraction and re-clustering of excitatory and inhibitory neuronal subsets for higher resolution; (ii) resolution selection: The optimal resolution was determined by analyzing average cell purity scores across a range of high-resolution settings; (iii) defining neuronal supertypes: Hierarchical clustering, quality control, and MetaNeighbor (v1.18.0) (*8*) analysis were combined to classify potential neuronal supertypes; and (iv) supertype annotation: Supertypes were named using two high-confidence nervous system marker genes identified during the initial FindAllMarkers analysis. Neuronal subclasses were subsequently determined by analyzing the spatial distribution of marker gene expression. These subclasses were then summarized and integrated with non-neuronal populations to create an integrative representation of the avian brain cellular landscape.

*1. Clustering and annotation of adult telencephalon excitatory neurons*

We began with 115,824 cells and performed integration using RPCA based on Sample_ID information, with 2,000 HVGs selected as anchors. Dimensionality reduction was conducted using the first 40 principal components (PCs) in both FindNeighbors and UMAP analyses. We then systematically tested resolution parameters ranging from 1 to 10, evaluating cluster purity by calculating the average ROGUE (v1.0) (*9*) score for all clusters at each resolution. After comprehensive assessment, we selected Glu_resolution = 4, which optimally balanced cluster purity with avoidance of over-clustering.

Using the resulting clusters, we performed hierarchical clustering on the integrated assay and constructed a cluster tree via the BuildClusterTree function from the “ape” package (v5.7), initially identifying 29 supertypes. Low-quality clusters (Ex5, defined by Feature counts <500 or Counts < 800) were excluded. To assess potential splitting of identical cell types across multiple clusters, we applied MetaNeighbor (4,070 anchor genes) to the remaining 28 Ex clusters, merging those with correlation scores exceeding the threshold of 0.85.

Finally, we annotated 23 supertypes (corresponding to 55 clusters) based on high-scoring marker genes identified by the FindAllMarkers function. These clusters were further consolidated into 10 subclasses according to marker gene expression patterns and original clustering results, yielding a final dataset of 109,395 excitatory neurons (fig. S12).

*2. Clustering and annotation of adult telencephalon inhibitory neurons*

The initial dataset comprised 56,454 inhibitory neurons from the adult telencephalon. Data integration was performed using RPCA (2,000 HVGs as anchors) based on Sample_id. Dimensionality reduction employed the first 40 PCs in FindNeighbors and UMAP. Resolution parameters were tested across a range of 0.5–8, with cluster purity evaluated via average ROGUE scores. The optimal parameter (GABA_integrated_snn_res.1.5) was selected for downstream analysis.

Hierarchical clustering initially revealed 28 inhibitory neuron subtypes (Inh). Marker gene analysis (FindAllMarkers) identified non-inhibitory contaminants (e.g., Inh16, Inh19, Inh28), which were subsequently filtered. MetaNeighbor (4,089 anchor genes) was applied to the remaining clusters, with a correlation threshold of 0.85 used to merge biologically redundant groups. This process identified 17 supertypes (33 clusters), which were further refined into 8 subclasses based on marker expression and initial clustering, resulting in 54,299 high-quality inhibitory neurons (fig. S13).

*3. Clustering and annotation of adult TeO excitatory neurons*

The initial dataset comprised 14,339 excitatory neurons from the adult optic tectum. Data integration was performed using RPCA (2,000 HVGs as anchors) based on Sample_id, followed by dimensionality reduction via FindNeighbors and UMAP (first 40 PCs). To optimize clustering, we tested resolution parameters from 1 to 5 (increments of 0.5), evaluating cluster purity using average ROGUE scores. The optimal resolution (Glu_integrated_snn_res.4) was selected for downstream analysis.

Given the limited cell count per cluster (dataset < 20,000 cells), hierarchical clustering was deemed unsuitable. Instead, we directly applied MetaNeighbor (450 HVGs) to assess inter-cluster correlations, merging groups with a cutoff of 0.8 to yield 29 Ex clusters. Low-quality clusters (Ex3, Ex6; defined by Feature counts < 500 or Counts < 1,000) were excluded.

Using high-scoring marker genes (FindAllMarkers), we annotated 23 supertypes (44 clusters), which were further consolidated into 6 subclasses based on marker expression patterns and initial clustering. The final dataset retained 13,122 high-quality excitatory neurons (fig. S15).

*4. Clustering and annotation of adult TeO inhibitory neurons*

The inhibitory neuron dataset initially contained 17,369 cells. After RPCA integration (2,000 anchor genes) and dimensionality reduction (40 PCs), resolution parameters (1–6, increments of 0.5) were evaluated via ROGUE scores. GABA_integrated_snn_res.2 was selected as optimal.

Analogous to the hierarchical clustering approach, MetaNeighbor (450 genes) identified 25 Inh clusters (correlation cutoff: 0.8). Low-quality clusters (Inh3, Inh5, Inh19; Feature counts < 500 or Counts < 1,000) were removed. Subsequent marker gene analysis (FindAllMarkers) revealed 19 supertypes (32 clusters), refined into 8 subclasses. The final dataset included 14,479 inhibitory neurons (fig. S16).

*5. Clustering and annotation of adult thalamus excitatory neurons*

The initial dataset comprised 22,317 excitatory neurons from the adult thalamus. Data integration was performed using RPCA with 2,000 HVGs as anchors, based on Sample_ID information. Dimensionality reduction was conducted using the first 40 principal components in both FindNeighbors and UMAP analyses.

To optimize clustering, we systematically tested resolution parameters ranging from 1 to 5 (in increments of 0.5), with cluster purity evaluated by calculating average ROGUE scores for each resolution. After comprehensive assessment, Glu_integrated_snn_res.4.5 was selected as the optimal resolution parameter.

During downstream analysis, we identified and removed a small population of contaminating inhibitory neurons from the excitatory neuron dataset. The remaining excitatory neurons were subjected to hierarchical clustering using the “ape” package, yielding 17 preliminary Ex clusters. We subsequently excluded low-quality clusters (Ex3, defined by Feature counts < 500 or Counts < 800).

Cluster merging was performed using MetaNeighbor with a correlation cutoff of 0.80 to consolidate biologically similar cell types. Based on high-scoring marker genes identified by FindAllMarkers, we annotated 12 supertypes corresponding to 36 clusters. These were further consolidated into 6 subclasses according to marker gene expression patterns and initial clustering results, resulting in a final dataset of 18,236 high-quality excitatory neurons (fig. S18).

*6. Clustering and annotation of adult thalamus inhibitory neurons*

The inhibitory neuron analysis began with 6,508 cells. Following RPCA integration (2,000 anchor genes) and dimensionality reduction (40 PCs), we evaluated resolution parameters from 1 to 5 (0.5 increments) using ROGUE scores, ultimately selecting GABA_integrated_snn_res.3.5 for downstream analysis.

We identified and removed a minor population of non-inhibitory neuronal contaminants during quality control. The purified inhibitory neurons were analyzed using MetaNeighbor (v4.066) with a 0.80 correlation cutoff, identifying 21 Inh clusters.

Marker gene analysis (FindAllMarkers) revealed 15 distinct supertypes (31 clusters), which were further organized into 5 subclasses based on expression patterns and clustering topology. The final curated dataset contained 5,909 inhibitory neurons (fig. S19).

*7. Clustering and annotation of adult cerebellar excitatory neurons*

The initial dataset comprised 34,855 cerebellar excitatory neurons. To address substantial batch effects observed across samples, we performed RPCA integration using Sample_id information with 2,000 highly variable genes as anchors (k.anchor = 20). Dimensionality reduction was conducted using the first 40 principal components for both FindNeighbors and UMAP visualization. We tested resolution parameters ranging from 0.3 to 2 and selected Glu_integrated_snn_res.1.2 as optimal based on ROGUE-calculated cluster purity scores.

Given the overall lower quality of cerebellar excitatory neurons (evidenced by reduced feature and count numbers), we bypassed hierarchical clustering and directly applied MetaNeighbor (489 anchor genes) to assess cluster correlations. This analysis identified 14 Ex clusters, from which we excluded low-quality populations (Ex1 and Ex18 [containing only 4 cells]) with Feature counts <300 or Counts <500. Using high-confidence marker genes identified by FindAllMarkers, we annotated 4 supertypescorresponding to 17 clusters. These were further consolidated into 3 major celltypes based on marker gene expression patterns and clustering topology, yielding a final dataset of 26,947 high-quality excitatory neurons (fig. S21).

*8. Clustering and annotation of adult cerebellar inhibitory neurons*

The inhibitory neuron analysis began with 3,374 cells. Similar batch effect correction was applied using RPCA integration (2,000 HVGs, k.anchor = 20) followed by dimensionality reduction (40 PCs). Resolution parameters (0.1–2) were evaluated using ROGUE scores, with GABA_integrated_snn_res.1.4 selected for downstream analysis. MetaNeighbor (450 genes) identified 13 Inh clusters, from which we removed one low-quality group (Inh2; Feature counts < 500 or Counts < 900).

Marker gene analysis revealed 8 distinct supertypes (17 clusters), which were organized into 3 subclasses based on expression patterns and cluster relationships. The final curated dataset contained 2,961 inhibitory neurons (fig. S21).

<u>Spatial transcriptomics (ST) data pre-processing</u>

For the budgerigar brain Stereo-seq data, the expression matrix was processed by dividing the tissue into non-overlapping spatial bins, each covering an area of 100 × 100 DNA nanoballs (DNBs), corresponding to a physical resolution of approximately 50 μm (referred to as bin100 data). Transcripts of the same gene were aggregated within each bin to generate a spatially resolved expression profile.

This binning strategy ensured high-resolution mapping of gene expression patterns across the brain while maintaining computational efficiency for downstream analyses.

<u>ST data annotation using spatial leiden</u>

Spatial transcriptomic data were clustered using Stereopy (v0.6.0) with spatially constrained (*10*) Leiden algorithm (spatial leiden) in high resolution, incorporating both gene expression profiles (top 2,000 HVGs) and spatial neighborhood information (fig. S5A). Cluster boundaries were anatomically validated against the Budgerigar Brain Atlas and Nissl-stained references (fig. S3). Gaussian smoothing (smooth_threshold = 90) was applied to enhance signal-to-noise ratio and sharpen domain boundaries, with resulting spatial scores normalized to a 0-10 scale (fig. S5B).

<u>Axial spatial genes scoring in budgerigar telencephalon</u>

Spatial axial scores were computed for dorsoventral (Y-axis, $\sigma DV$) and mediolateral (X-axis, $\sigma ML$) dimensions across 10 coronal Stereo-seq sections (0.3–11 mm) of the budgerigar telencephalon. For each gene, axial specificity was quantified as the normalized standard deviation of its expression distribution relative to the spatial centroid ($\mu X$, $\mu Y$) of all spots in a section:

$$\sigma_{ML}=\sqrt{\frac{\sum(x_i-\mu_X)^2}{N}}\quad \sigma_{DV}=\sqrt{\frac{\sum(y_i-\mu_Y)^2}{N}}$$

where $\sigma ML$ and $\sigma DV$ represent mediolateral and dorsoventral specificity scores, respectively; $xi$ and $yi$ are the spatial coordinates of the $i$-th spot expressing the gene; $\mu X$ and $\mu Y$ denote the mean centroid coordinates along X/Y-axes for all spots in the section; $N$ is the number of spots with detectable gene expression. Top-scoring genes were clustered based on $\sigma ML$ and $\sigma DV$ values to identify axial-enriched transcriptional patterns (*11*).

<u>Pearson correlation analysis</u>

Single-cell datasets from multiple brain regions were integrated and analyzed to assess transcriptional similarity across neuroanatomical domains. Annotated datasets (.h5ad files) were loaded using scanpy (v1.9.2), concatenated into a unified AnnData object with batch identifiers, and saved for reproducibility. Brain regions were reclassified into broader anatomical categories (e.g., *Nido*, *Str*, *Hp*) using a predefined mapping schema that aggregated fine-grained cluster labels (e.g., "NF", "NI") into functionally coherent groups. Gene expression matrices were grouped by original cluster labels, and mean expression values per cluster were computed to generate region-specific transcriptional profiles.

Pearson correlation coefficients between all region pairs were calculated from the mean expression matrix using pandas (v1.3.4), and the resultant correlation matrix was visualized as a heatmap with R package pheatmap (v1.0.12). The heatmap was annotated with correlation values, scaled between -1 and 1.

<u>Enrichment analysis of pathways and custom annotation package construction</u>

Pathway enrichment analyses were restricted to one-to-one orthologs, identified using eggNOG-mapper v5 (--tax_scope auto, --target_orthologs one2one, --seed_ortholog_evalue 0.001, --seed_ortholog_score 60, --query-cover 20, --subject-cover 0) with protein sequences from the budgerigar (*Melopsittacus undulatus*). Gene Ontology (GO) and KEGG pathway annotations were extracted from the eggNOG output file (MM_s5dlclua.emapper.annotations.tsv), including gene identifiers (GID), GO terms, KEGG Orthology (KO), and pathway associations. GO terms were expanded into individual entries, while KEGG pathways were standardized to "KO" nomenclature using a locally curated reference (kegg_pathway.txt).

To enable species-specific enrichment analyses, a custom R package (org.Mundulatus.eg.db) was built using AnnotationForge (v1.40.2), integrating gene-to-GO, gene-to-KEGG mappings, and taxonomic identifiers. The package was validated by confirming database integrity and annotation retrievability. Enriched GO categories were statistically assessed using the clusterProfiler package (v4.6.0), with significance thresholds set at $p < 0.05$. This workflow ensured reproducibility and taxonomic specificity for functional profiling of budgerigar transcriptomic data.

<u>Cross-species analysis</u>

*1. SAMap*

Cross-species cell type alignment was performed using SAMap (v1.0.16) to systematically compare transcriptional profiles across amniote-species (*12-17*). Subsampled datasets were processed through SAMap's standardized workflow (https://github.com/atarashansky/SAMap), aligning orthologous genes and computing pairwise similarity scores between species. The resultant correlation matrix was visualized as a heatmap with R package pheatmap. The heatmap was annotated with correlation values, scaled between 0 and 1.

*2. Label transfer based on Gene-specific-index (GSI)*

Cell type annotation transfer across species was performed using the Seurat integration pipeline to project chicken (*Gallus gallus*) and zebra finch (*Taeniopygia guttata*) brain single-nucleus RNA-seq datas onto a reference dataset of glutamatergic neurons from the budgerigar (*Melopsittacus undulatus*) telencephalon. The reference dataset (adult_Tel_Glu), query chicken (Gg_adult_snRNA_seg_srt) (*18*) and zebra finch datasets (*7*) were subset to shared gene specificity index (GSI) (*14*), ensuring compatibility for cross-species comparisons. Feature selection and scaling were applied to the reference dataset, and transcriptional anchors between species were identified using FindTransferAnchors with default parameters. Cell type labels (supertype_annotation) from the budgerigar reference were transferred to the chicken/zebra finch datasets via the TransferData function, which computes prediction scores for each query cell based on transcriptional similarity to reference cell states (threshold = 0.5). The results generated a cross-avian-species cell type correspondence table (fig. S26E).

<u>Spatial mapping of telencephalic single-cell neuronal data</u>

snRNA-seq data from the budgerigar telencephalon were integrated with spatially resolved transcriptomic data using the Robust Cell Type Decomposition (RCTD) algorithm implemented in the "spacexr" R package (v2.2.1) (*19*). Neuronal subpopulations were annotated based on previously defined subclasses and supertype labels. To balance cell type representation, scRNA-seq data were downsampled by randomly selecting up to 100 cells per supertype cluster (seed = 42), excluding non-neuronal populations (e.g., blood, mural cells, glia). Downsampled counts and metadata were exported as a digital gene expression (DGE) matrix and annotation tables for subclass- and supertype-level analyses.

Spatial transcriptomic data from coronal brain sections were preprocessed to remove vascular-associated spots, and bead coordinates were extracted to map spatial locations. The RCTD pipeline was executed by constructing a Reference object from scRNA-seq DGE and metadata, and a SpatialRNA object from spatial counts and coordinates. Cell type proportions were estimated in "full" doublet mode (max_cores = 8), with weights normalized to sum to 1 across cell types. Normalized spatial weights were exported for downstream visualization. The integrative and reference-informed tissue segmentation (IRIS, v1.0) analysis (*20*) was performed to validate the spatial distribution patterns of RCTD results.

Developmental cell maturity prediction and subtype annotation via label transfer

To assess the maturity of developing telencephalic glutamatergic neurons, cell type labels from adult budgerigar reference data were projected onto developmental-stage cells using Seurat's label transfer framework (*21*). The reference dataset (adult supertypes) and the query developmental dataset were subset to shared genes to enable cross-stage transcriptional alignment. The FindTransferAnchors function was applied to identify integration anchors between the adult (reference) and developmental (query) datasets, using the "RNA" assay and default parameters.

Developmental cell type labels were transferred to the adult dataset via TransferData, generating prediction scores for each adult cell reflecting its similarity to developmental-stage transcriptional states (cutoff: 0.8). Predicted labels and confidence scores were appended to the adult dataset metadata, enabling retrospective evaluation of developmental maturity in adult cells. Subsequently, immature neuronal clusters lacking adult-like transcriptional profiles were annotated by leveraging subclass-specific marker genes from the adult reference. This dual approach resolved developmental trajectories of telencephalic excitatory neurons into distinct Dev subtypes, capturing transitional and mature states.

All analyses utilized Harmony-corrected UMAP embeddings to minimize batch effects, with reproducibility ensured by fixed random seeds and version-controlled dependencies.

Pseudotime analysis

Pseudotime trajectory analysis of developing budgerigar telencephalic glutamatergic neurons was performed using Monocle3 (v1.3.1). The preprocessed Seurat object was converted to a cell_data_set (CDS) structure while retaining raw counts, metadata (including Sample_ID), and gene annotations. After normalization and scaling, principal component analysis (PCA) was conducted using 100 dimensions, followed by batch correction via the align_cds function using "Sample_ID" as the alignment key. Dimensionality reduction was achieved through UMAP, with cells clustered at low resolution to preserve broad developmental transitions. A principal graph was constructed to model transcriptional dynamics, with the root node algorithmically determined based on Progenitor_Ex cells (putative stem cell population) by identifying the most frequent nearest graph vertex. Pseudotime values were calculated relative to this root, and Harmony-corrected UMAP coordinates were integrated for visualization. Complementary analyses included CytoTRACE2 (v1.0.0) (*22*) (https://github.com/digitalcytometry/cytotrace2) for potency assessment and scVelo (v0.2.5) (*23*) (https://scvelo.readthedocs.io) for RNA velocity analysis, together providing a comprehensive reconstruction of glutamatergic neuron differentiation pathways while accounting for technical variability.

Venn diagram analysis of DEGs

Differentially expressed genes (DEGs) between the adult DACH2_MEIS2_Ex and developmental Pallium_Ipc subtypes were identified using the FindAllMarkers with a threshold of |log2FC| > 0.5. The resulting DEG lists were visualized as a Venn diagram using the online tool Venn Analysis (http://www.ehbio.com/test/venn) (*24*) to compare overlapping and unique gene signatures between the two clusters.

EdU labeling

To label proliferating cells, adult animals received intraperitoneal injections of 5-ethynyl-2′-deoxyuridine (EdU; Thermo Fisher A10044) at a dose of 10 mg/kg body weight. EdU was dissolved in sterile phosphate-buffered saline to a final concentration of 5 mg/mL. Injections were administered on days 1, 7, 14, and 28.

Smart-seq3xpress from EdU labeled budgerigar brain tissues

*Frozen tissue for snRNA-seq:*

Frozen budgerigar brain tissues (~10–30 mg) were dissected and homogenized in ice-cold Nuclei EZ Lysis Buffer (Sigma, N3408) supplemented with 1% BSA, 1 mM DTT, and 1 U/μl RNase Inhibitor (40 U/μl, added at 25 μl per 1 ml buffer). Mechanical dissociation was performed using a Dounce homogenizer: 5–10 strokes with a loose pestle (A) followed by 15–20 strokes with a tight pestle (B) on ice. Lysates were incubated for 5 min on ice, filtered through a 30-μm cell strainer, and centrifuged at 500 × g (4°C, 7 min). The pelleted nuclei were washed with Nuclei Suspension Buffer (NSB: 10 mM Tris-HCl pH 7.5, 10 mM NaCl, 3 mM $MgCl_2$, 0.1% Tween-20, 1% BSA, 1 U/μl RNase Inhibitor) and resuspended in reaction buffer containing 1% SUPERase In RNase Inhibitor (Thermo Fisher) for subsequent EdU labeling.

*Fresh tissue for scRNA-seq:*

Adult budgerigar brain tissues were sliced (150–200 μm) in carbogen-bubbled (95% $O_2$/5% $CO_2$) artificial cerebrospinal fluid (ACSF), digested with activated papain (25–30 U/mL) and DNase I (~400 U/mL) at 37°C for 30 min, and gently triturated using fire-polished Pasteur pipettes.

*EdU labeling and FACS isolation:*

For both nuclei and single-cell suspensions, EdU detection was carried out using the Click-iT™ Plus EdU Alexa Fluor™ 488 Imaging Kit (Thermo Fisher Scientific, C10632). Samples were fixed with Click-iT fixative and permeabilized using a saponin-based reagent. The EdU reaction cocktail was applied for 30 min at room temperature in the presence of 1% SUPERase In RNase Inhibitor to preserve RNA integrity. For viability assessment, DRAQ7 (1:100) was used for nuclei. $EdU^+$ nuclei or cells were sorted using a BD FACSAria III-2 cell sorter (BD Biosciences) into 384-well plates preloaded with lysis buffer.

Smart-seq3xpress library preparation and sequencing were performed as previously described (*25*).

Signature scoring of adult bird neural stem cells using AddModuleScore

To evaluate the transcriptional signature of neural stem cell populations in adult avian brains, a gene set scoring approach was implemented using the AddModuleScore function from the Seurat. The predefined gene set (DACH2_MEIS2_Ex_genes), representing a neural stem cell marker signature, was scored against the $EdU^+$ excitatory neuronal dataset. Background noise correction was performed by sampling 100 control genes matched for expression magnitude. Cells were classified into two groups based on their predicted identities: "DACH2_MEIS2_Ex" (potential NSCs) and "Other" (other populations). The Wilcoxon rank-sum test was applied to compare signature scores between groups.

HCR RNA-FISH

Hybridization chain reaction RNA fluorescence *in situ* hybridization (HCR RNA-FISH) was performed on 20-μm cryosections using DNA probes and hairpin amplifiers synthesized by Sangon Biotech (Shanghai, China). Probe sequences are listed in Data S2. All probes were designed against the longest expressed transcript of each target gene. To minimize off-target binding, we performed BLAST against the budgerigar genome to exclude homologous regions using a MATLAB script (https://github.com/uhlen-lab/TRISCO). HCR RNA-FISH was performed according to the Molecular Instruments protocol (v3.0) for frozen tissue samples. Fluorescence imaging was performed using an Akoya/PhenoImager Fusion microscope with a 20× objective.

EdU and NeuN Immunofluorescence

Ten-micrometer frozen sections were processed for EdU and NeuN double labeling. Cells incorporating EdU were detected using the Click-iT™ Plus EdU Alexa Fluor™ 488 Imaging Kit (Thermo Fisher Scientific C10632) according to manufacturer's instructions. Antigen retrieval was performed in Tris-EDTA buffer (pH 9.0) by microwave treatment at high power for 8 minutes, followed by a 7-minute pause, then 8 minutes at medium-low power. Sections were blocked with 5% fetal bovine albumin in PBS for 30 minutes at room temperature, then incubated overnight at 4°C with anti-NeuN (1:1,000; Proteintech 26975-1-AP). After washing, sections were incubated for 50 minutes at room temperature with HRP-conjugated goat anti-rabbit IgG (HUABIO HA1001). Opal 620 Reagent (Akoya FP1495001KT) was used for signal amplification, and nuclei were counterstained with DAPI (5 µg/ml; Sigma D9542). Imaging was performed on an Akoya/PhenoImager Fusion microscope at 20× magnification.

#### Supplementary Figure 1

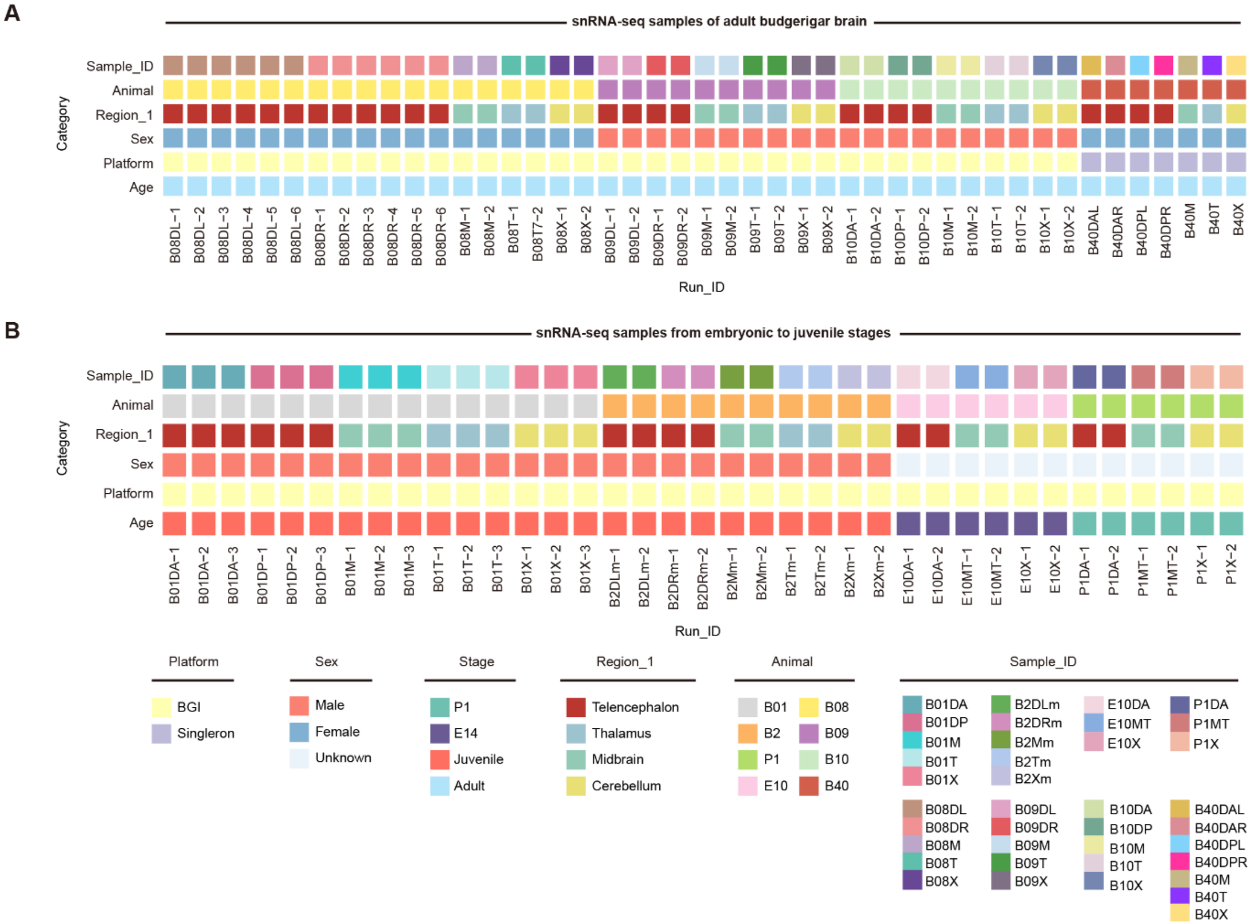


**Fig. S1. Overview of snRNA-seq samples from the budgerigar brain across embryonic stage to adulthood.**

(**A**) snRNA-seq samples from adult stage. (**B**) snRNA-seq samples across three developmental stages. Each column represents a sequenced sample labeled by unique Run_ID (Technical replicate identifier).Platform: Single-cell sequencing platform (BGI or Singleron); Sex: Biological sex (male, female, or unknown for embryonic samples); Stage: Developmental stage (E14 [embryonic day 14], P1 [postnatal day 1], Juvenile, Adult); Region_1: Sampled brain region; Animal: Biological replicate identifier; Sample_ID: Unique sampling event identifier.

#### Supplementary Figure 2

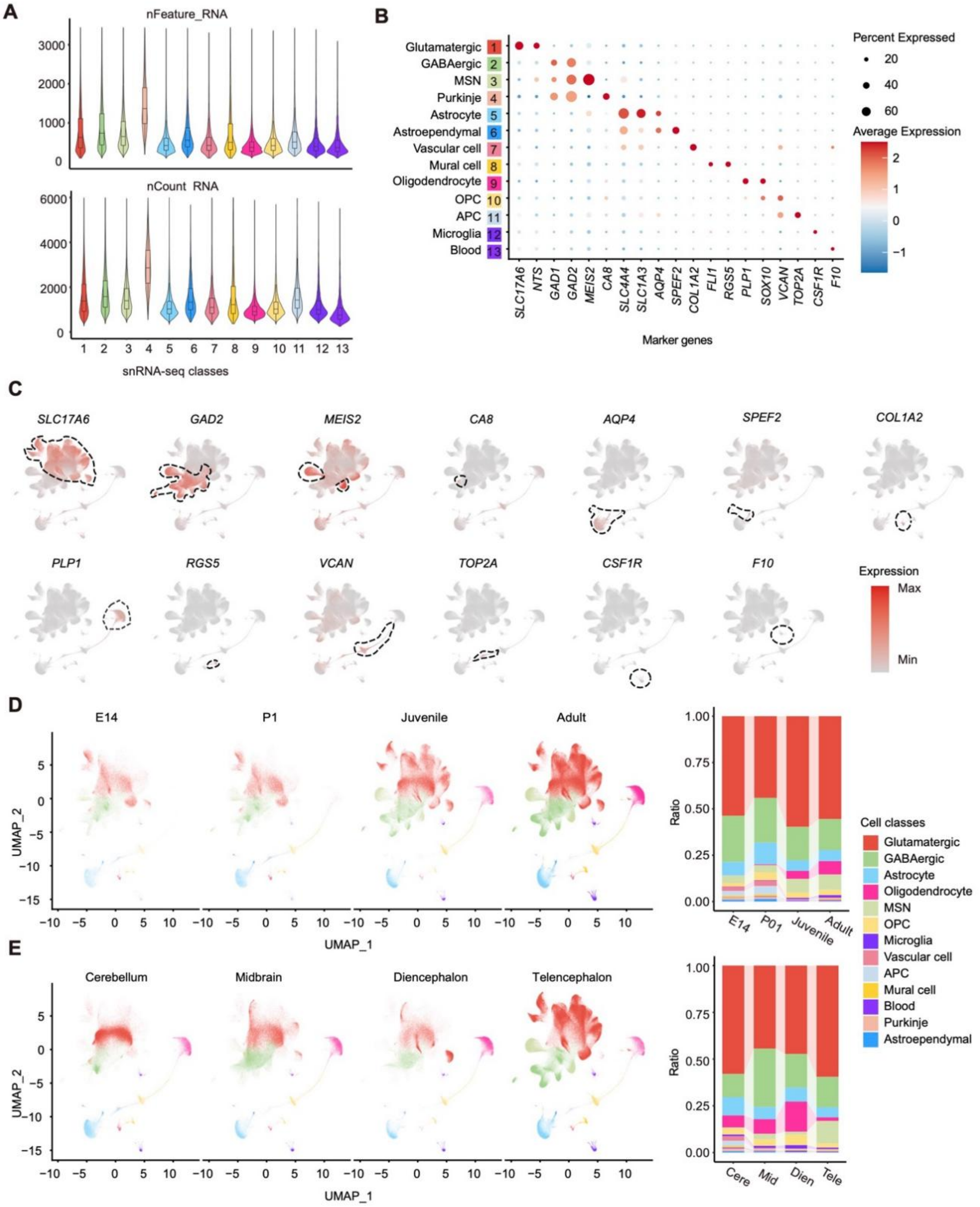


**Fig. S2. Molecular classification of major brain cell classes in the budgerigar.**
(**A**) Violin plots of gene and transcript counts per nucleus across cell classes from snRNA-seq. (**B**) Dot plot of canonical marker expression, showing scaled expression (color) and detection frequency (dot size). (**C**) UMAPs of selected marker genes; high-expression domains are outlined with dashed contours. (**D**) UMAP and barplots showing cell class composition across four developmental stages. (**E**) UMAP and barplots showing cell class distribution across four brain regions.

Supplementary Figure 3

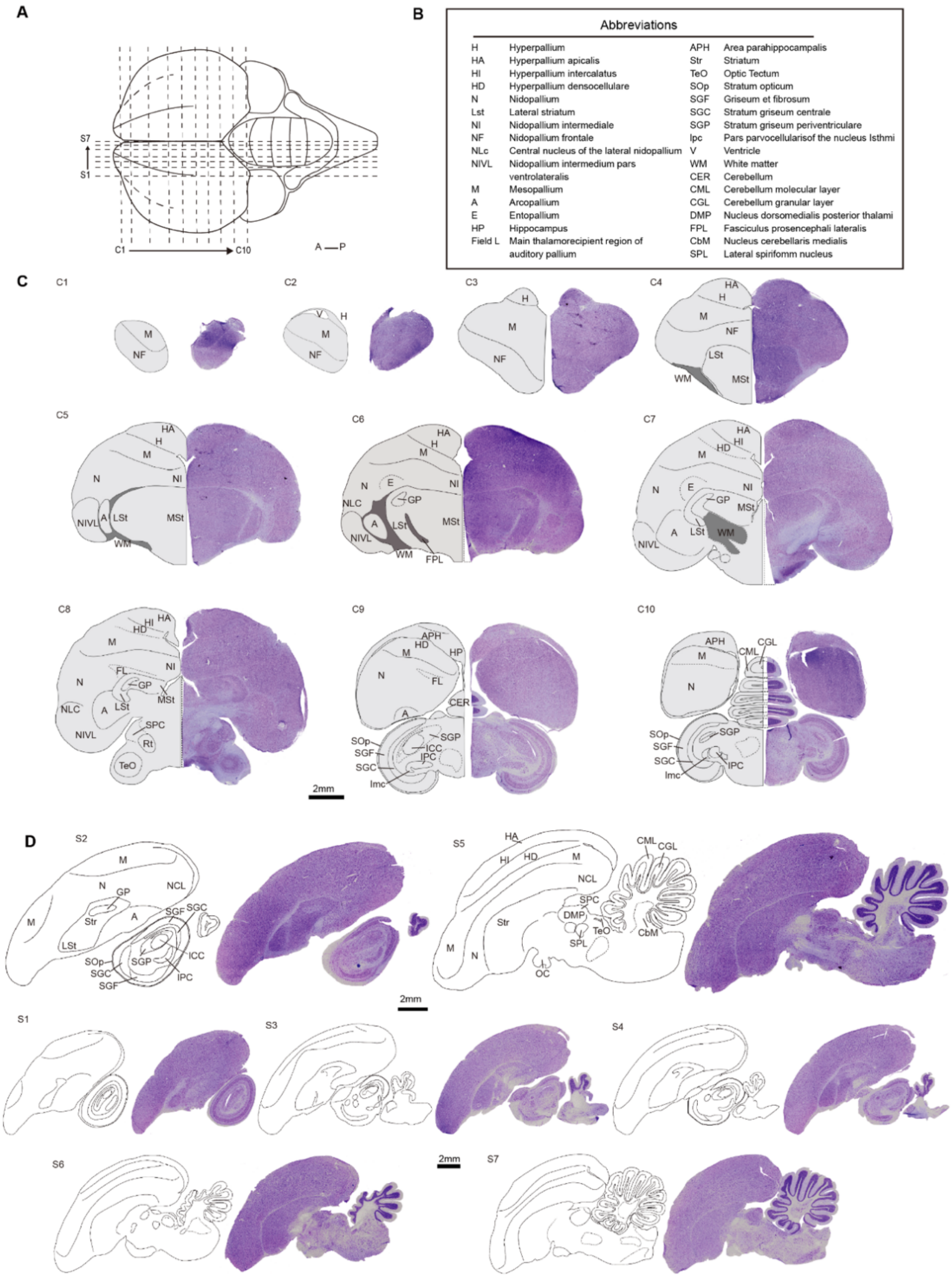


**Fig. S3. Nissl-staining based anatomical reference atlas of the budgerigar brain.**
(**A**) Sampling strategy for Nissl staining. Coronal sections (C1–C10) and sagittal sections (S1–S7) were collected as indicated. (**B**) Abbreviations of anatomical structures based on the nomenclature for the avian brain. (**C, D**) Nissl-stained sections with corresponding schematic parcellations. Major regions were manually annotated based on cytoarchitectonic landmarks.

Supplementary Figure 4

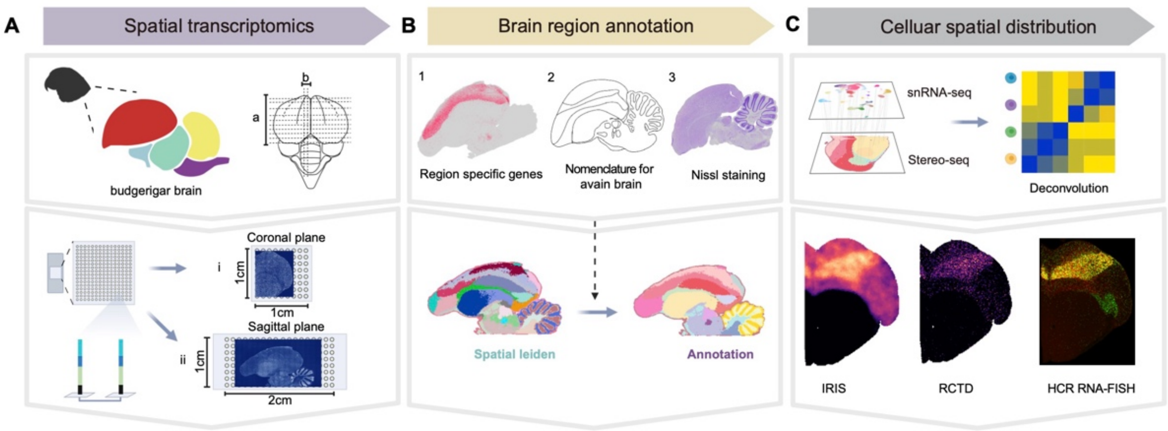


**Fig. S4. Workflow of spatial transcriptomics analysis in the budgerigar brain.**
(**A**) Stereo-seq was performed on 12 brain sections, including 10 coronal (i) and 2 sagittal (ii) planes. Owing to bilateral symmetry, coronal profiling was conducted on one hemisphere. (**B**) Brain regions were annotated based on established avian brain nomenclature (www.brauthlab.umd.edu/atlas.htm). Regional boundaries were manually verified using region-specific marker genes (1) and Nissl-stained adjacent or parallel sections (2–3). Spatial Leiden clusters (ST-level 3) were subsequently merged into broader functional regions (ST-level 2) based on marker expression. (**C**) Cellular spatial distribution was inferred via deconvolution of Stereo-seq using matched snRNA-seq data. Results were validated by HCR RNA-FISH.

Supplementary Figure 5

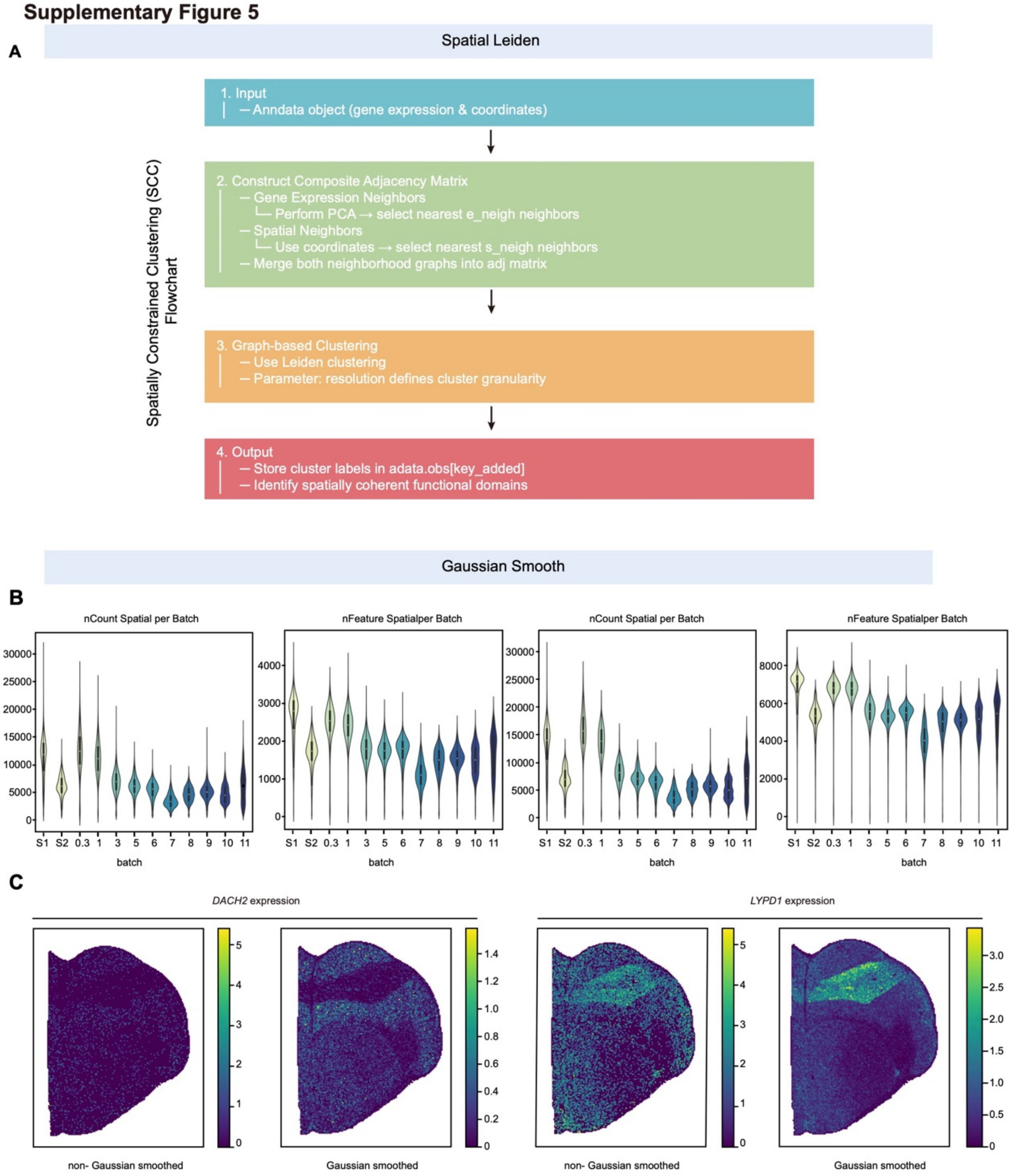


**Fig. S5. Spatial clustering and signal enhancement in budgerigar spatial transcriptomic datasets.**

(**A**)Schematic diagram of the Spatial Leiden workflow using spatially constrained clustering (SCC) (*26*). (**B**) Violin plots showing total transcript counts (nCount) and detected gene features (nFeature) per ST slice across batches, before (left) and after (right) Gaussian smoothing. Smoothing reduces batch variability while preserving local transcriptomic complexity. (**C**) Gaussian smoothing enhances spatial signal-to-noise ratio and delineates anatomical boundaries more sharply.

Supplementary Figure 6

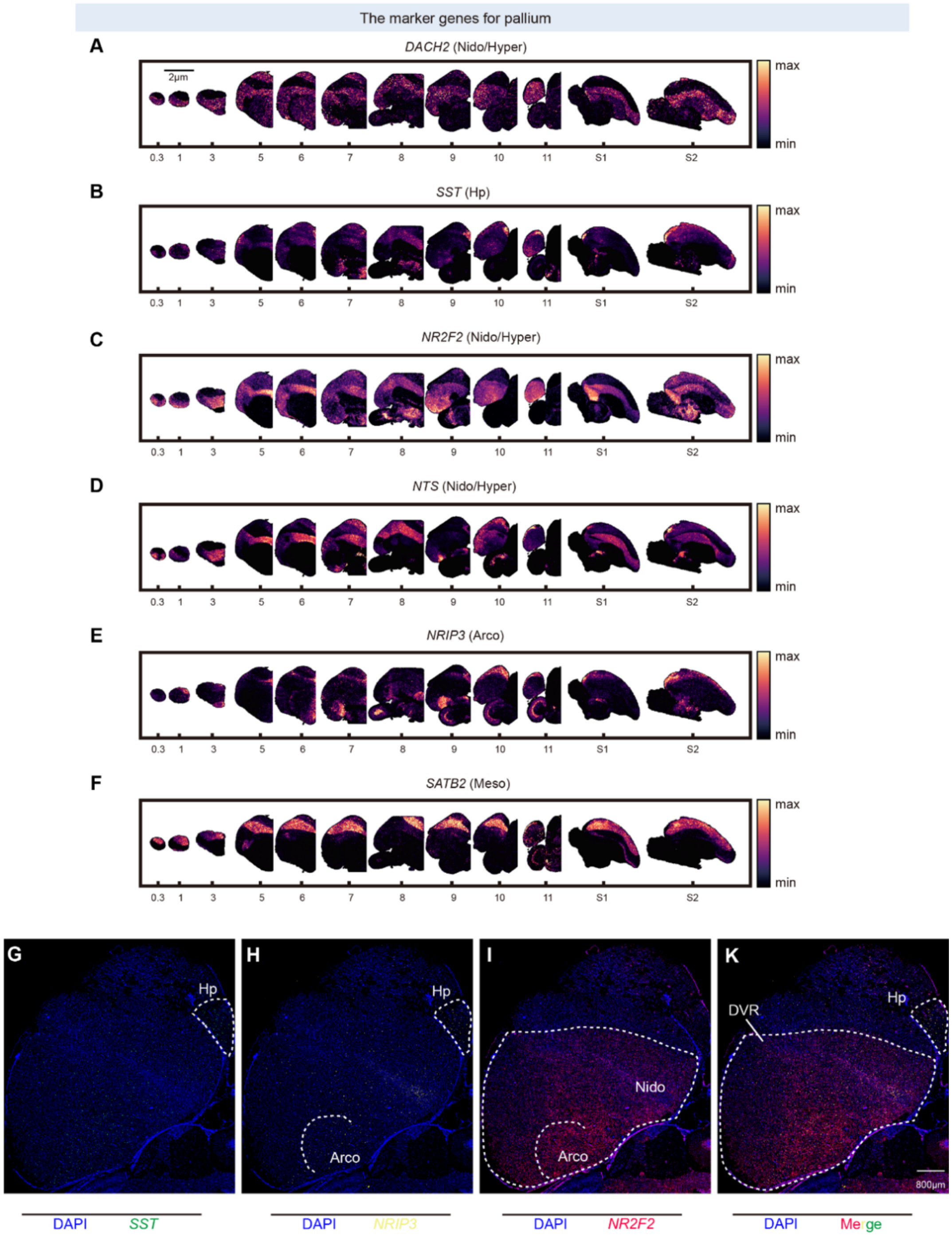


**Fig. S6. Spatial expression of canonical pallial marker genes in the budgerigar brain.**
(**A–F**) Spatial transcriptomic maps showing the distribution of six pallium-associated markers across the coronal ST series. Expression intensity is color-coded from low (black) to high (yellow). Each gene is associated with a known pallial subdivision: *NR2F2*, *DACH2,* and *NTS* (nidopallium/hyperpallium), *SST* (hippocampus), *SATB2* (mesopallium), and *NRIP3* (arcopallium). (**G–K**) HCR RNA-FISH validation of selected markers (*SST*, *NRIP3*, *NR2F2*) in matched coronal sections.

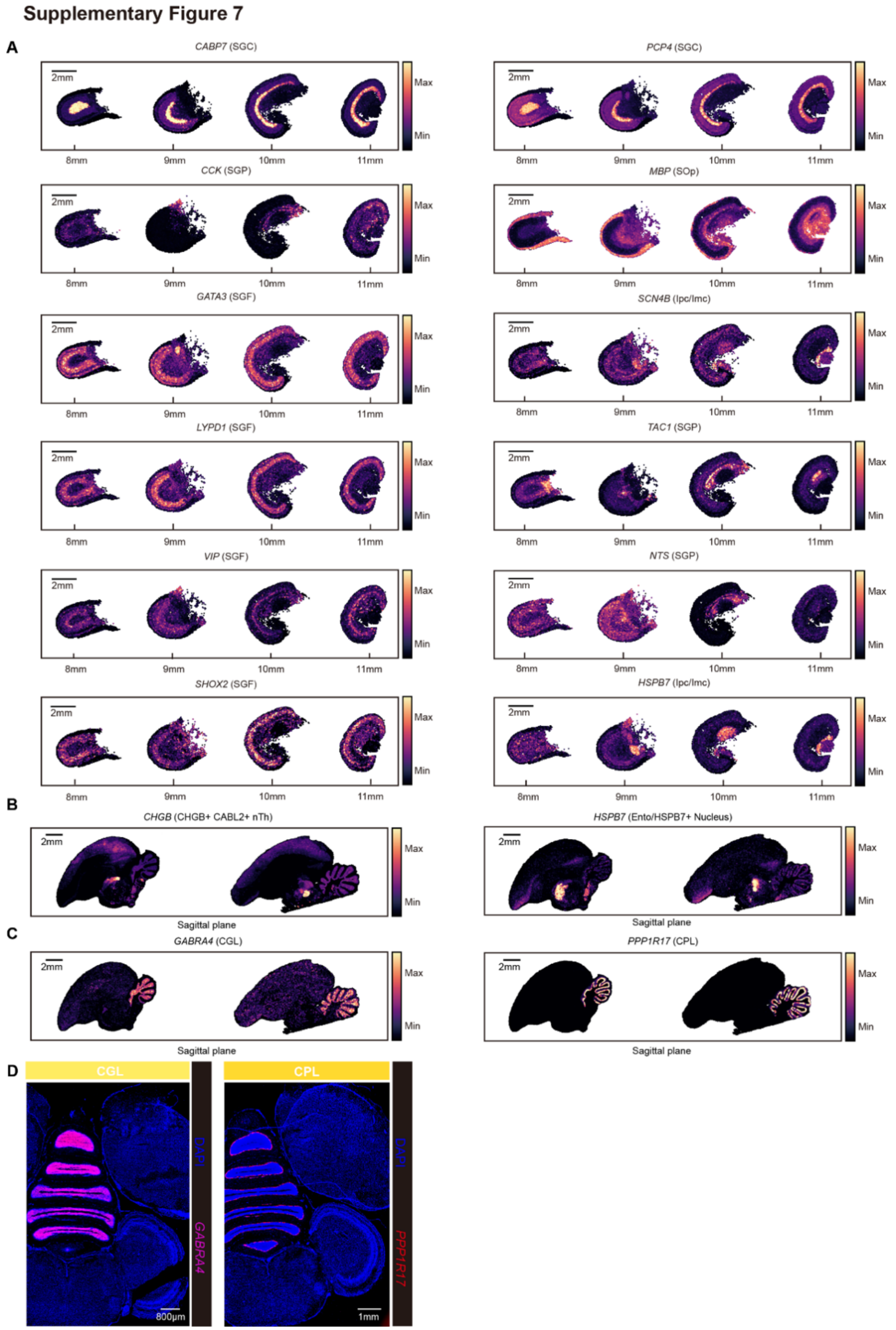


**Fig. S7. Spatial expression of region-specific markers in the budgerigar brain.**
(**A–C)** Spatial transcriptomic maps of canonical markers for midbrain (**A**), diencephalon (**B**), and cerebellum (**C**). Color intensity reflects relative expression levels across developmental stages. (**D**) HCR RNA-FISH validation of cerebellar markers *GABRA4* (CGL) and *PPP1R17* (CPL).

**Supplementary Figure 8**

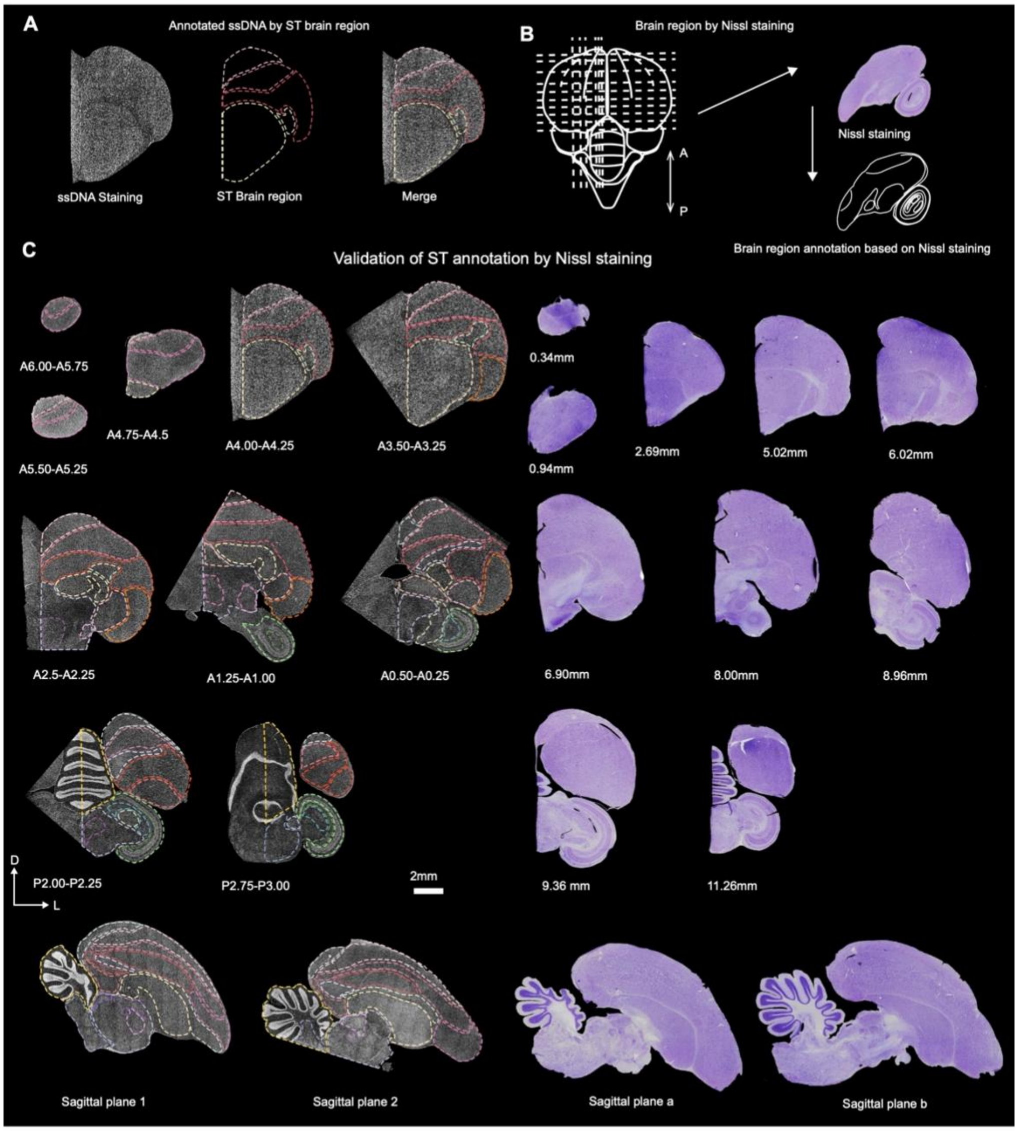


**Fig. S8. Alignment of molecular and anatomical brain regions via ssDNA and Nissl staining.** **(A)** Overlay of ST-defined brain regions onto ssDNA-stained sections. Brain boundaries were manually projected based on ST cluster maps. **(B)** Schematic workflow of anatomical region annotation using Nissl-stained coronal and sagittal sections. **(C)** Validation of ST-based regional annotation using both ssDNA (left) and Nissl (right) signals across coronal and sagittal planes. Dashed lines indicate anatomical boundaries consistently observed across modalities, reflecting the alignment between transcriptional domains and cytoarchitectural features. Spatial coordinates were determined using both an online atlas (www.brauthlab.umd.edu/atlas.htm) and physical distance measurements from the olfactory bulb.

#### Supplementary Figure 9

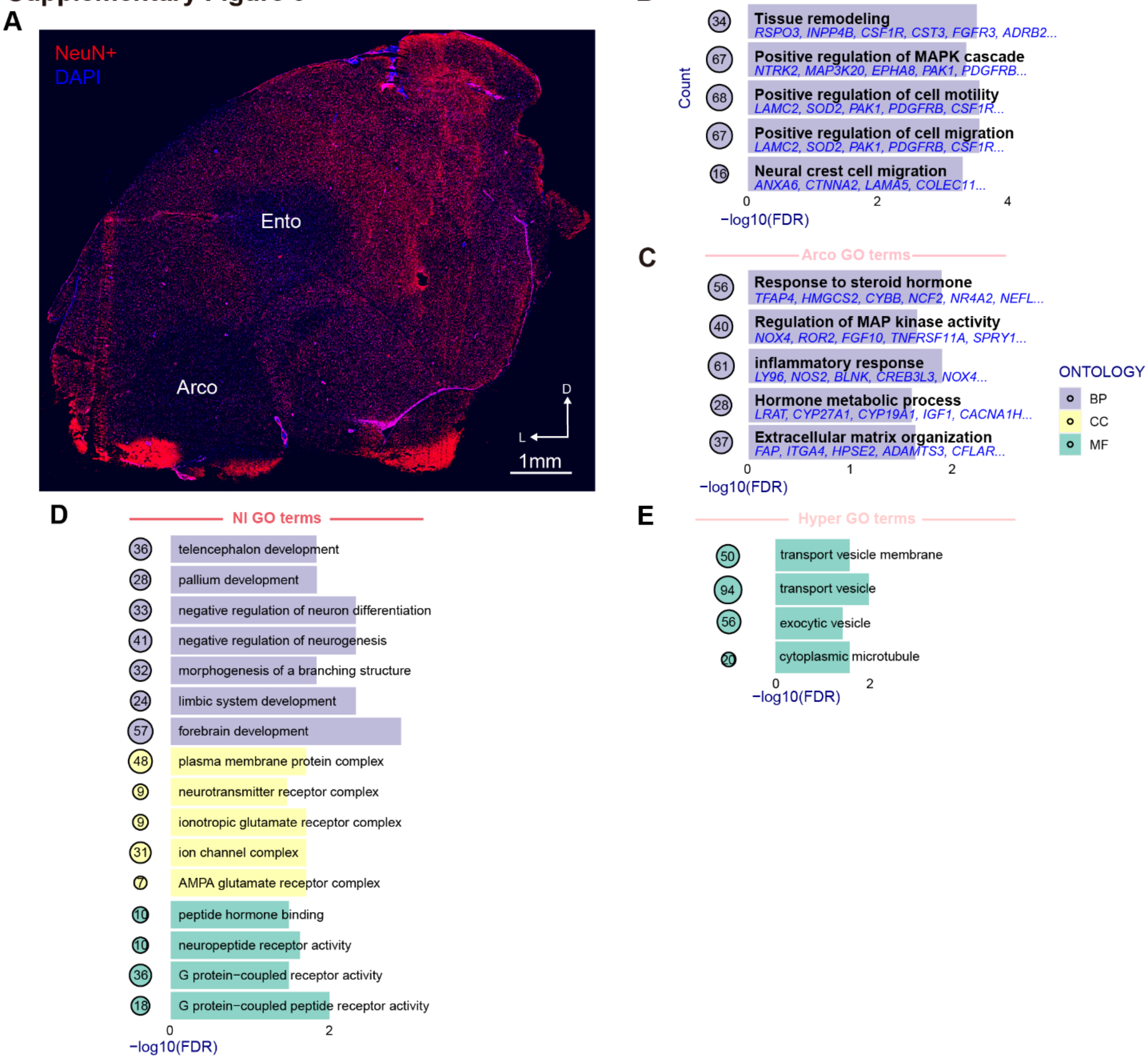


**Fig. S9. Functional specialization of telencephalic subregions in the adult budgerigar brain.** (**A**) NeuN immunostaining of a representative coronal section showing neuron-sparse domains in the entopallium (Ento) and arcopallium (Arco). (**B–E**) Gene Ontology (GO) enrichment analysis of region-specific upregulated genes across major telencephalic subregions: (**B**) Ento; (**C**) Arco; (**D**) NI; (**E**) Hyper. Top enriched GO terms are grouped by ontology domain and plotted by significance (–log$_{10}$FDR). BP, biological process; CC, cellular component; and MF, molecular function.

Supplementary Figure 10

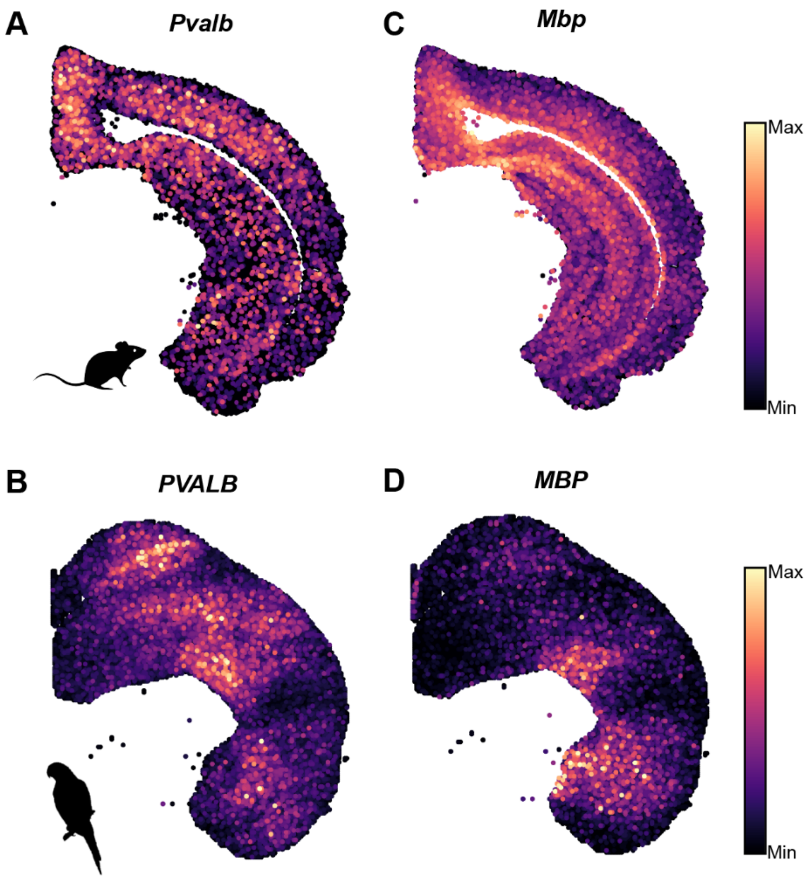


**Fig. S10. Convergent expression patterns in budgerigar entopallium and mouse cortical layer V.**
(**A–D**) Spatial expression of *Pvalb* (**A**) and *Mbp* (**C**) in mouse cortex (*16*), and their orthologs *PVALB* (**B**) and *MBP* (**D**) in budgerigar entopallium.

#### Supplementary Figure 11

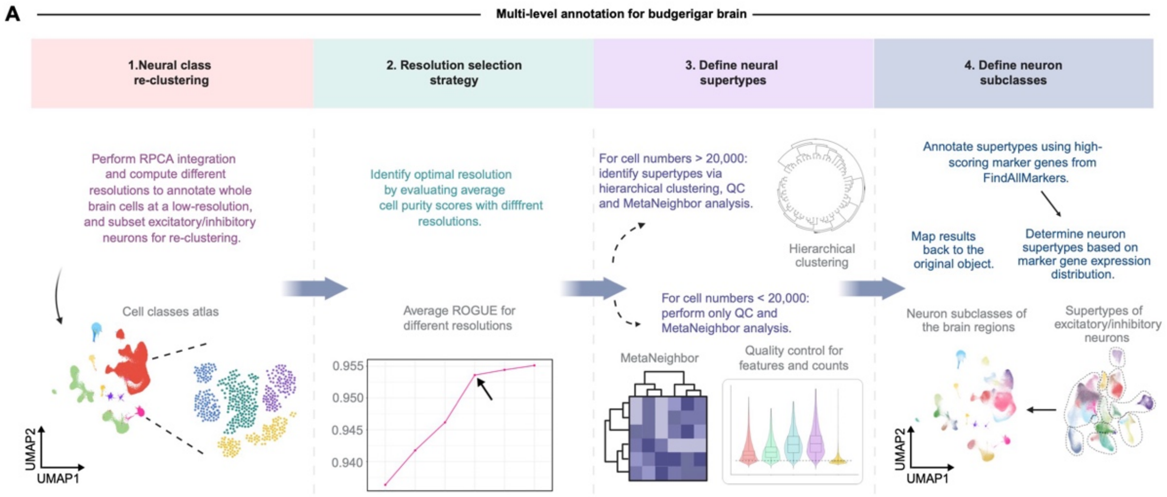


**Fig. S11. Workflow for multi-level neuronal taxonomy in the adult budgerigar brain.**

#### Supplementary Figure 12

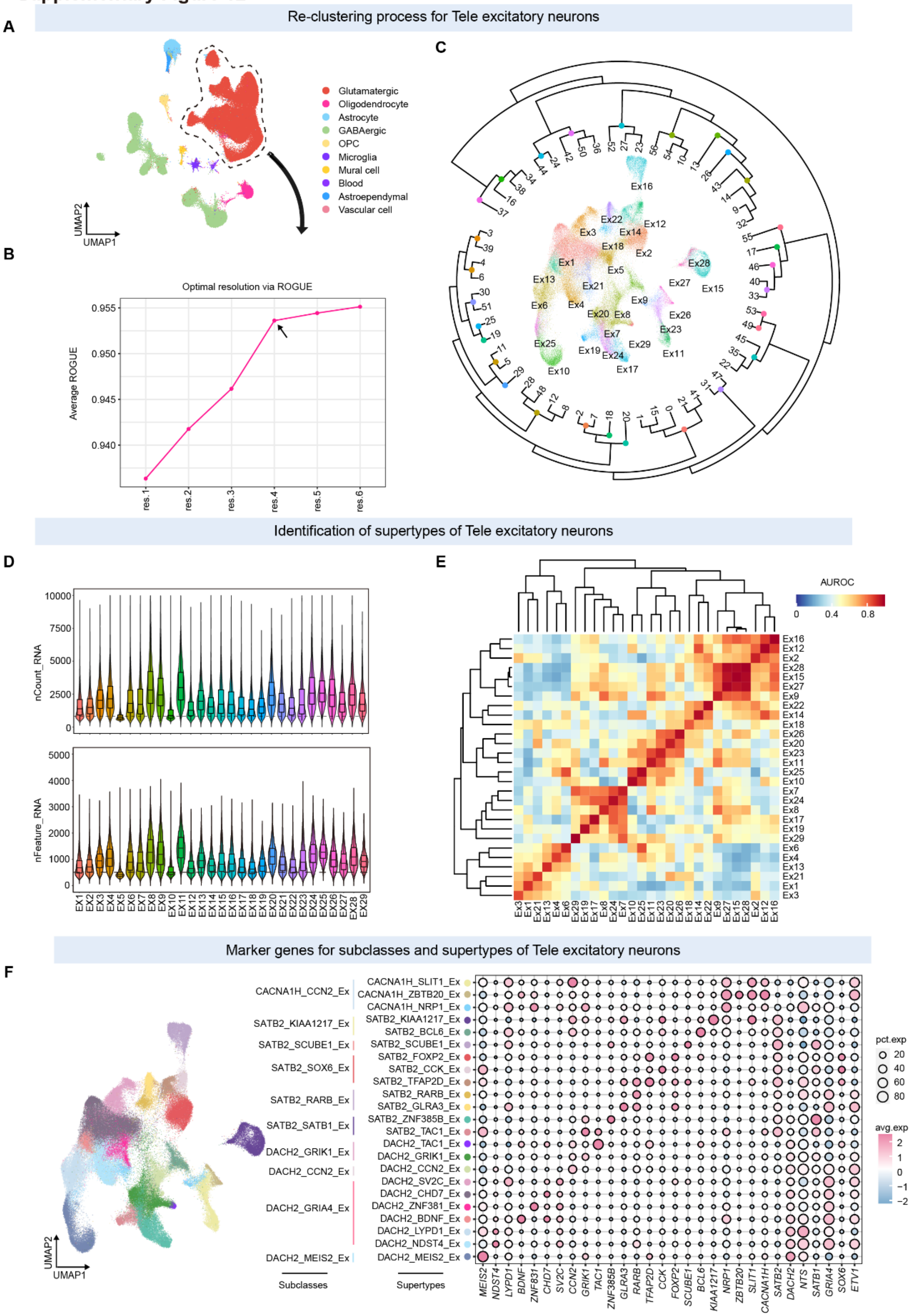

**Fig. S12. Identification and classification of excitatory neuronal supertypes in the budgerigar telencephalon.**

(**A**) UMAP of major cell classes in the telencephalon. Glutamatergic neurons were extracted for re-clustering. (**B**) Determination of optimal clustering resolution based on average ROGUE scores. (**C**) Hierarchical clustering of excitatory neuron subclasses at the optimal resolution, revealing putative supertypes. (**D**) Quality assessment of each cluster via gene and transcript count distributions. (**E**) Supertype-level similarity assessed by MetaNeighbor, confirming transcriptional coherence across excitatory subclasses. (**F**) Final annotation of excitatory neuron supertypes and subclasses. Left: UMAP showing subclass distribution. Right: Dot plot of marker gene expression, highlighting supertype-specific profiles.

**Supplementary Figure 13** Re-clustering process for Tele inhibitory neurons

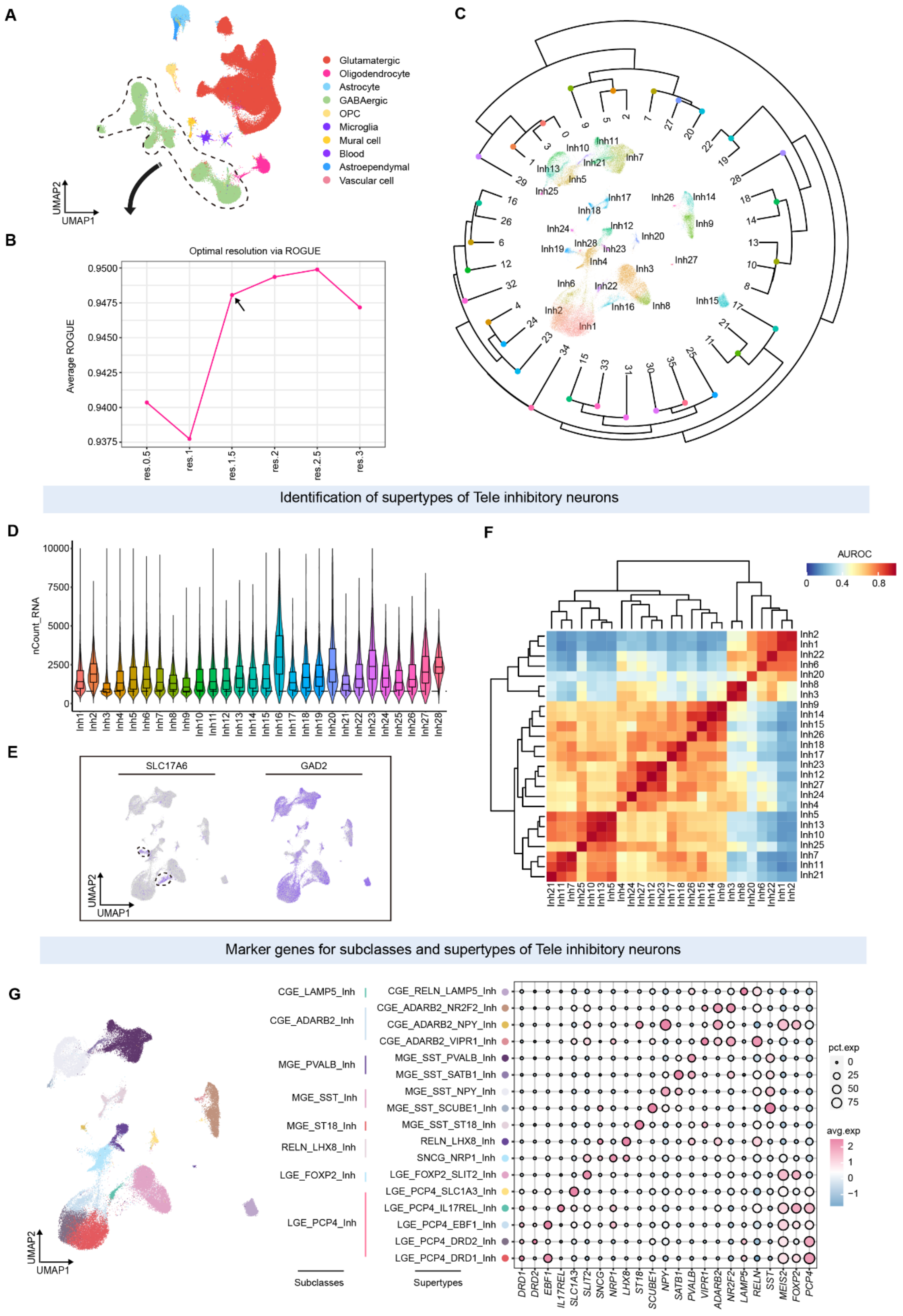

**Fig. S13. Identification and classification of inhibitory neuronal supertypes in the budgerigar telencephalon.**

(**A**) UMAP of major cell classes. GABAergic neurons were selected for inhibitory supertype analysis. (**B**) ROGUE-based evaluation of clustering resolutions to determine optimal purity. (**C**) Hierarchical clustering of inhibitory neuron subclasses at the selected resolution, revealing candidate supertypes. (**D**) Violin plots showing transcript count distributions across putative supertypes. (**E**) UMAP expression plots for *SLC17A6* (glutamatergic marker) and *GAD2* (GABAergic marker) for cell identity validation. (**F**) MetaNeighbor analysis of AUROC-based similarity between supertypes. (**G**) Final classification of inhibitory supertypes and subclasses. Left: UMAP of subclass identities. Right: Dot plot of marker gene expression across supertypes.

#### Supplementary Figure 14

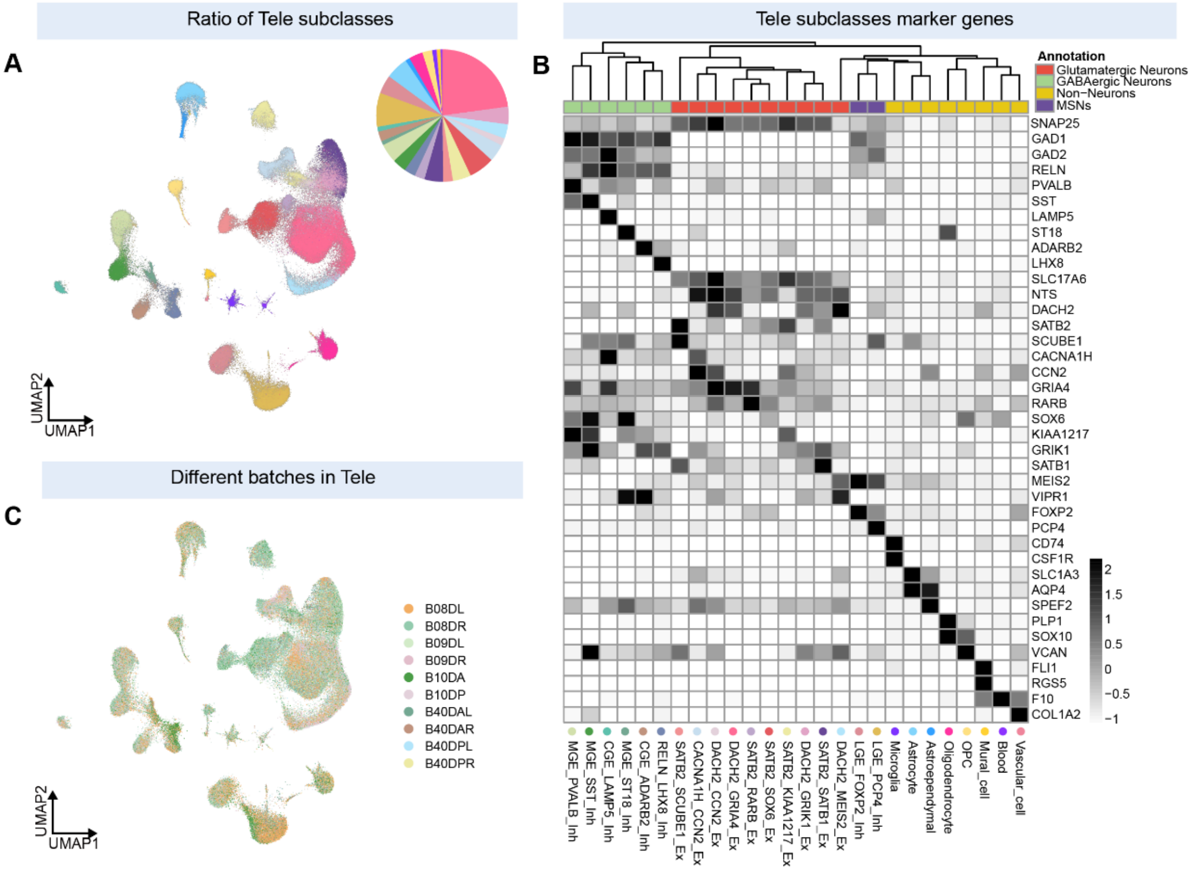


**D** — **Cell taxonomy in Tele**

| Classes | Subclasses | Supertypes | Representative genes: Common genes | Representative genes: Other DEGs | Ratio |
|---|---|---|---|---|---|
| Excitatory neuron | CACNA1H_CCN2_Ex | CACNA1H_SLIT1_Ex | *CACNA1H* | *FGFR2, SEMA3D, KLHL1, CCN2, SLIT1...* | 4.28% |
| | | CACNA1H_ZBTB20_Ex | | *BVES, ST18, MYLK, SLIT2, ZBTB20, PROX1...* | |
| | | CACNA1H_NRP1_Ex | | *NRP1, LHX9, SLIT2, NRP2, SLC1A3...* | |
| | SATB2_KIAA1217_Ex | SATB2_KIAA1217_Ex | *SATB2* | *KIAA1217, IL18R1, IL1R1, IL18RAP, SLIT1...* | 4.46% |
| | | SATB2_BCL6_Ex | | *MYBPC1, BCL6, ADARB2, SLC6A7, SLIT2...* | |
| | SATB2_SCUBE1_Ex | SATB2_SCUBE1_Ex | | *NTNG2, ITPR2, GRIK3, SCUBE1, ZNF385D...* | 2.41% |
| | SATB2_SOX6_Ex | SATB2_FOXP2_Ex | | *TFAP2D, FOXP2, SOX6, DPP10, CADPS2...* | 6.27% |
| | | SATB2_CCK_Ex | | *PRRX1, CEMIP, CADPS2, LMO1, SOX6, CCK...* | |
| | | SATB2_TFAP2D_Ex | | *SMOC2, TFAP2D, SIPA1L2, CADPS2, CDH13...* | |
| | SATB2_RARB_Ex | SATB2_RARB_Ex | | *CNTNAP5, RARB, FSTL4, MEF2C, NTNG1...* | 2.46% |
| | | SATB2_GLRA3_Ex | | *GLRA3, NTNG1, CA2, ETV1, FOXP2, RARB...* | |
| | SATB2_SATB1_Ex | SATB2_ZNF385B_Ex | | *ZNF385B, SULF1, CDH12, RSPO3, SATB1...* | 4.54% |
| | | SATB2_TAC1_Ex | | *CALB1, TAC1, CALD1, MEGF11, SATB1...* | |
| | DACH2_GRIK1_Ex | DACH2_TAC1_Ex | *DACH2, GRIK1* | *TAC1, CEMIP, ALDH1A2, NR2F2, KIF26A...* | 4.30% |
| | | DACH2_GRIK1_Ex | | *CNTN6, GRTP1, PREF1, ALDH1A2, GRIK1...* | |
| | DACH2_CCN2_Ex | DACH2_CCN2_Ex | *DACH2, CCN2* | *TGFB2, PTPRM, EYA1, CDH12, NDST3, CCN2...* | 1.80% |
| | DACH2_GRIA4_Ex | DACH2_SV2C_Ex | *DACH2, GRIA4* | *MMEL1, EYA1, MYBPC3, CALD1, CCN4, SV2C...* | 22.85% |
| | | DACH2_CHD7_Ex | | *CHD7, FBP1, THSD7B, CDH22...* | |
| | | DACH2_ZNF381_Ex | | *ZNF381, CALD1, ITGA6, ADGRB3, SYT10...* | |
| | | DACH2_BDNF_Ex | | *NR4A3, BDNF, NR4A2, AKR1D1...* | |
| | | DACH2_LYPD1_Ex | | *CFTR, MYOM1, VIPR1, NTS...* | |
| | | DACH2_NDST4_Ex | | *NDST4, MYOM1, LRMP, ETV1...* | |
| | DACH2_MEIS2_Ex | DACH2_MEIS2_Ex | *DACH2* | *SEMA5A, RET, KIF26A, VCAN, MEIS2...* | 3.51% |
| Inhibitory neuron | CGE_LAMP5_Inh | CGE_RELN_LAMP5_Inh | *RELN, LAMP5* | *CRTAC1, ANO1, CDH7, IGF1...* | 1.18% |
| | CGE_ADARB2_Inh | CGE_ADARB2_NR2F2_Inh | *RELN, ADARB2* | *CNR1, PROX1, PENK, VIPR1, NR2F2...* | 2.43% |
| | | CGE_ADARB2_NPY_Inh | | *TLL1, GLP1R, PROX1, NPY, CDH13...* | |
| | | CGE_ADARB2_VIPR1_Inh | | *VIPR1, GRIK1, NRP2, VIPR2, CCN4...* | |
| | MGE_PVALB_Inh | MGE_SST_PVALB_Inh | *SST* | *NXPH1, LAMB1, PVALB, SVIL, ETV1...* | 4.63% |
| | | MGE_SST_SATB1_Inh | | *MYBPC3, SATB1, CDH23, GLRA3, ALKAL2...* | |
| | MGE_SST_Inh | MGE_SST_NPY_Inh | | *CGHL1, NPY, RGS6, CDH23, SEMA3D, OTOF...* | 3.40% |
| | | MGE_SST_SCUBE1_Inh | | *CHODL, SFRP2, ITPR2, SCUBE1, CEMIP...* | |
| | MGE_ST18_Inh | MGE_SST_ST18_Inh | | *ST18, TLL1, CHD7, SMOC2, NDST4...* | 1.01% |
| | RELN_LHX8_Inh | RELN_LHX8_Inh | *RELN, LHX8* | *EYA4, EYA1, SLC6A11, MYO1B, SNCG...* | 2.77% |
| | | SNCG_NRP1_Inh | | *ZIC1, NRP1, NRP2, SLIT1...* | |
| | LGE_FOXP2_Inh | LGE_FOXP2_SLIT2_Inh | *MEIS2, FOXP2* | *CADPS2, SLIT2, NTNG1, FOXP2, SULF1...* | 4.53% |
| | LGE_PCP4_Inh | LGE_PCP4_SLC1A3_Inh | *MEIS2, PCP4* | *EGF, SLC13A5, RFX4, CA8, SLC1A3...* | 8.29% |
| | | LGE_PCP4_IL17REL_Inh | | *IL17REL, KCNH8, NTNG1, FOXP2, PROM1...* | |
| | | LGE_PCP4_EBF1_Inh | | *EBF1, ISL1, TAC1, BVES, PENK.SGCG...* | |
| | | LGE_PCP4_DRD2_Inh | | *PDE10A, RGS12, ZFHX4...* | |
| | | LGE_PCP4_DRD1_Inh | | *IAPP, EBF1, RASGRP1, MYOM1...* | |

**Fig. S14. Cell type taxonomy of the budgerigar telencephalon.**
(**A**) UMAP projection of snRNA-seq data from the telencephalon, with cells colored by subclass identity. Pie chart indicates the proportional representation of each subclass. (**B**) Dot plot showing expression of marker genes across telencephalic subclasses. Color indicates scaled expression; dot size reflects the proportion of expressing cells. (**C**) UMAP colored by batch labels, showing no obvious batch effect among sampled brains. (**D**) Summary table of telencephalic neuronal taxonomy, including subclass and supertype annotations, shared and representative marker genes, and subclass proportion.

#### Supplementary Figure 15

Re-clustering process for Mid excitatory neurons

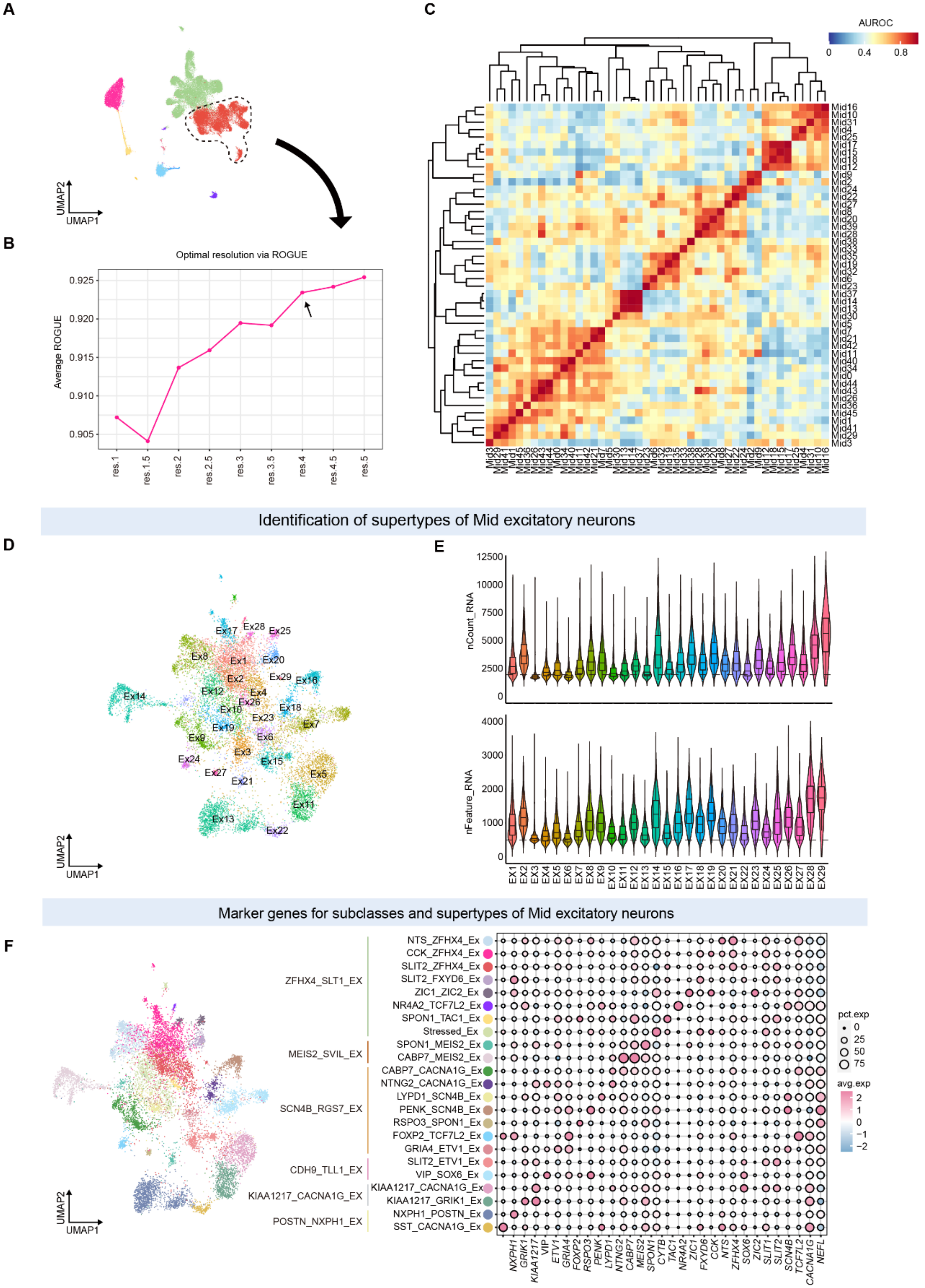

**Fig. S15. Identification of excitatory neuronal supertypes in the budgerigar midbrain.**
(**A**) UMAP of midbrain cell classes. Glutamatergic neurons were selected for downstream re-clustering. (**B**) ROGUE-based evaluation of clustering resolution to determine optimal subclass separation. (**C**) Hierarchical clustering of excitatory neurons at optimal resolution, identifying transcriptionally distinct supertypes. (**D**) UMAP projection of excitatory neuron supertypes following re-clustering. (**E**) Violin plots showing transcript count and feature number distributions across clusters. (**F**) Final annotation of excitatory supertypes and subclasses. Left: UMAP of subclass distribution. Right: Dot plot showing marker gene expression profiles across supertypes.

**Supplementary Figure 16**

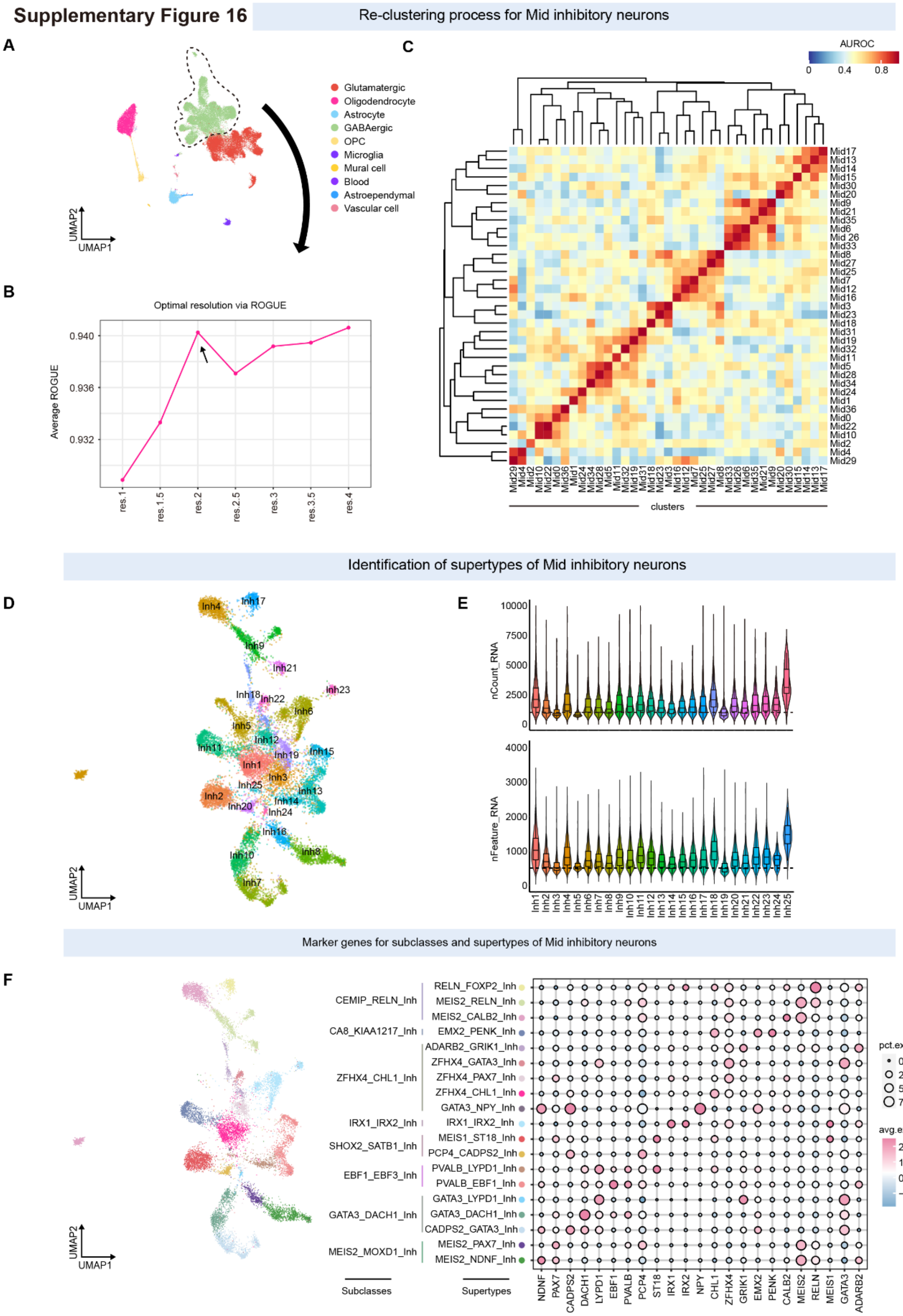

**Fig. S16. Identification of inhibitory neuronal supertypes in the budgerigar midbrain.**
(**A**) UMAP of major midbrain cell classes. GABAergic neurons were extracted for downstream analysis. (**B**) ROGUE-based evaluation of clustering resolution to determine optimal subclass separation. (**C**) Hierarchical clustering of inhibitory neuron clusters at the optimal resolution, revealing candidate supertypes. (**D**) UMAP projection of inhibitory neuronal supertypes. (**E**) Violin plots showing transcript and gene count distributions across putative supertypes. (**F**) Final annotation of midbrain inhibitory neuronal subclasses and supertypes. Left: UMAP colored by subclass identity. Right: Dot plot showing expression of representative marker genes across supertypes.

#### Supplementary Figure 17

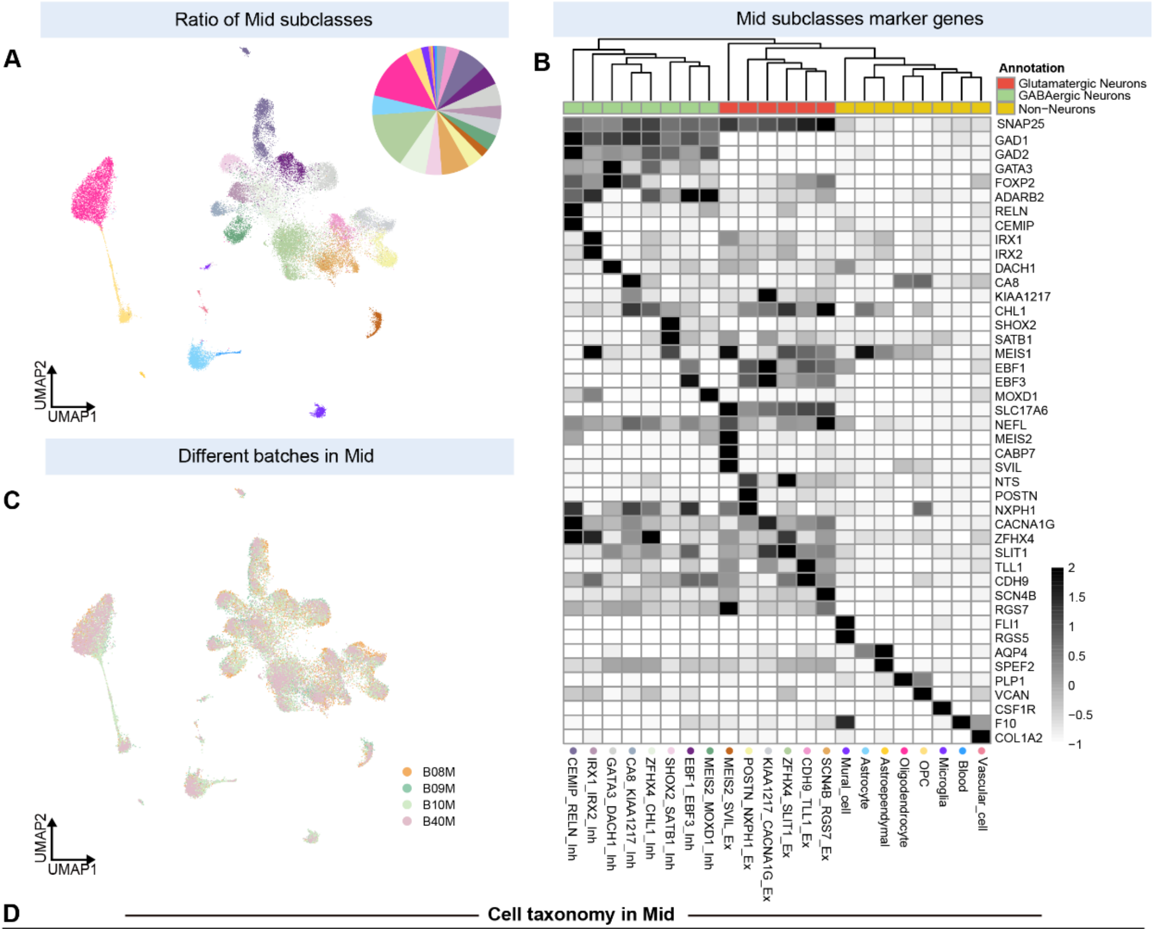


**D** — **Cell taxonomy in Mid**

| Classes | Subclasses | Supertypes | Representative genes: Common genes | Representative genes: Other DEGs | Ratio |
|---|---|---|---|---|---|
| Excitatory neuron | ZFHX4_SLIT1_Ex | NTS_ZFHX4_Ex | ZFHX4 | NTS, NPNT, SYT10, TCF7L2, SFRP2... | 14.19% |
| | | CCK_ZFHX4_Ex | | VCAN, DSCAM, CCK, VAT1L... | |
| | | SLIT2_ZFHX4_Ex | | TNNT3, DSCAM, KLHL1, ARGLU1, SLIT2... | |
| | | SLIT2_FXYD6_Ex | | CHD7, NFIA, NFIB, NXPH1, DACH1, FXYD6... | |
| | | ZIC1_ZIC2_Ex | | ZIC1, ZIC2, COL13A1, CHD7, ST18... | |
| | | NR4A2_TCF7L2_Ex | | NR4A2, TEC, IL1R1, CALB2... | |
| | | SPON1_TAC1_Ex | | GLI3, TAC1, RSPO3, MEIS2, MEIS1... | |
| | | Stressed_Ex | | ND3, COX3, COX1, CYTB... | |
| | | SPON1_MEIS2_Ex | | SPON1, PTPRM, MEIS1, CHL1, IRX1... | |
| | MEIS2_SVIL_Ex | CABP7_MEIS2_Ex | MEIS2 | BVES, SVIL, RGS2, CABP7, GADL1... | 2.13% |
| | SCN4B_RGS7_Ex | CABP7_CACNA1G_Ex | SCN4B | CA2, SNTG2, TCF7L2, RGS7, CABP7... | 6.91% |
| | | NTNG2_CACNA1G_Ex | | MDGA1, VIP, NTNG2, NDNF, SEMA3D... | |
| | | LYPD1_SCN4B_Ex | | STC2, SMOC2, SYT2, LYPD1, CALD1, PTPRC... | |
| | | PENK_SCN4B_Ex | | FREM1, COL6A3, PENK, FOXP2, NEFL... | |
| | | RSPO3_SPON1_Ex | | MYBPC1, NFIB, CNTN6, NEUROD6, RSPO3... | |
| | | FOXP2_TCF7L2_Ex | | TCF7L2, FOXP2, CXCL12, CACNA2D1, CA8... | |
| | | GRIA4_ETV1_Ex | | ETV1, OTOF, SYT2, FOXP2, POSTN, GRIA4... | |
| | CDH9_TLL1_Ex | SLIT2_ETV1_Ex | CDH9, TLL1 | ETV1, MPP7, ITGA8, BCL11B, TFAP2D, SLIT2... | 3.42% |
| | | VIP_SOX6_Ex | | VIP, PENK, MEGF11, SOSTDC1, SOX6... | |
| | KIAA1217_CACNA1G_Ex | KIAA1217_CACNA1G_Ex | KIAA1217, CACNA1G | PREX2, BNC2, SOX6, ZEB2... | 4.29% |
| | | KIAA1217_GRIK1_Ex | | FXN, NR5A2, ZEB2 ,GRIK1... | |
| | POSTN_NXPH1_Ex | NXPH1_POSTN_Ex | POSTN, NXPH1 | SHOX, GFRA1, MEF2C, CDH7... | 4.16% |
| | | SST_CACNA1G_Ex | | SST, PIEZO2, NTF3, NRP2, CACNA1G... | |
| Inhibitory neuron | CEMIP_RELN_Inh | RELN_FOXP2_Inh | CEMIP, RELN | FOXP2, SEMA3A, SEMA3C, FOXP4, GRIA1... | 7.46% |
| | | MEIS2_RELN_Inh | | ADCYAP1, MEIS2, MEF2C, CDH7, RARB... | |
| | | MEIS2_CALB2_Inh | | SEMA3A, SLC2A42, TRPC7, CACNA1G, NRP1... | |
| | CA8_KIAA1217_Inh | EMX2_PENK_Inh | PENK | EMX2, CA8, KIAA1217, PTPRK, NTNG1... | 2.42% |
| | ZFHX4_CHL1_Inh | ADARB2_GRIK1_Inh | ZFHX4 | KCNH8, ADARB2, SYT10, CDH7, GRIK1... | 6.81% |
| | | ZFHX4_GATA3_Inh | | GATA3, PTHLH, RSPO3, SEMA5A, KCNT2... | |
| | | ZFHX4_PAX7_Inh | | TLL1, CDH13, PCDH17, PAX7... | |
| | | ZFHX4_CHL1_Inh | | CHL1, NR2F2, STMN2, FXYD6, SNCG... | |
| | | GATA3_NPY_Inh | | NPY, SYT10, SGCG, DLK1... | |
| | IRX1_IRX2_Inh | IRX1_IRX2_Inh | IRX1, IRX2 | CHODL, NTNG1, TAC1, GRIK2, MEIS1... | 3.38% |
| | SHOX2_SATB1_Inh | MEIS1_ST18_Inh | SHOX2, SATB1 | ST18, RSPO3, TAFA2, CACNA1E... | 4.12% |
| | | PCP4_CADPS2_Inh | | RUNX2, LMO1, NFIB, CADPS2, PCP4... | |
| | EBF1_EBF3_Inh | PVALB_LYPD1_Inh | PVALB | COL13A1, ST18, LYPD1, EBF3, CHD7... | 4.92% |
| | | PVALB_EBF1_Inh | | EBF1, EBF3, PCDH18, GLI3... | |
| | GATA3_DACH1_Inh | GATA3_LYPD1_Inh | GATA3 | LYPD1, SOX21, GRIK1, PTPRK, SYT10... | 5.55% |
| | | GATA3_DACH1_Inh | | DACH1, PAX3, GLI3, PTPRT, CDH11, DACH2... | |
| | | CADPS2_GATA3_Inh | | ZEB2, GATA2, ANOS1, FOXP2, PTPRD... | |
| | MEIS2_MOXD1_Inh | MEIS2_PAX7_Inh | MEIS2 | ADGRG2, LMO1, CDH9, PAX7... | 4.08% |
| | | MEIS2_NDNF_Inh | | MOXD1, NDNF, PLCH1, KIT, GRIK2... | |

**Fig. S17. Cell taxonomy of the adult budgerigar midbrain.**
(**A**) UMAP projection of midbrain snRNA-seq data colored by neuronal subclasses. The pie chart indicates the proportional representation of each subclass. (**B**) Heatmap showing the expression of subclass-specific marker genes across midbrain neuronal types. Genes are grouped by functional annotation; dot intensity reflects scaled expression. (**C)** UMAP projection colored by sample batch identity, showing consistent subclass structure across animals. (**D**) Summary table of excitatory and inhibitory neuronal subclasses and supertypes in the midbrain. Common marker genes and representative DEGs are listed for each subclass. The final column indicates the proportion of each subclass within the midbrain dataset.

#### Supplementary Figure 18

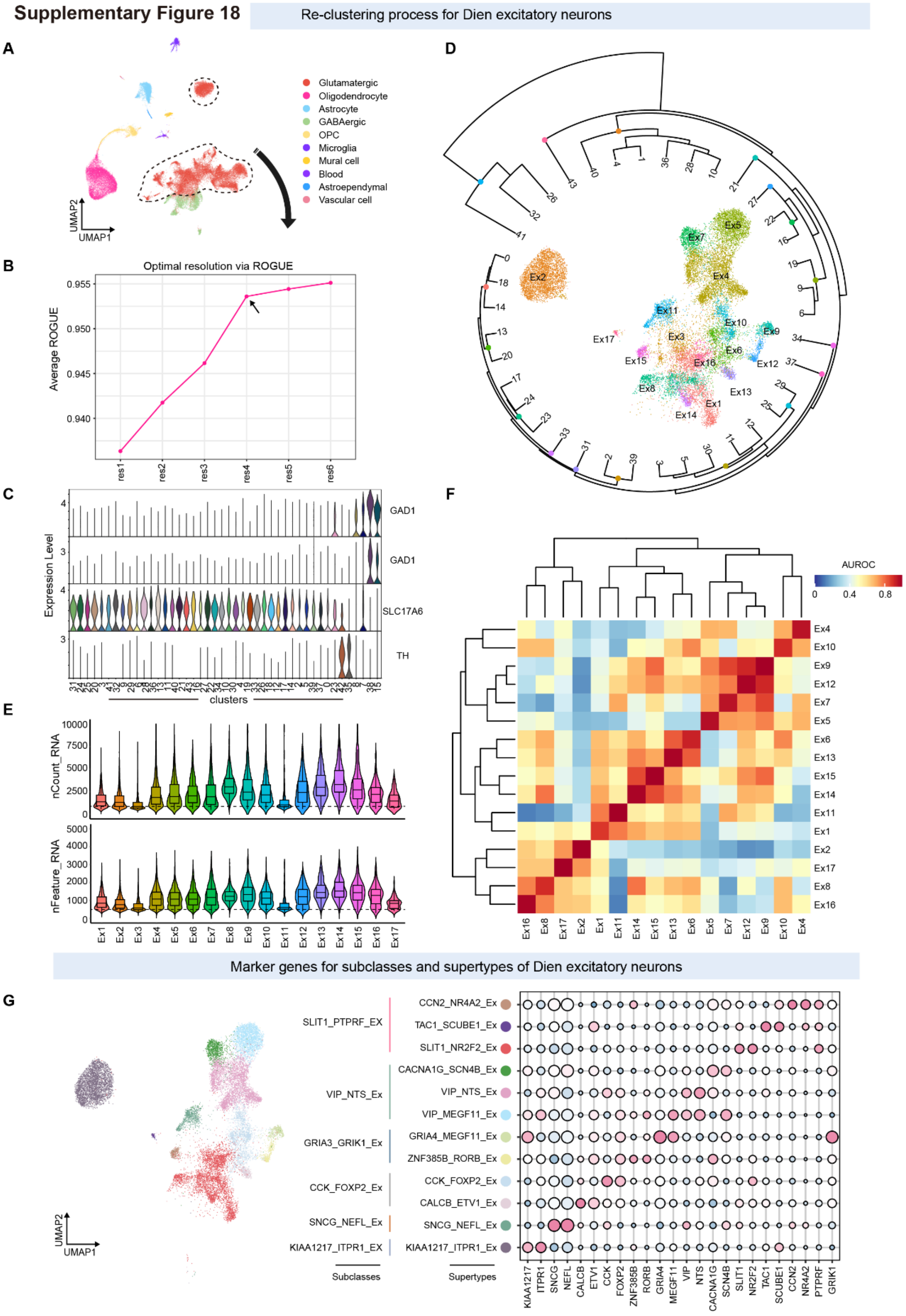

**Fig. S18. Re-clustering of diencephalic excitatory neuronal supertypes.**
(**A**) Cellular composition classification of budgerigar diencephalon. The glutamatergic cluster was selected to be re-clustered into excitatory neuronal supertypes. (**B**) Assessment of ROGUE-based cellular purity scores across clustering resolutions, with optimal resolution determined at the inflection point for supertype identification. (**C**) Expression of glutamatergic neuronal makers and GABAergic neuronal markers across potential supertypes. (**D**) Identification of potential excitatory supertypes through hierarchical clustering at the optimal resolution from (**B**). (**E**) Violin plots display feature distribution and count metrics across putative supertypes. (**F**) Supertype consolidation through Metaneighbor similarity analysis. (**G**) Final classification of diencephalic excitatory neuronal supertypes with subclass annotations. (Left) UMAP visualization of diencephalic excitatory neuronal supertypes. (Right) Dot plot representation of supertype-specific marker expression patterns.

#### Supplementary Figure 19

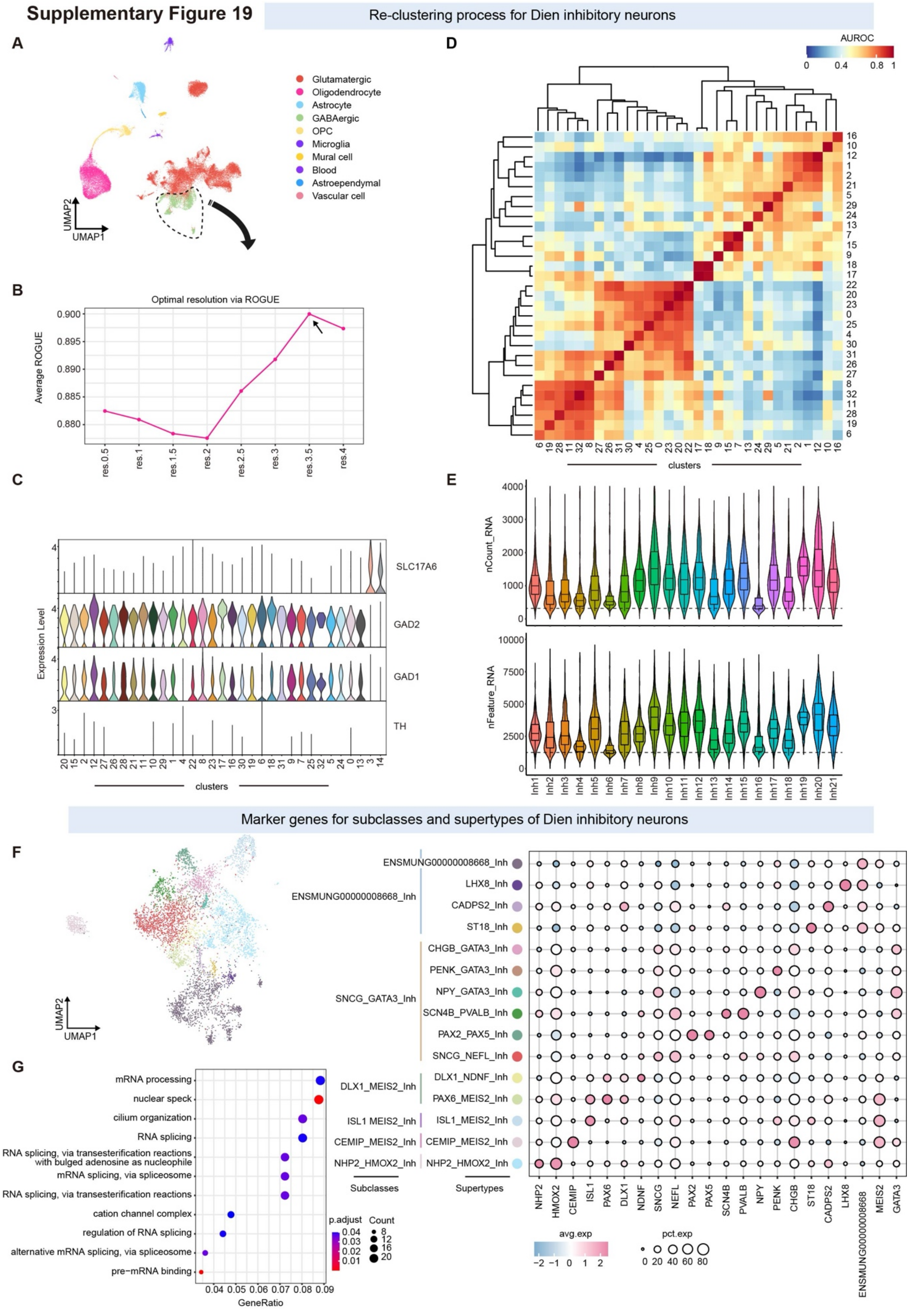

**Fig. S19. Identification of inhibitory neuronal supertypes in the budgerigar diencephalon.** (**A**) UMAP of major diencephalic cell classes. GABAergic neurons were selected for inhibitory supertype re-clustering. (**B**) ROGUE score evaluation to determine optimal clustering resolution for subtype purity. (**C**) Expression profiles of GABAergic (*GAD1*, *GAD2*), glutamatergic (*SLC17A6*), and dopaminergic (*TH*) markers across clusters. (**D**) Supertype similarity matrix derived from MetaNeighbor AUROC analysis. (**E**) Violin plots of RNA counts and gene feature distributions per cluster. (**F**) Final classification of inhibitory supertypes and subclasses. Left: UMAP of subclass identities. Right: Dot plot of representative marker gene expression across supertypes. (**G**) GO term enrichment for *ENSMUNG00000008668_Inh*, indicating functional specialization in mRNA processing and RNA splicing.

#### Supplementary Figure 20

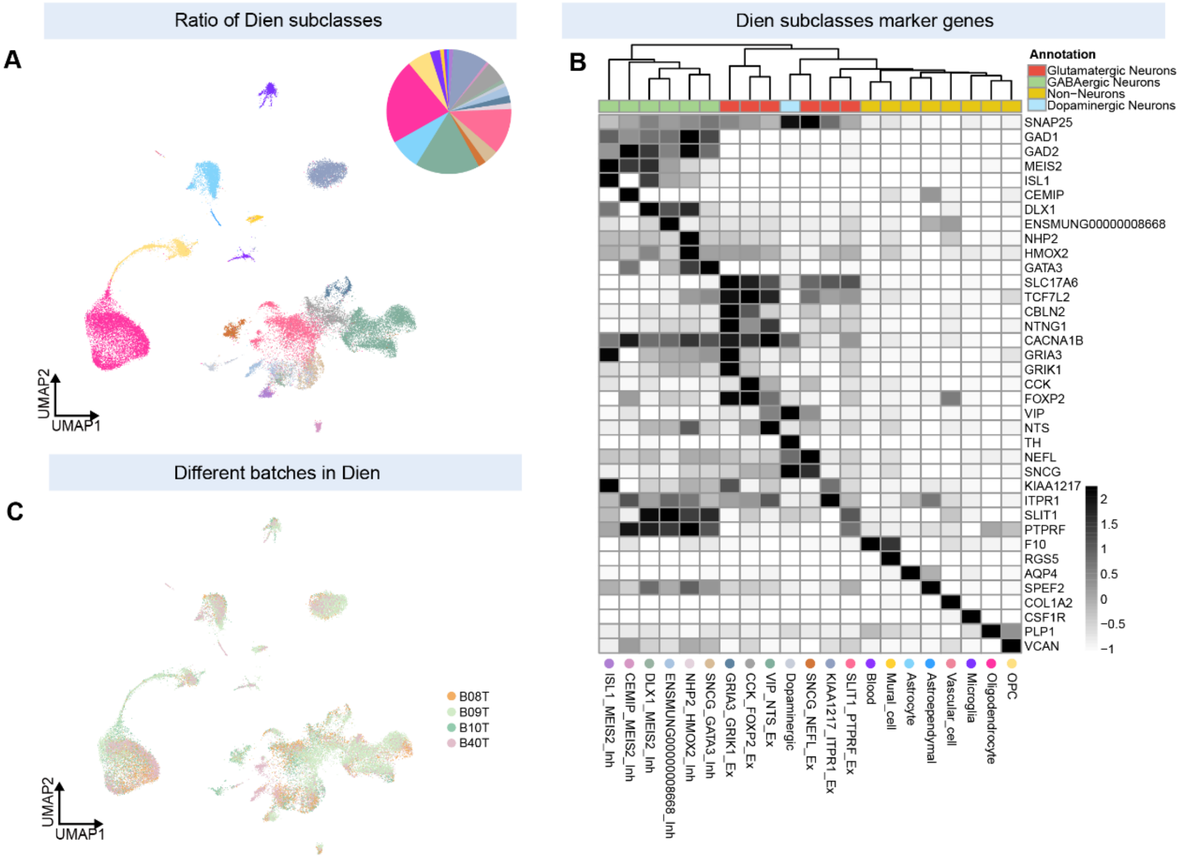


**D** Cell taxonomy in Dien

| Classes | Subclasses | Supertypes | Representative genes: Common genes | Representative genes: Other DEGs | Ratio |
|---|---|---|---|---|---|
| Excitatory neuron | SLIT1_PTPRF_EX | CCN2_NR4A2_Ex | *SLIT1, PTPRF* | *TFAP2D, NR4A2, IL1R1, KCNQ5, CCN2...* | 11.95% |
| | | TAC1_SCUBE1_Ex | | *KCTD8, CXCL14, SYT9, IRX1, NR4A3...* | |
| | | SLIT1_NR2F2_Ex | | *NR2F2, TAFA2, PCDH17, TRPC4, CDH4...* | |
| | VIP_NTS_Ex | CACNA1G_SCN4B_Ex | *VIP* | *SCN4B, BVES, RGS3, CDH13, PTPRD, CACNA1G...* | 16.76% |
| | | VIP_NTS_Ex | | *NTS, PTPRC, NPTX2, CACNA1B, NR2F1...* | |
| | | VIP_MEGF11_Ex | | *SCN4B, MYO1B, MEGF11, VIPR2, PTPRD, NTNG1...* | |
| | GRIA3_GRIK1_Ex | GRIA4_MEGF11_Ex | *GRIA3, GRIK1* | *CA2, PVALB, GPR157, GRIA4, MEGF11...* | 1.96% |
| | | ZNF385B_RORB_Ex | | *ACTN2, FGFR2, CA2, DLK1, RORB, ZNF385B...* | |
| | CCK_FOXP2_Ex | CCK_FOXP2_Ex | *CCK, FOXP2* | *SEMA3C, CBLN2, PIEZO2, ZFHX4...* | 5.25% |
| | | CALCB_ETV1_Ex | | *CALCB, TMEM45A, GBX2, SEMA3A, ETV1...* | |
| | SNCG_NEFL_Ex | SNCG_NEFL_Ex | *NEFL* | *HSPA8, GAPDH, CALM2, TUBB4B, CALM1, SCNG...* | 2.02% |
| | KIAA1217_ITPR1_EX | KIAA1217_ITPR1_Ex | *KIAA1217* | *ST18, NFIB, SULF1, CDH23, CDH7, ITPR1...* | 9.23% |
| Inhibitory neuron | ENSMUNG00000008668_Inh | ENSMUNG00000008668_Inh | *ENSMUNG00000008668* | *ENSMUNG00000011795, SRRM1, CACNA1E...* | 2.42% |
| | | LHX8_Inh | | *LHX8, SLC10A4, SLC5A7...* | |
| | | CADPS2_Inh | | *CNTN6, MEOX2, CADPS2...* | |
| | | ST18_Inh | | *ST18, MIR181A1, MIR181B1...* | |
| | SNCG_GATA3_Inh | CHGB_GATA3_Inh | *GATA3* | *GATA2, CD74, TCF7L2...* | 3.75% |
| | | PENK_GATA3_Inh | | *GATA2, LMO1, PCP4, NXPH1, PENK...* | |
| | | NPY_GATA3_Inh | | *NPY, CD36, SOX6, NDST4, TCF7L2, CCN4...* | |
| | | SCN4B_PVALB_Inh | | *PVALB, SCN4B, SYT2, ADARB2, CA2, TCF7L2...* | |
| | | PAX2_PAX5_Inh | | *PAX2, PAX5, NXPH1, OTOF, SULF2, ZNF536...* | |
| | | SNCG_NEFL_Inh | | *COX1.ND6, HSPA8, COX3, GAPDH...* | |
| | DLX1_MEIS2_Inh | DLX1_NDNF_Inh | *MEIS2* | *PAX6, NDNF, CDH8, NR2F1, DLX1...* | 1.05% |
| | | PAX6_MEIS2_Inh | | *PAX6, CDH23, GFRA1, ISL1, DACH1...* | |
| | ISL1 MEIS2_Inh | ISL1_MEIS2_Inh | | *PTPRD, ISL1, KIAA1217, GRIA3, CHL1...* | 1.15% |
| | CEMIP_MEIS2_Inh | CEMIP_MEIS2_Inh | | *CEMIP, SVEP1, SEMA3E, NXPH2...* | 0.74% |
| | NHP2_HMOX2_Inh | NHP2_HMOX2_Inh | *NHP2* | *HMOX2, DSCAM, DND1, ADGRB3...* | 1.56% |

**Fig. S20. Cell taxonomy of the budgerigar diencephalon.**
(**A**) UMAP projection of snRNA-seq data from the diencephalon, with cells colored by neuronal subclass. The pie chart shows the proportional abundance of each subclass. (**B**) Heatmap of subclass-specific marker gene expression across diencephalic neurons. Gene annotations include glutamatergic, GABAergic, non-neuronal, and dopaminergic markers. (**C**) UMAP projection colored by batch label, indicating high consistency across biological replicates. (**D**) Summary table of neuronal subclasses and supertypes in the diencephalon, with shared marker genes, representative DEGs, and subclass proportions.

**Supplementary Figure 21** Re-clustering process for Cere excitatory and inhibitory neurons

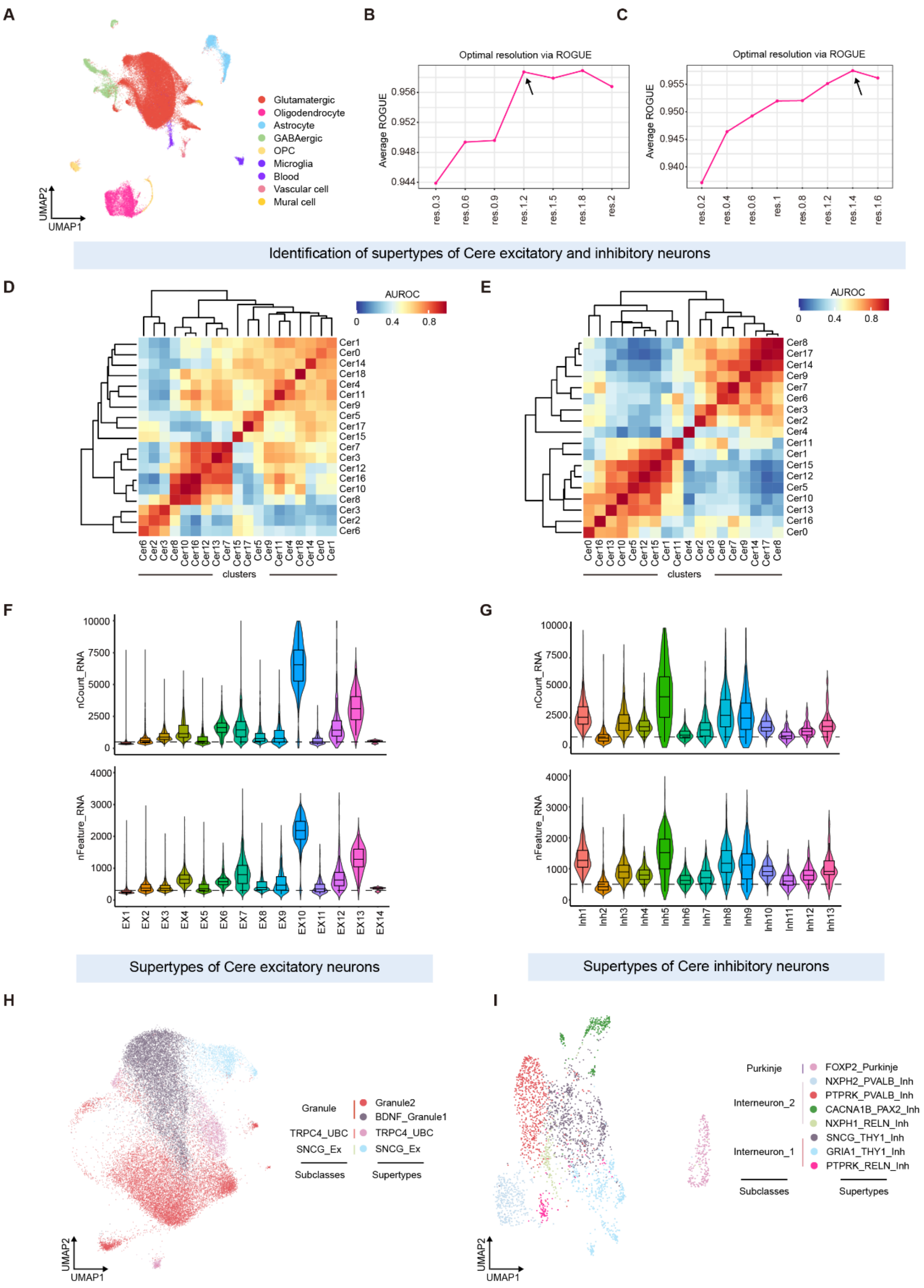

**Fig. S21. Identification of excitatory and inhibitory neuronal supertypes in the budgerigar cerebellum.**

(**A**) UMAP projection of cerebellar cells, colored by major class identity. Glutamatergic and GABAergic neurons were selected for re-clustering. (**B, C**) ROGUE-based evaluation of clustering resolutions for excitatory (**B**) and inhibitory (**C**) neurons to determine optimal subtype separation. (**D, E**) MetaNeighbor AUROC similarity matrices for excitatory (**D**) and inhibitory (**E**) clusters, identifying coherent supertypes. (**F, G**) Violin plots showing transcript counts and gene feature distributions across excitatory (**F**) and inhibitory (**G**) neuronal supertypes. (**H, I**) Final classification of cerebellar excitatory (**H**) and inhibitory (**I**) neuronal supertypes and subclasses. Left panels show UMAP projections colored by subclass identity; right panels annotate supertypes with representative subclass markers.

Supplementary Figure 22

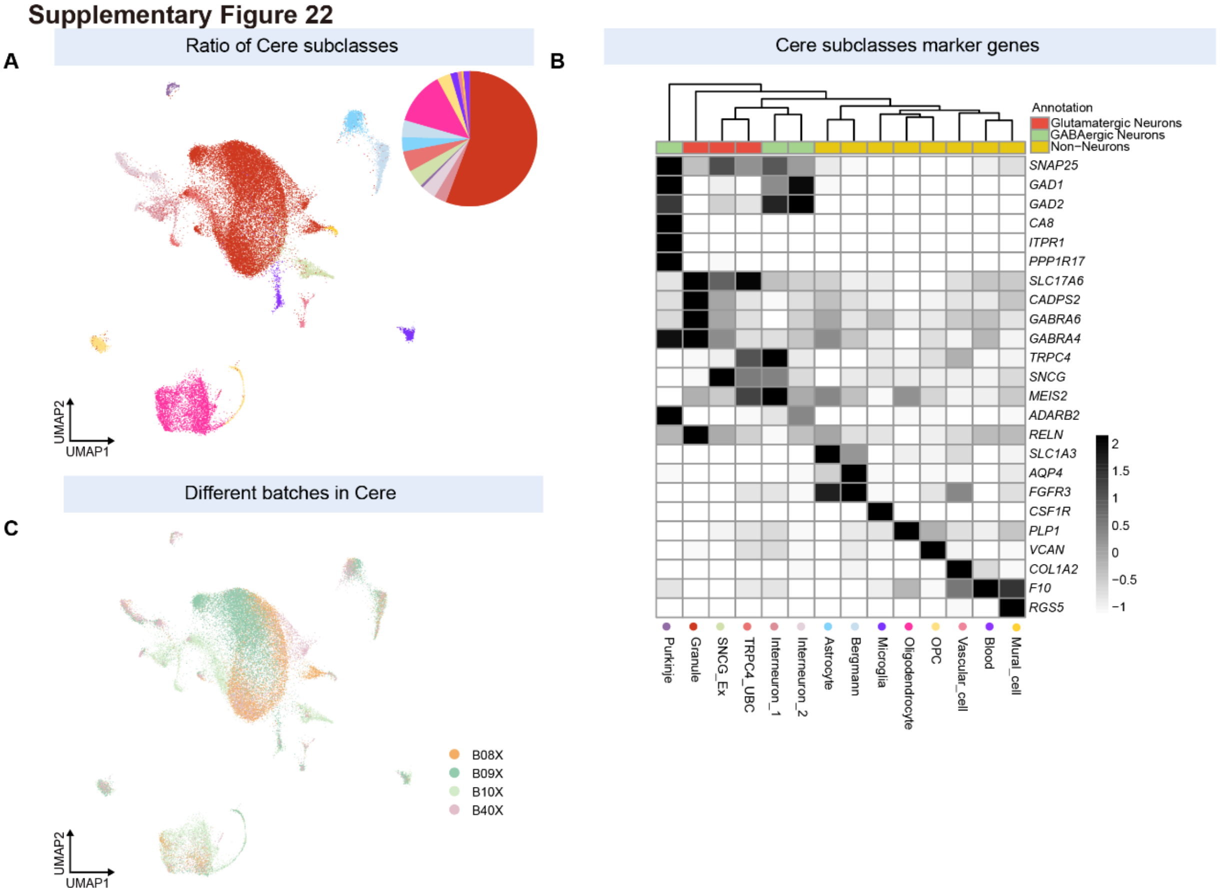


**Fig. S22. Cell taxonomy of the budgerigar cerebellum.**
(**A**) UMAP projection of cerebellar snRNA-seq data with subclass-level annotation. The pie chart shows proportional representation of each subclass, including granule cells, Purkinje neurons, unipolar brush cells (UBC), interneurons, and glial types. (**B**) Heatmap showing expression of subclass-specific marker genes. Annotation categories include glutamatergic neurons, GABAergic neurons, and non-neuronal cell types. (**C**) UMAP colored by biological replicate (batch), indicating consistent subclass structure across individuals.

**Supplementary Figure 23**

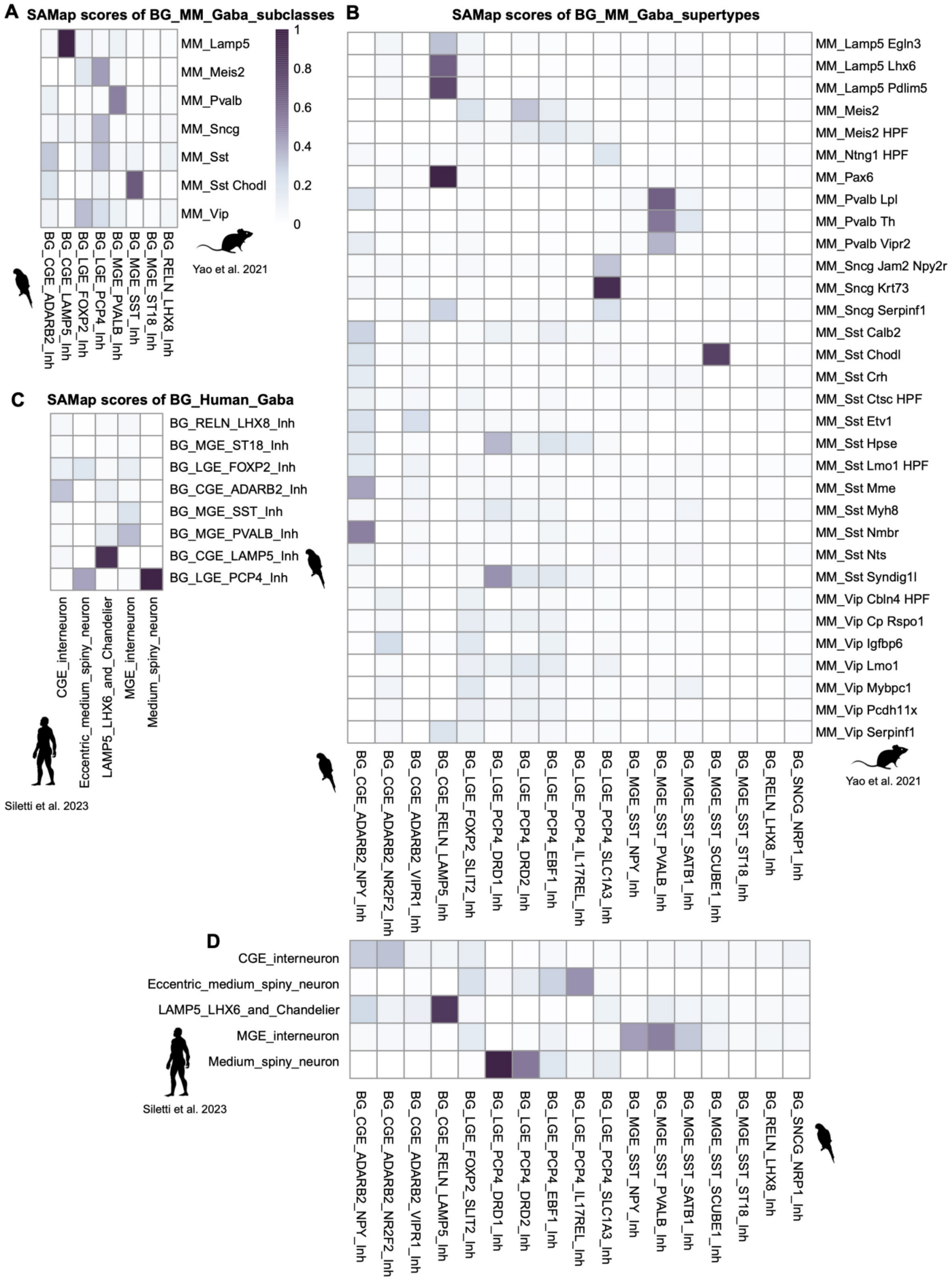

**Fig. S23. Cross-species comparison of inhibitory neuron types between budgerigar telencephalon and mammalian cortex.**

(**A, B**) SAMap alignment of inhibitory neurons between budgerigar and mouse (*12*), at the subclass (**A**) and supertype (**B**) levels. Heatmaps show cross-species transcriptomic similarity scores between annotated GABAergic populations. (**C, D**) SAMap comparison between budgerigar and human (*17*) inhibitory neurons, at the subclass (**C**) and supertype (**D**) levels.

#### Supplementary Figure 24

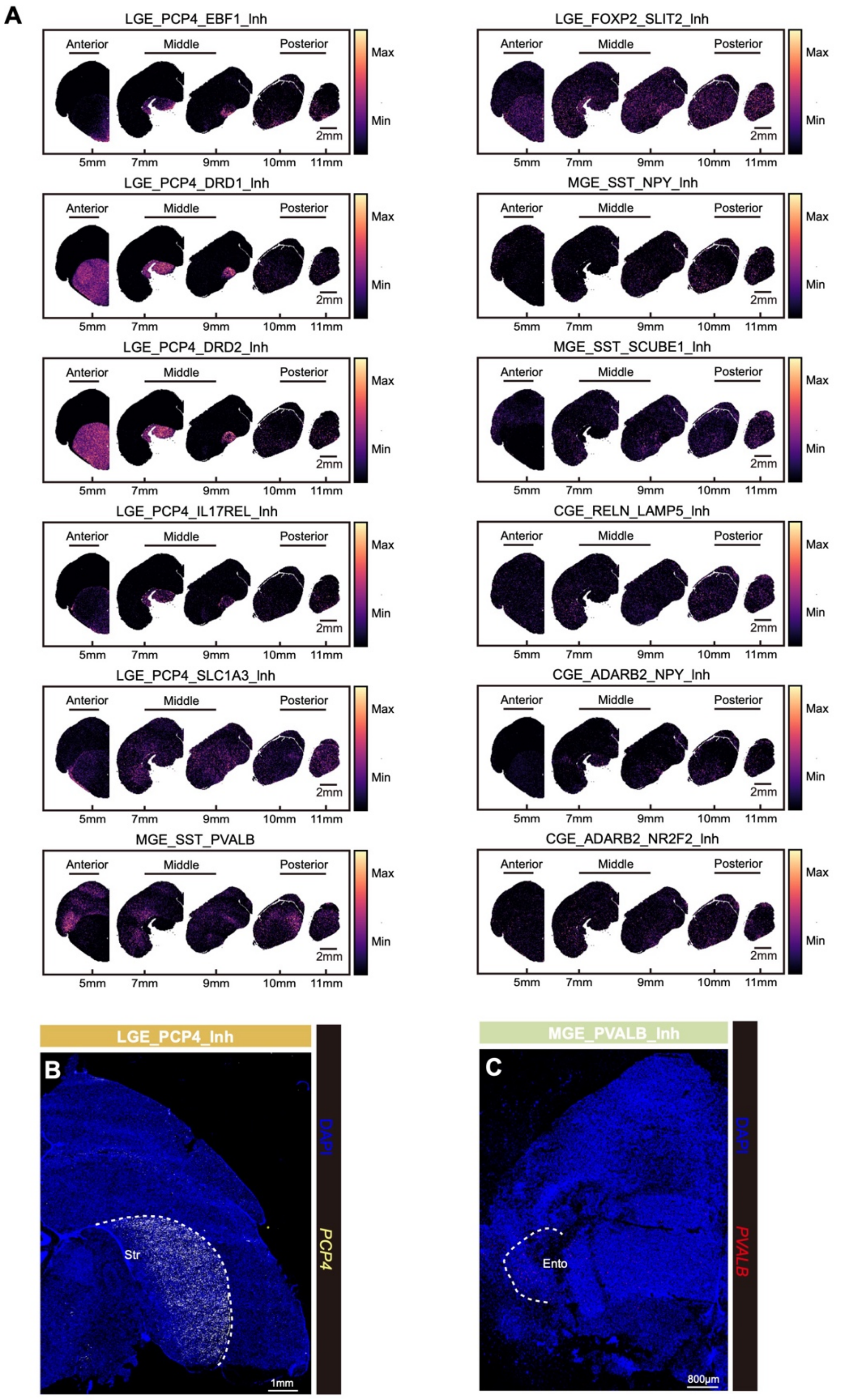

**Fig. S24. Spatial distribution of telencephalic inhibitory neuronal supertypes.**
(**A**) Spatial expression maps of 12 inhibitory supertypes across anterior, middle, and posterior telencephalic sections, inferred by RCTD deconvolution. Each map shows relative expression intensity along the anterior–posterior axis. (**B, C**) Experimental validation by HCR RNA-FISH. Fluorescent in situ hybridization confirms spatial enrichment of *PCP4* (**B**, striatum) and *PVALB* (**C**, entopallium) at single-cell resolution.

#### Supplementary Figure 25

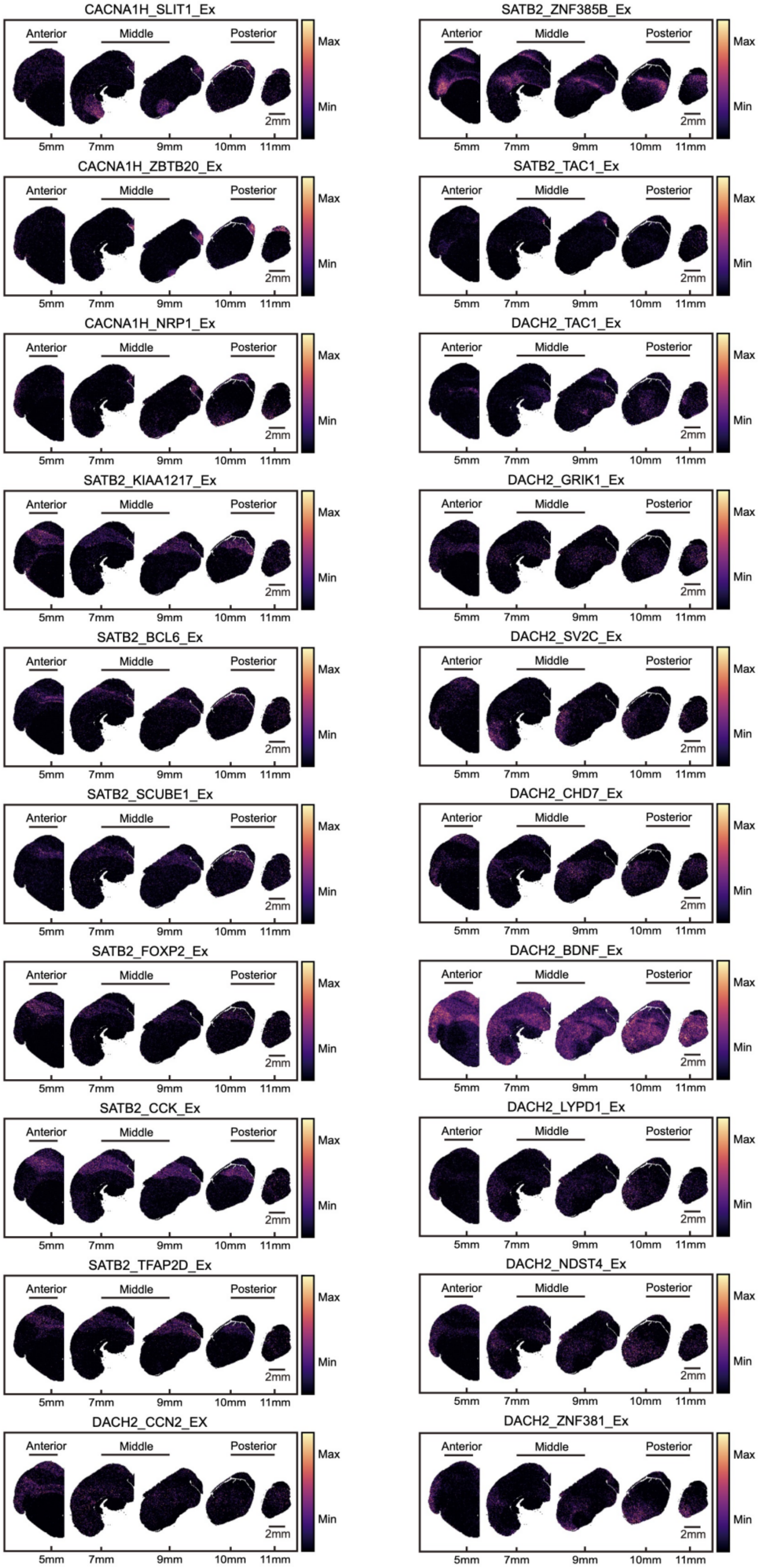

**Fig. S25. Spatial distribution of telencephalic excitatory neuronal supertypes.**
Spatial distribution patterns of excitatory neuronal supertypes in the budgerigar telencephalon, inferred using RCTD deconvolution across anterior, middle, and posterior coronal sections. Each panel shows relative spatial enrichment of a given supertype, annotated by subclass markers such as *SATB2*, *DACH2*, and *CACNA1H*.

#### Supplementary Figure 26

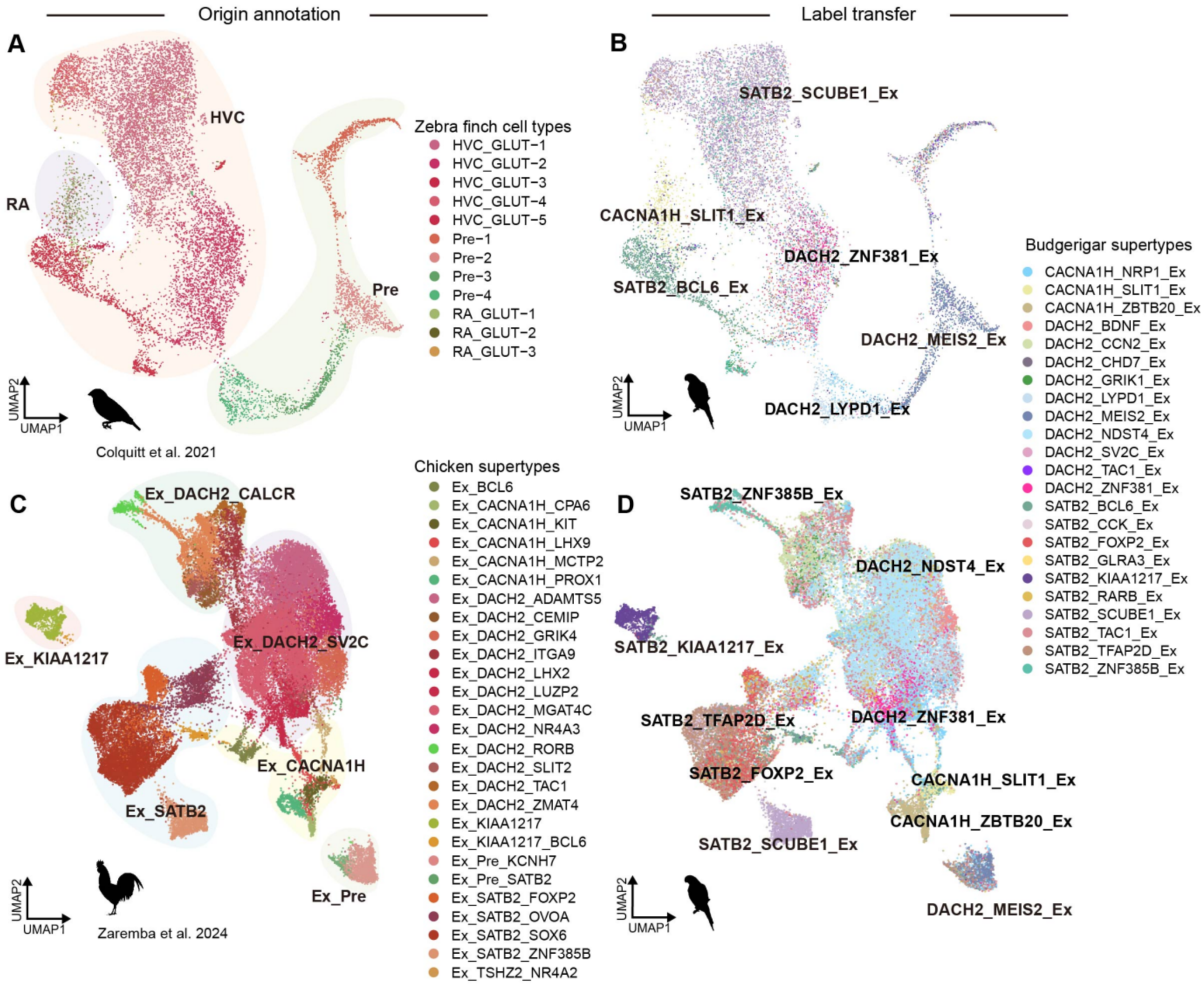


**E**

| Species | zebra finch | chicken_Henrik Kaessmann | chicken_Stein Aerts | budgerigar |
|---|---|---|---|---|
| Cell type | RA_GLUT1/2/3 | Ex_CACNA1H_KIT | EXC GLU-6b | CACNA1H_SLIT1_Ex |
| | HVC_GLUT-1/4 | Ex_SATB2_ZNF385B | EXC GLU-5a | SATB2_SCUBE1_Ex |
| | HVC_GLUT-2 | Ex_DACH2_LHX2/LUZP2 | EXC GLU-1a-2 | DACH2_ZNF381_Ex |
| | HVC_GLUT-3 | Ex_KIAA1217_BCL6 | EXC GLU-7-1 | SATB2_BCL6_Ex |
| | HVC_GLUT-5 | Ex_DACH2_RORB | EXC GLU-3 | SATB2_ZNF385B_Ex |
| | Pre-4 | ? | ? | DACH2_LYPD1_Ex |
| | Pre-2 | Ex_Pre_KCNH7 | EXC IMN | DACH2_MEIS2_Ex |
| | | Ex_KIAA1217 | EXC GLU-7-2 | SATB2_KIAA1217_Ex |
| | | Ex_DACH2_NR4A3 | EXC GLU-1b-2 | DACH2_BDNF_Ex |
| | | Ex_DACH2_ZMAT4 | EXC GLU-2 | DACH2_CCN2_Ex |
| | | Ex_DACH2_ADAMTS5/GRIK4 | EXC GLU-1b-1 | DACH2_NDST4_Ex |
| | | Ex_DACH2_MGAT4C | EXC GLU-1a-1/3 | DACH2_NDST4_Ex |
| | | Ex_DACH2_TAC1 | EXC GLU-2a-3 | SATB2_TAC1_Ex |
| | | Ex_CACNA1H_MCTP2/LHX9 | EXC GLU-6b | CACNA1H_NRP1_Ex |
| | | Ex_CACNA1H_CPA6/PROX1 | EXC GLU-6a | CACNA1H_ZBTB20_Ex |
| | | Ex_SATB2_SOX6/FOXP2 | EXC GLU-4-1/3, EXC GLU-5b | SATB2_SOX6_Ex |

**Fig. S26. Evolutionary comparison of telencephalic excitatory neuron types across avian species.**

(**A**) UMAP projection of telencephalic neurons from the zebra finch dataset (*7*), annotated by original region- and type-specific glutamatergic populations from the HVC and RA. (**B**) Label transfer of budgerigar-defined excitatory neuronal supertypes onto the zebra finch dataset, revealing conserved subtype-specific expression patterns. (**C**) UMAP of chicken telencephalic excitatory neurons (*18*) with original annotations based on gene-defined supertypes. (**D**) Cross-species projection of budgerigar excitatory supertypes onto chicken, showing molecular correspondence to *SATB2*, *CACNA1H*, *DACH2*, and *KIAA1217* subclasses. (**E**) Cross-species alignment table comparing excitatory neuron identities in zebra finch (*7*), chicken (*18, 27*) and budgerigar. Cell types are matched based on transcriptomic similarity and marker gene homology.

Supplementary Figure 27

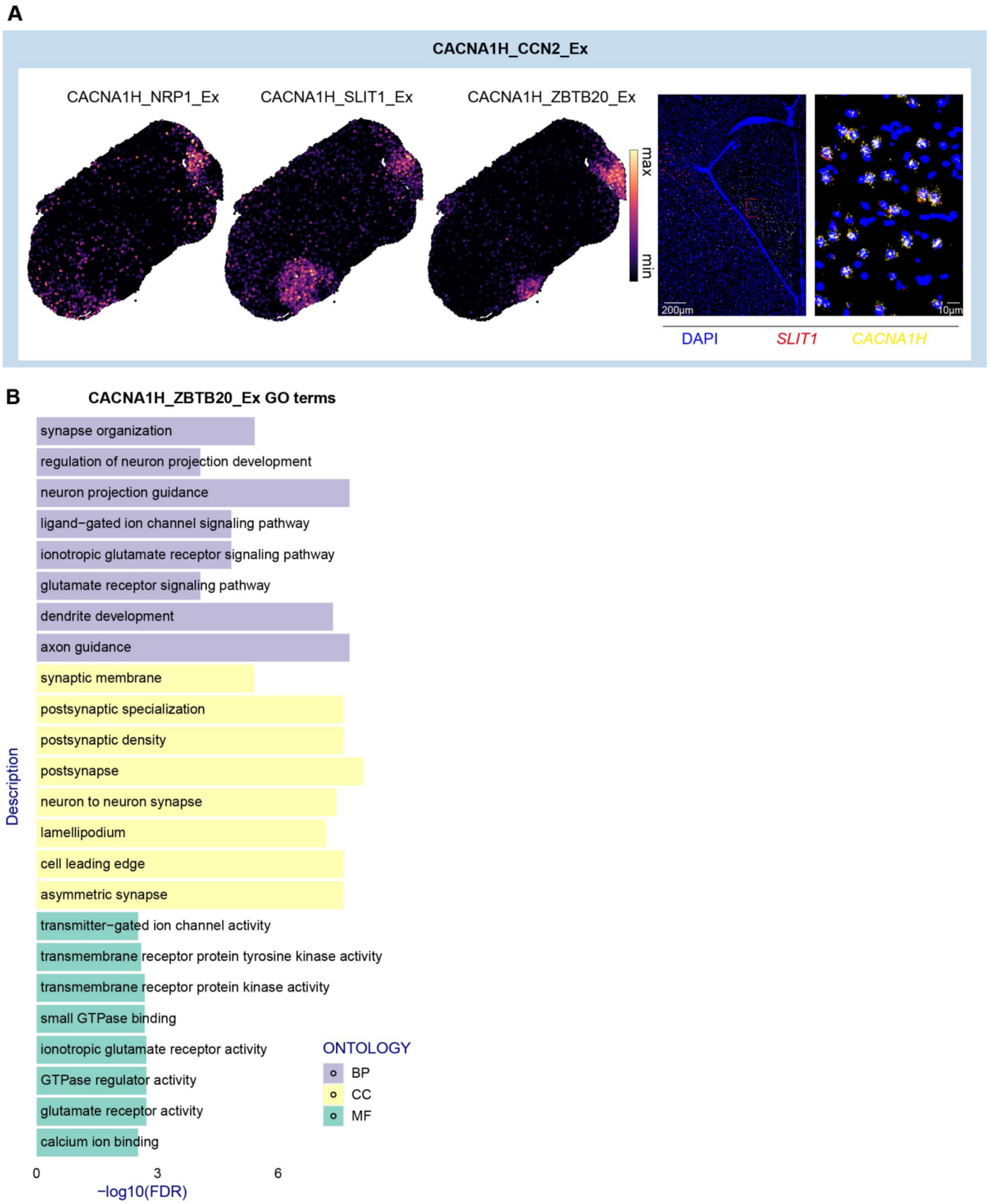


**Fig. S27. Spatial localization and functional annotation of CACNA1H-expressing excitatory supertypes.**

(**A**) Left: Spatial distribution of CACNA1H_CCN2 subclass (CACNA1H_NRP1_Ex, CACNA1H_SLIT1_Ex, CACNA1H_ZBTB20_Ex) in the telencephalon, inferred by RCTD. Right: HCR RNA-FISH images showing co-localization of *CACNA1H* and *SLIT1* expression. (**B**) GO enrichment analysis of DEGs in CACNA1H_ZBTB20_Ex. Functional terms highlight roles in synapse organization, neuron projection guidance, and ionotropic glutamate signaling.

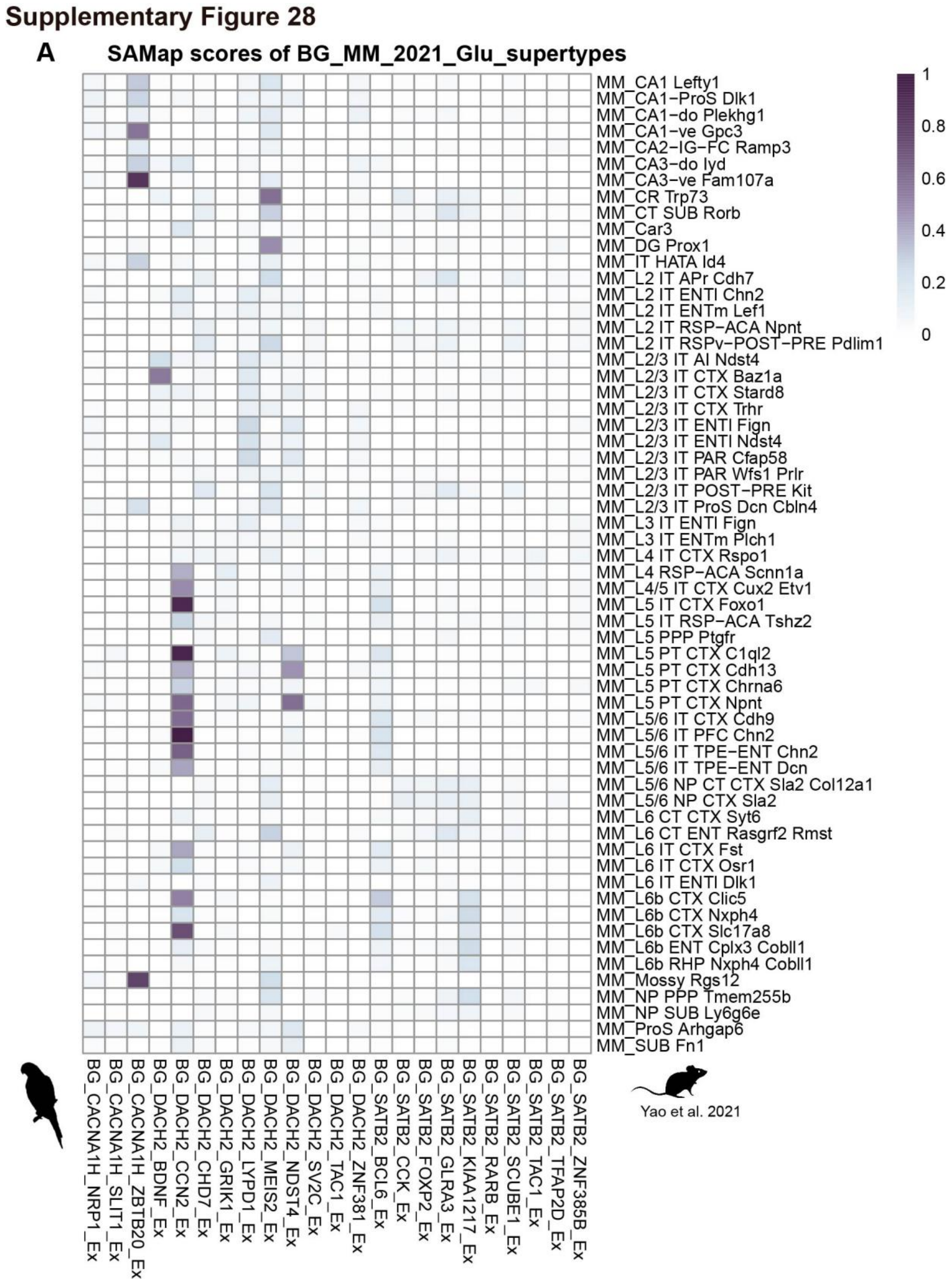


**Fig. S28. Cross-species comparison of excitatory neuron supertypes between budgerigar telencephalon and mouse isocortex.**

(**A**) SAMap analysis of glutamatergic neuronal supertypes from the budgerigar telencephalon and the mouse cortex (*12*). Heatmap displays transcriptomic similarity scores between each budgerigar supertype (bottom axis) and annotated mouse excitatory populations (right axis), including layer-specific cortical, hippocampal, and subplate neurons.

Supplementary Figure 29

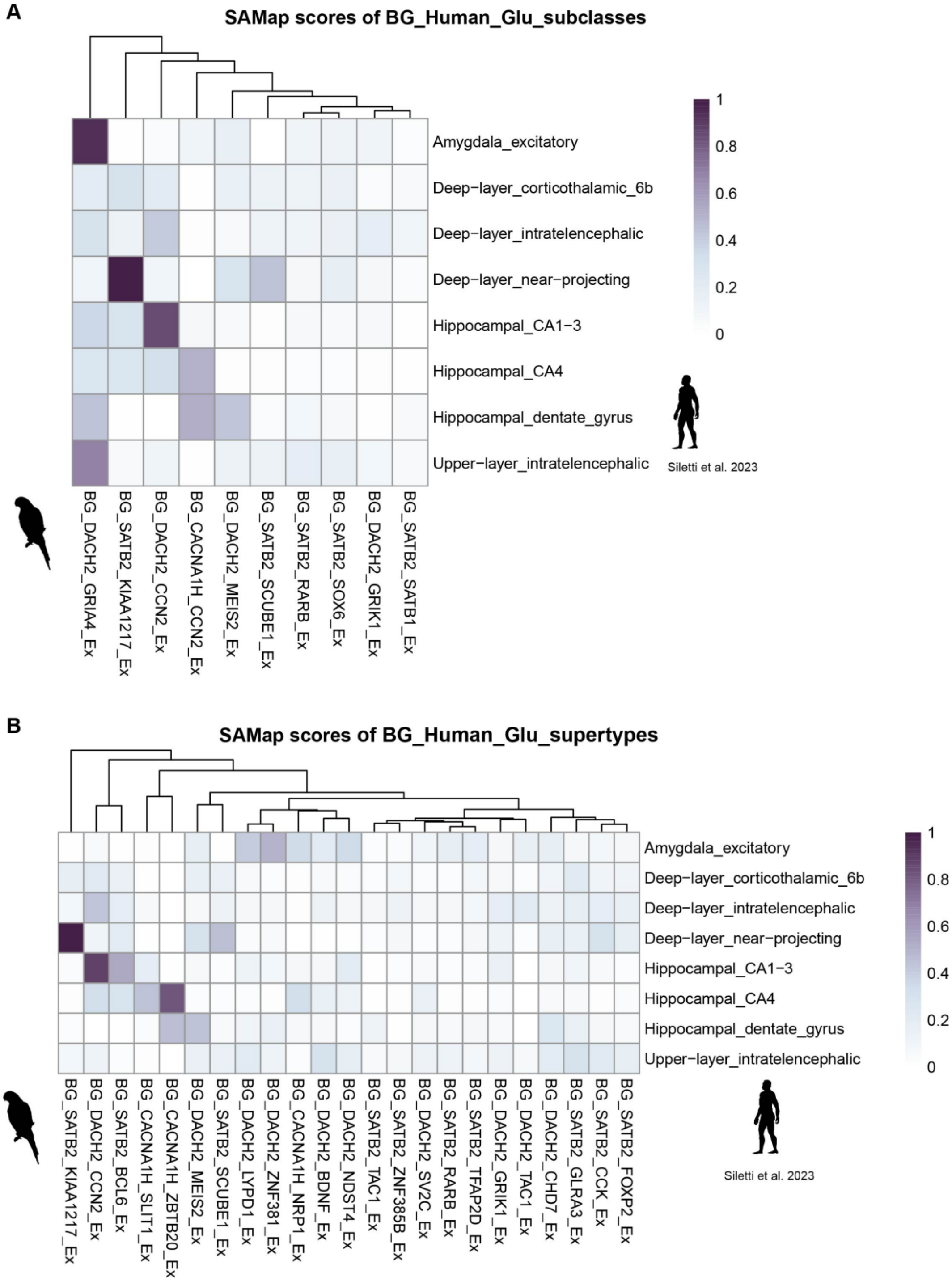


**Fig. S29. Cross-species comparison of excitatory neuron types between human cortex and budgerigar telencephalon.**

(**A**) SAMap analysis of transcriptomic similarity between human excitatory neuron subclasses (*17*) and budgerigar telencephalic subclasses. (**B**) SAMap alignment between human excitatory neuron subclasses and budgerigar excitatory neuronal supertypes. Heatmaps show matched similarity scores, highlighting evolutionary relationships between upper-layer, deep-layer, and hippocampal neuron types across species.

#### Supplementary Figure 30

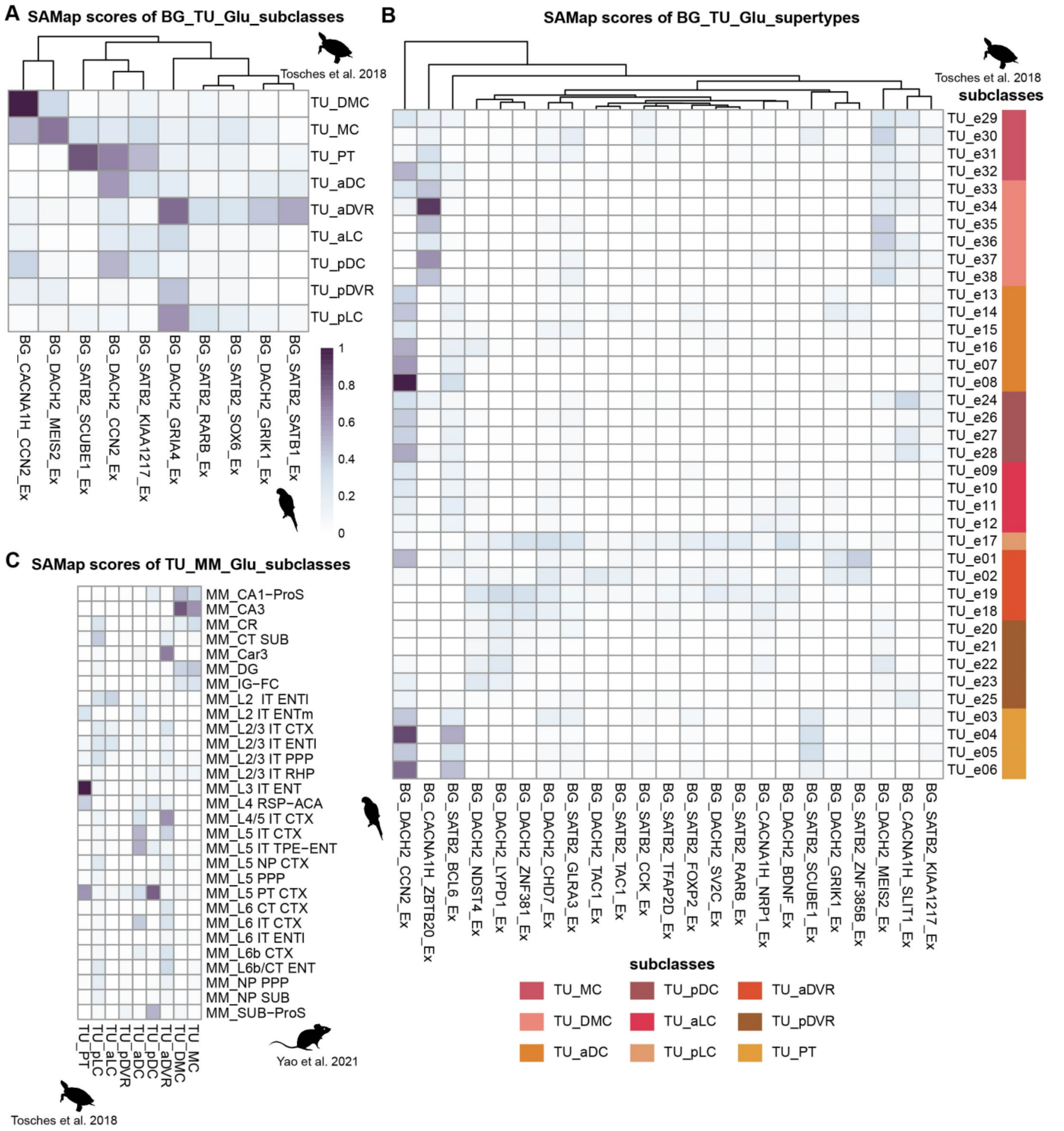


**Fig. S30. Cross-species comparison of telencephalic excitatory neurons between budgerigar, turtle, and mouse.**
(**A**) SAMap similarity scores between budgerigar excitatory neuron subclasses and major turtle telencephalic regions (*14*). (**B**) SAMap alignment between budgerigar excitatory supertypes and turtle excitatory subtypes, revealing molecular correspondences across pallial domains. (**C**) SAMap comparison of turtle telencephalic regions with mouse excitatory subclasses from isocortex, hippocampus, and subiculum (*12*), enabling cross-referencing of conserved cell identities among sauropsids and mammals.

Supplementary Figure 31

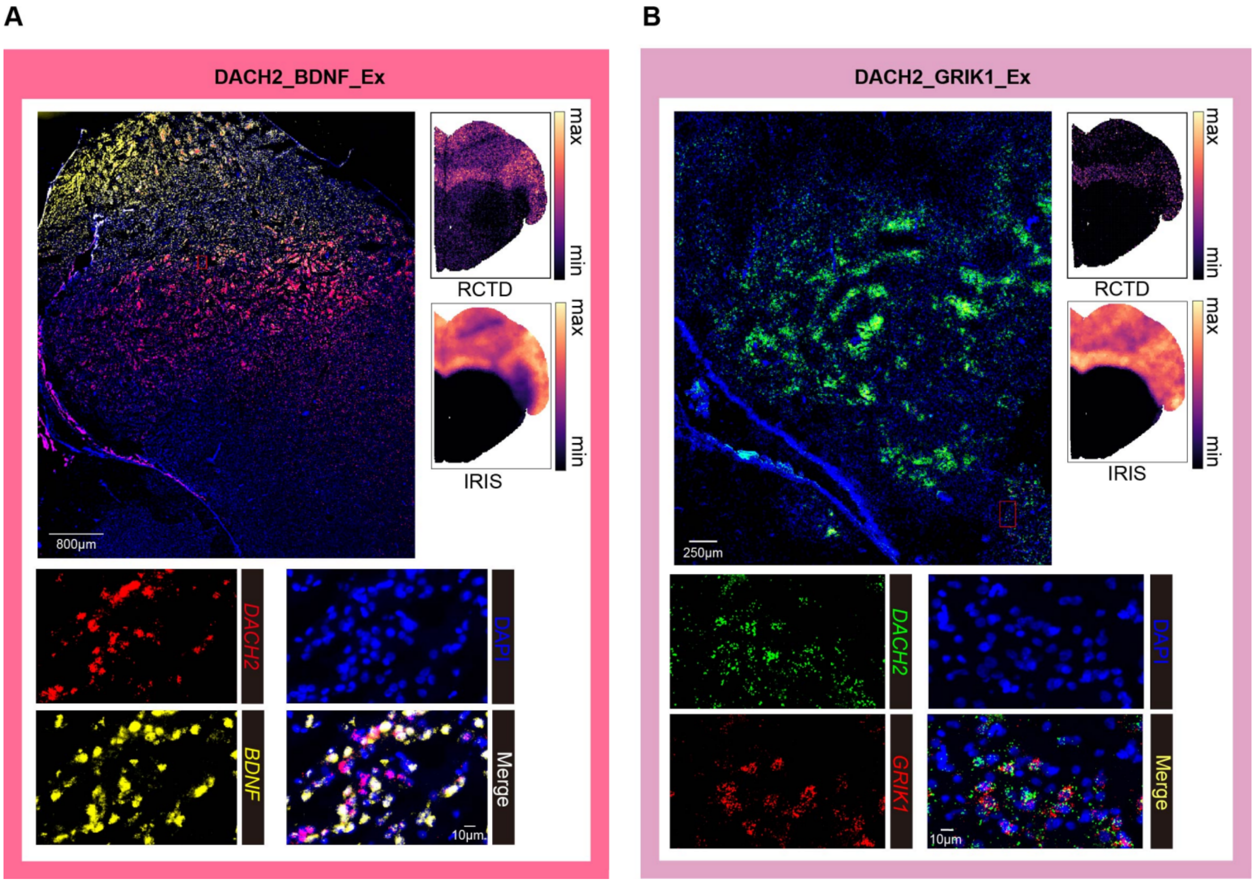


**Fig. S31. Spatial validation of telencephalic *DACH2* supertypes by HCR RNA-FISH.**
(**A**) Spatial localization of DACH2_BDNF_Ex neurons inferred by RCTD and IRIS deconvolution. Corresponding HCR RNA-FISH shows co-expression of *DACH2* and *BDNF* in the telencephalon. (**B**) Spatial localization of DACH2_GRIK1_Ex neurons, validated by co-localized expression of *DACH2* and *GRIK1*. In both panels, upper right insets show predicted spatial maps (RCTD and IRIS), and bottom insets show high-resolution fluorescence images confirming supertype identity.

Supplementary Figure 32

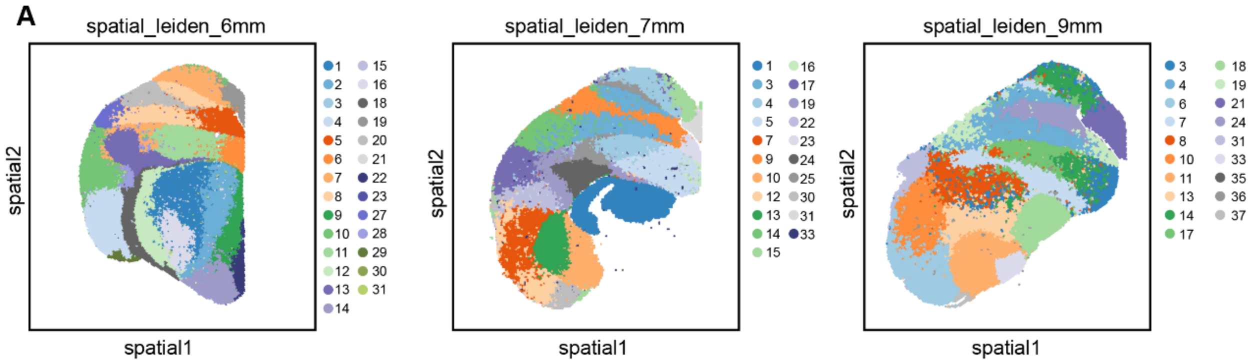


**Fig. S32. Spatial clustering of the budgerigar telencephalon at different anterior–posterior levels.**

(**A**) Spatial Leiden clustering of coronal sections at 6 mm, 7 mm, and 9 mm posterior to the olfactory bulb (resolution = 2). The telencephalon (comprising pallial and subpallial domains) was subset for regional analysis.

#### Supplementary Figure 33

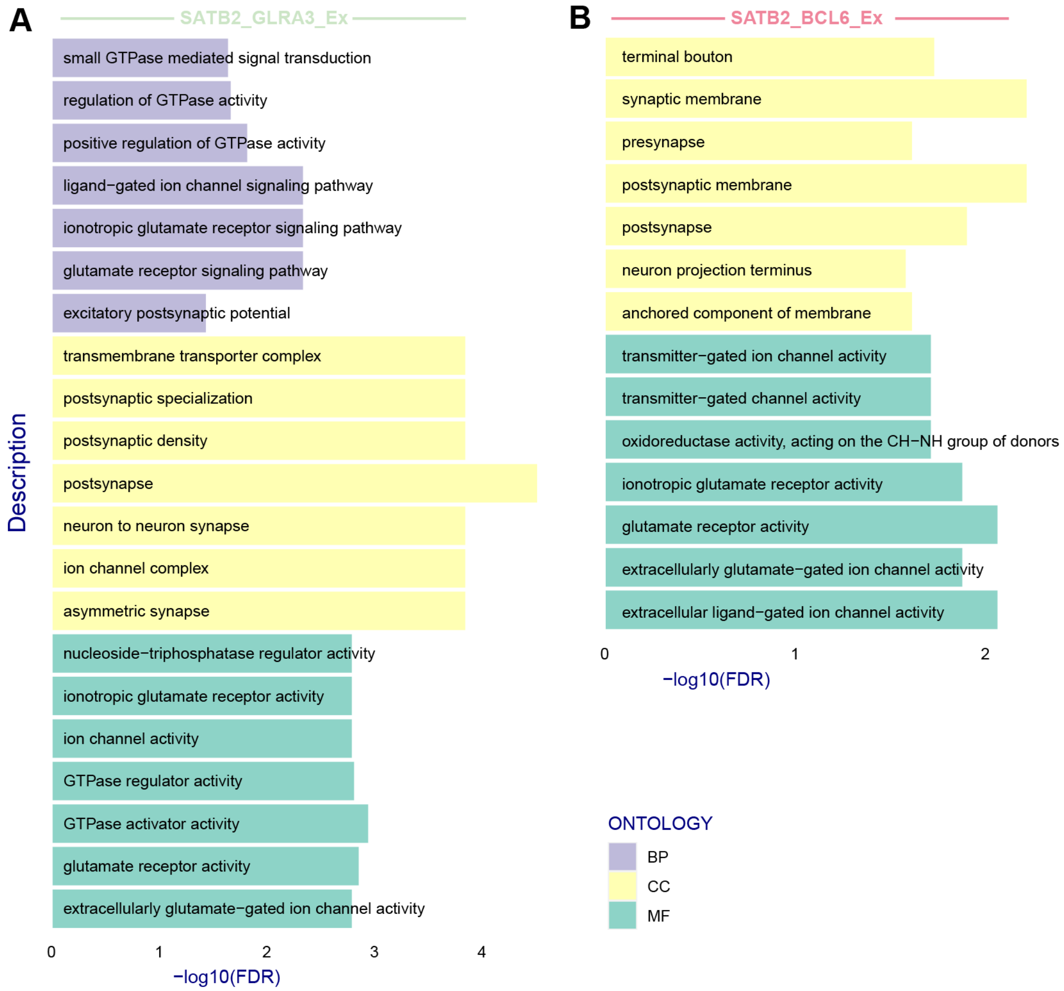


**Fig. S33. Molecular and functional divergence of excitatory neuron supertypes with complementary lamina mesopallialis intermedia (LMI) distribution patterns.**
(**A**) GO enrichment analysis of the *SATB2*$^{+}$ *GLRA3*$^{+}$ excitatory neuron supertype, which shows exclusion from the LMI region. (**B**) GO enrichment analysis of the *SATB2*$^{+}$ *BCL6*$^{+}$ excitatory neuron supertype, which exhibits specific localization within the LMI.

Supplementary Figure 34

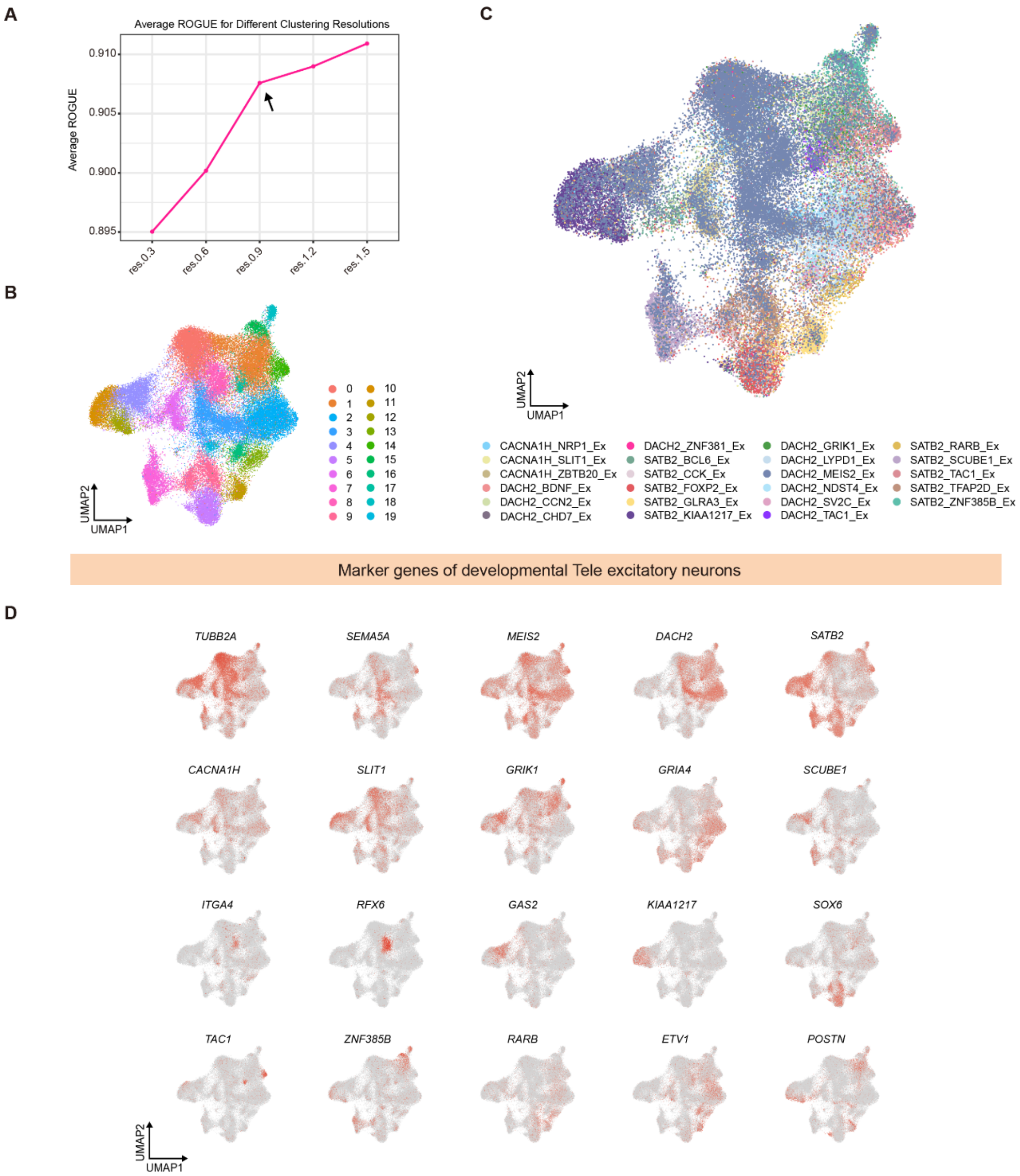


**Fig. S34. Identification of excitatory neuronal supertypes in the developing budgerigar telencephalon.**

(**A**) ROGUE-based evaluation of clustering resolutions to optimize developmental supertype classification. (**B**) UMAP of clusters at optimal resolution (0.9). (**C**) Annotation of developmental clusters by label transfer from adult excitatory supertypes. (**D**) Expression of supertype-specific marker genes in developmental excitatory neurons.

#### Supplementary Figure 35

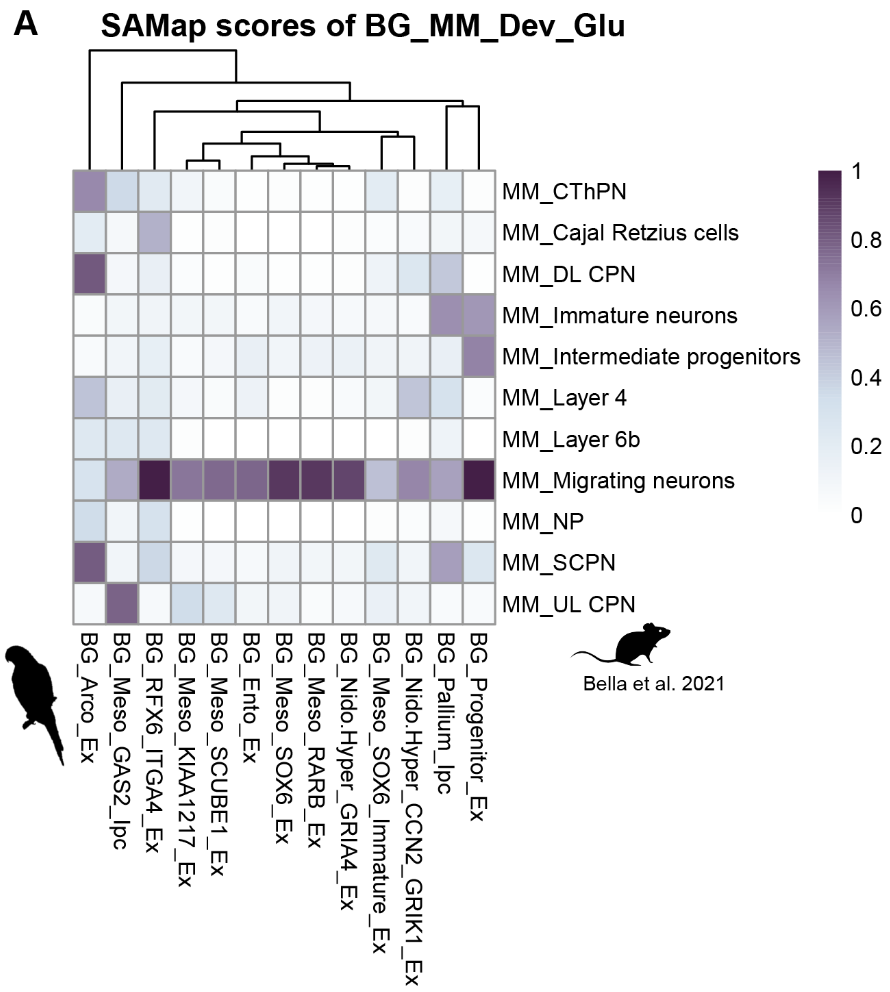


**Fig. S35. Cross-species comparison of developing excitatory neurons between budgerigar and mouse neocortex.**

(**A**) SAMap analysis comparing E14 excitatory neuron types in the budgerigar telencephalon and mouse neocortex [subset of (*15*) embryonic days 16 to 18].

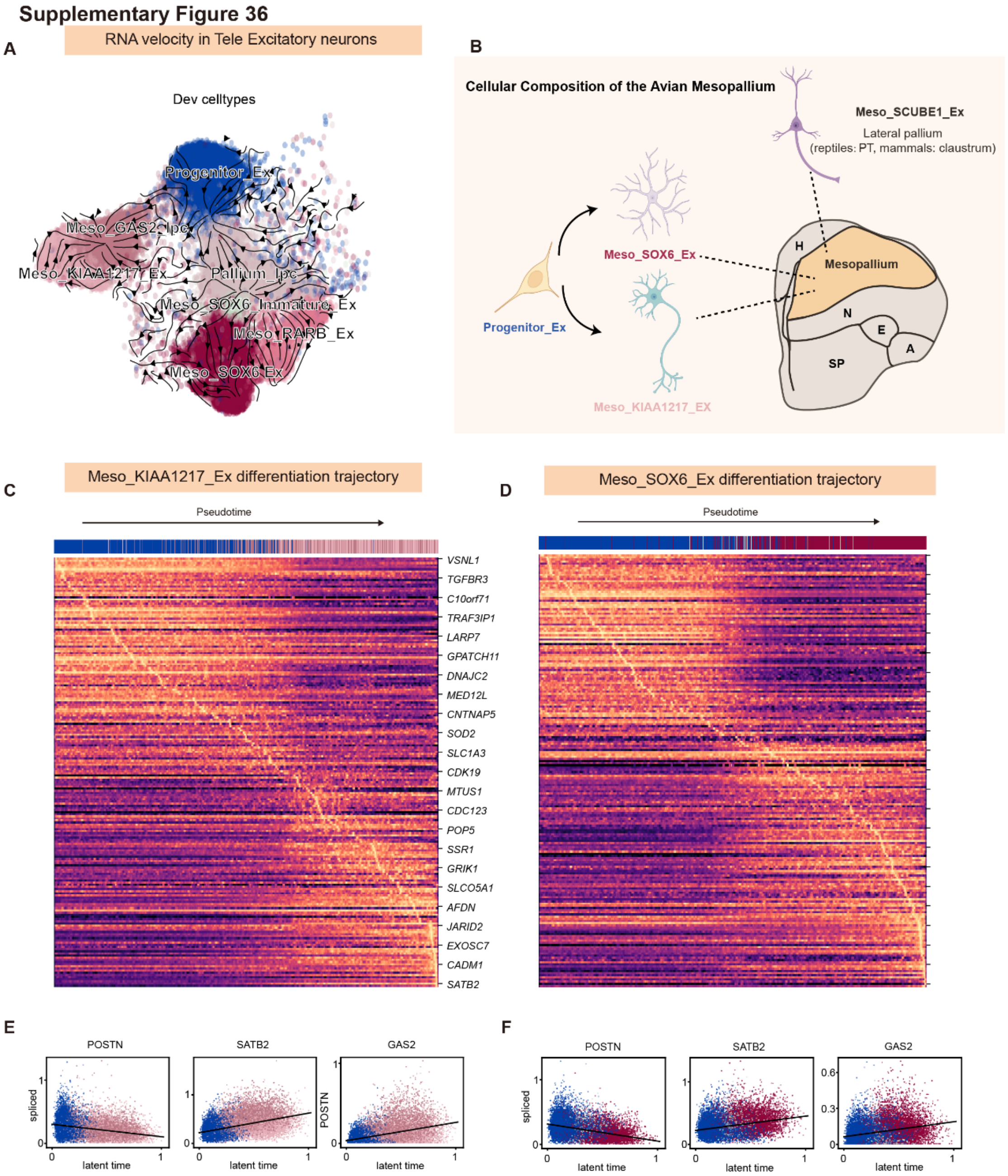


**Fig. S36. Developmental trajectories of mesopallial excitatory neuron subclasses.**
(**A**) RNA velocity analysis reveals lineage trajectories of telencephalic excitatory neurons, with Meso_KIAA1217_Ex and Meso_SOX6_Ex derived from Progenitor_Ex. (**B**) Schematic of mesopallial composition. Meso_KIAA1217_Ex and Meso_SOX6_Ex arise from pallial progenitors, while Meso_SCUBE1_Ex originates from the lateral pallium lineage. (**C, D**) Gene expression dynamics across pseudotime for Meso_KIAA1217_Ex (**C**) and Meso_SOX6_Ex (**D**) trajectories. (**E, F**) Latent time plots showing expression trends of *POSTN*, *SATB2*, and *GAS2* during differentiation of Meso_KIAA1217_Ex (**E**) and Meso_SOX6_Ex (**F**).

Supplementary Figure 37

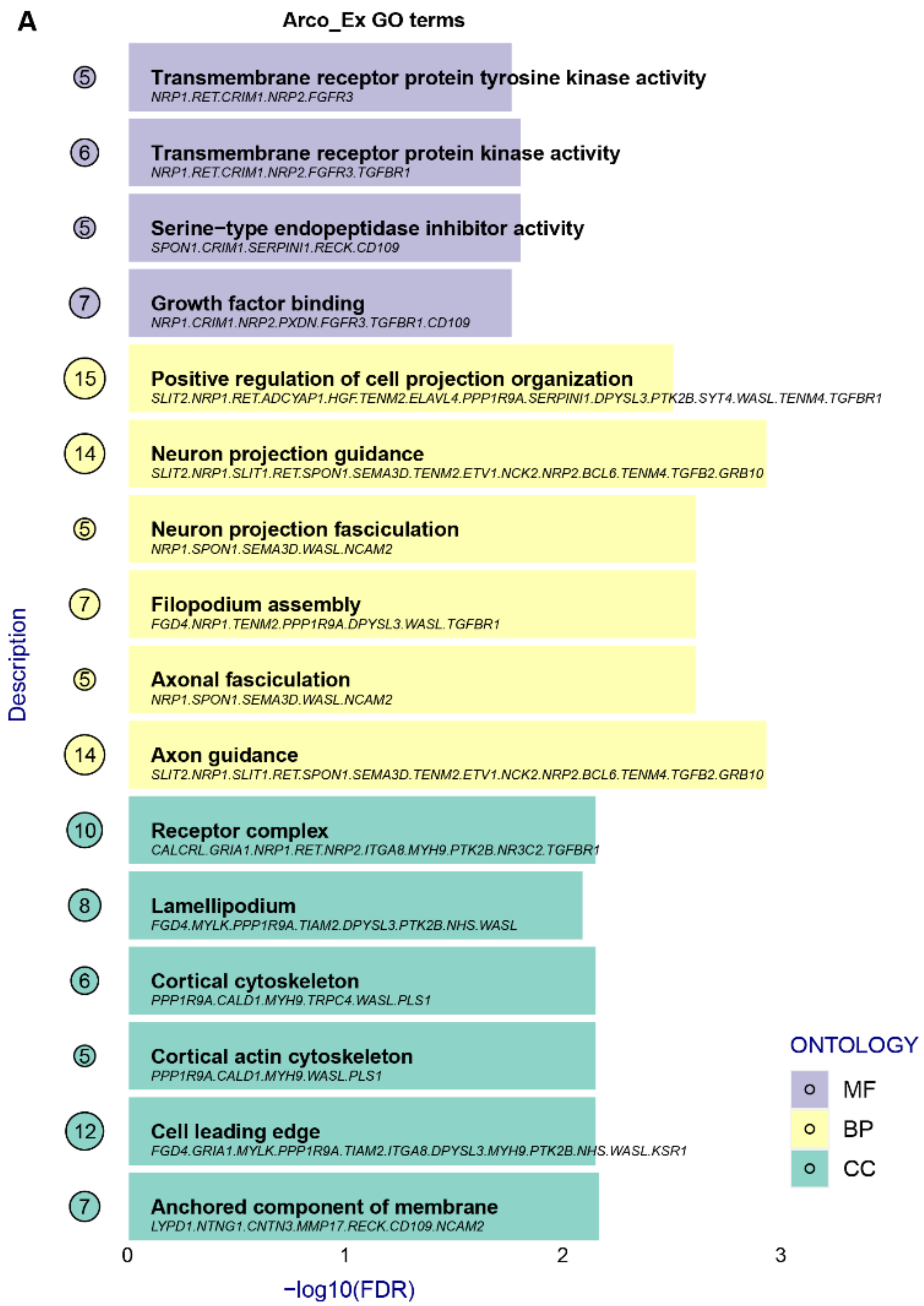


**Fig. S37. Functional enrichment analysis of Arco_Ex excitatory neurons.**
(**A**) GO terms enriched in *Arco_Ex* neurons, highlighting molecular functions (MF), biological processes (BP), and cellular components (CC) associated with axon guidance, cytoskeletal dynamics, and growth factor signaling.

#### Supplementary Figure 38

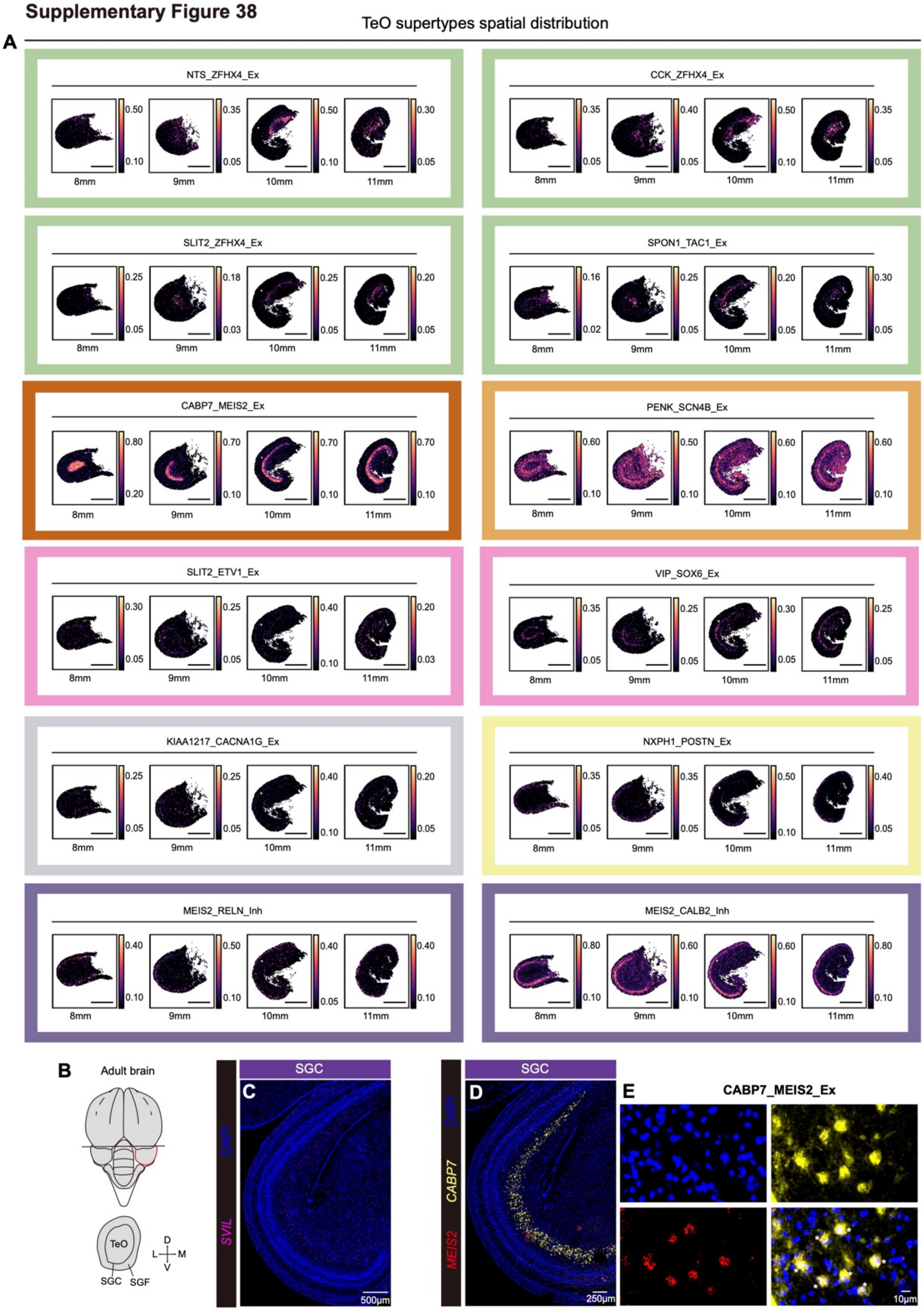

**Fig. S38. Spatial organization and marker validation of optic tectum (TeO) excitatory neuron supertypes.**

(**A**) Spatial distributions of excitatory and inhibitory TeO supertypes across coronal sections (8–11 mm), inferred by RCTD. Colored frames group supertypes by laminar domain (e.g., SGP, SGC, SGF). (**B**) Schematic of adult tectal anatomy and lamination. (**C to E**) HCR RNA-FISH validation of TeO marker genes. (**C**) Layer-specific expression of *SVIL*. (**D**) Co-localization of *CABP7* and *MEIS2* in the SGC. (**E**) High-magnification views confirming *CABP7_MEIS2_Ex* identity via dual-marker expression.

Supplementary Figure 39

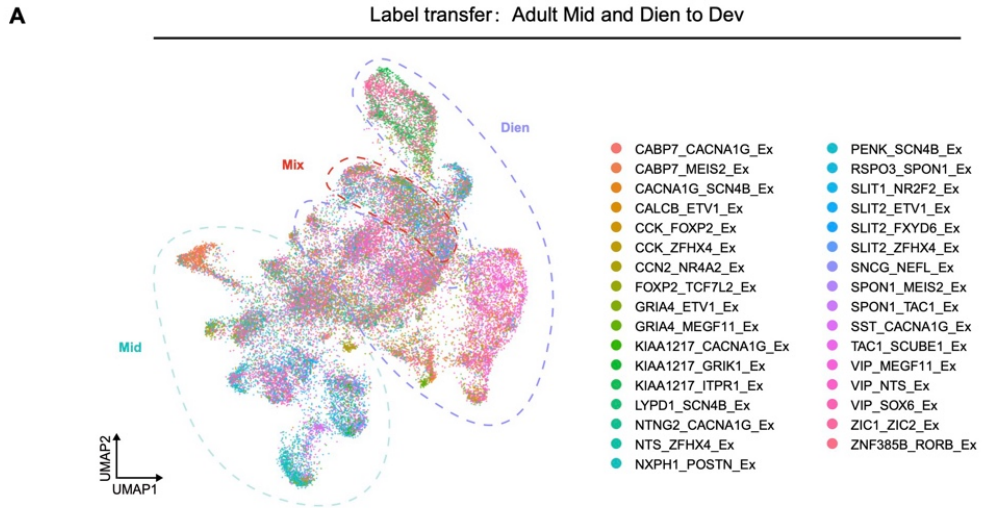


**Fig. S39. Label transfer from adult midbrain and diencephalon excitatory supertypes to developmental datasets.**

(**A**) UMAP showing predicted developmental distribution of adult-defined excitatory neuron supertypes from the midbrain (Mid), diencephalon (Dien), and mixed regions (Mix). Labels were transferred to developmental single-cell data to infer spatial and lineage origin.

#### Supplementary Figure 40

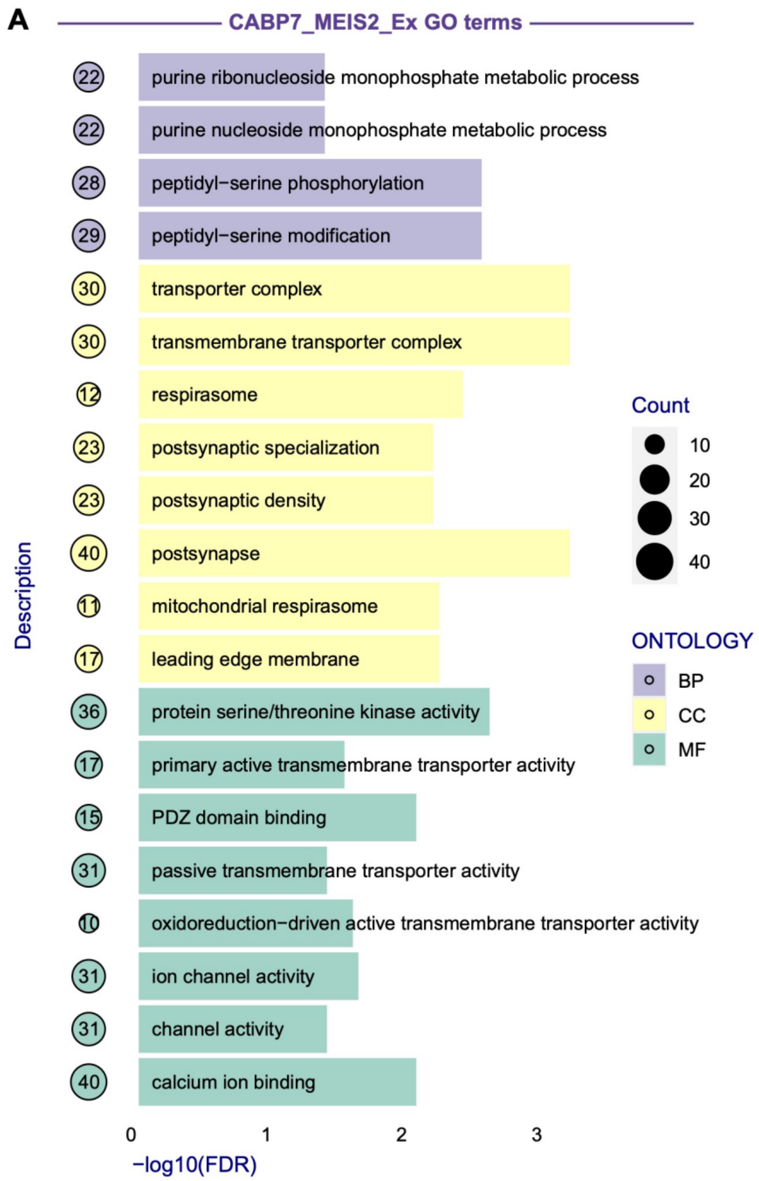


**Fig. S40. Functional enrichment of CABP7_MEIS2_Ex excitatory neurons.**
(**A**) GO analysis of CABP7_MEIS2_Ex subtype reveals enrichment in synaptic specialization, ion transport, mitochondrial respiration, and phosphorylation-related pathways.

**Supplementary Figure 41**

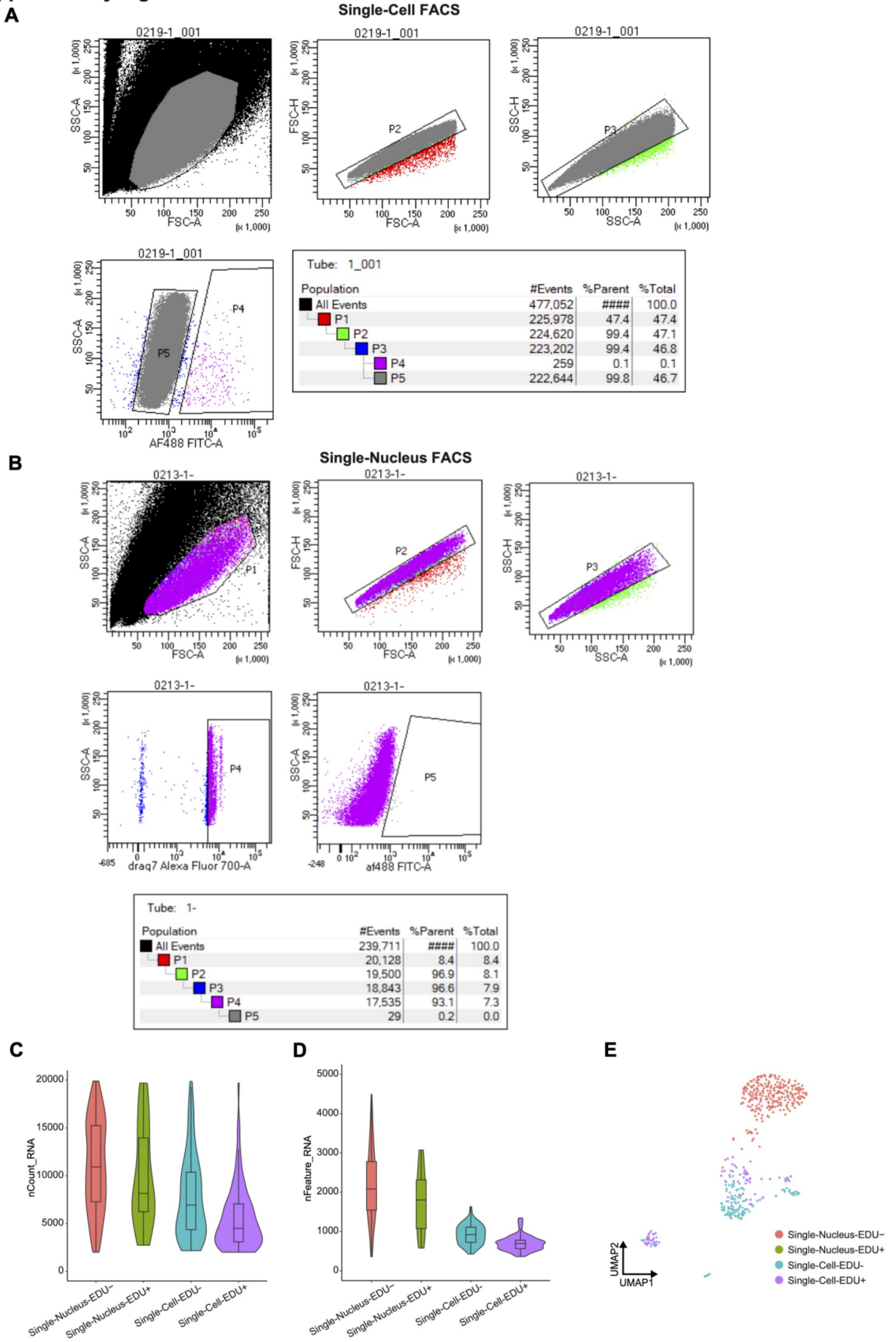


**Fig. S41. FACS gating and library quality assessment of EdU-labeled single cells and nuclei.** (**A**) Flow cytometry gating strategy for isolating EdU$^{+}$ and EdU$^{-}$ single cells from budgerigar brain tissue. (**B**) Gating strategy for isolating EdU$^{+}$ and EdU$^{-}$ single nuclei. (**C, D)** Violin plots showing RNA molecule (nCount_RNA, **C**) and gene (nFeature_RNA, **D**) recovery across four sorted populations. (**E**) UMAP plot showing transcriptional clustering of $EdU^+$ and $EdU^-$ cells from both isolation strategies.

Supplementary Figure 42

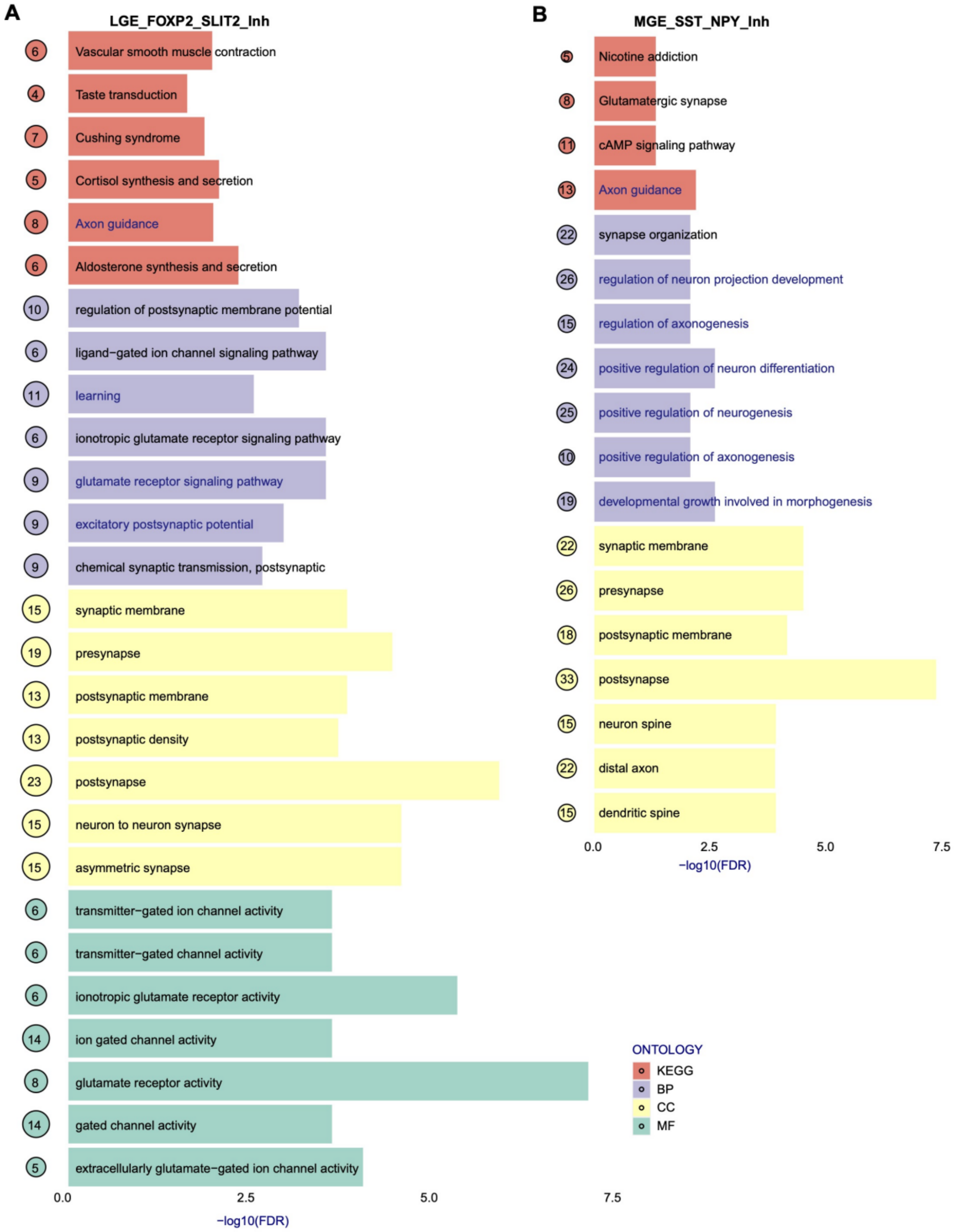


**Fig. S42. Functional enrichment of inhibitory neuron supertypes derived from adult neurogenesis.**

(**A**) GO and KEGG enrichment for LGE-derived FOXP2_SLIT2_Inh supertype highlights pathways involved in synaptic transmission, ion channel activity, and axon guidance. (**B**) Enrichment for MGE-derived SST_NPY_Inh supertype reveals roles in neuron projection, synaptic organization, and neurodevelopmental regulation.

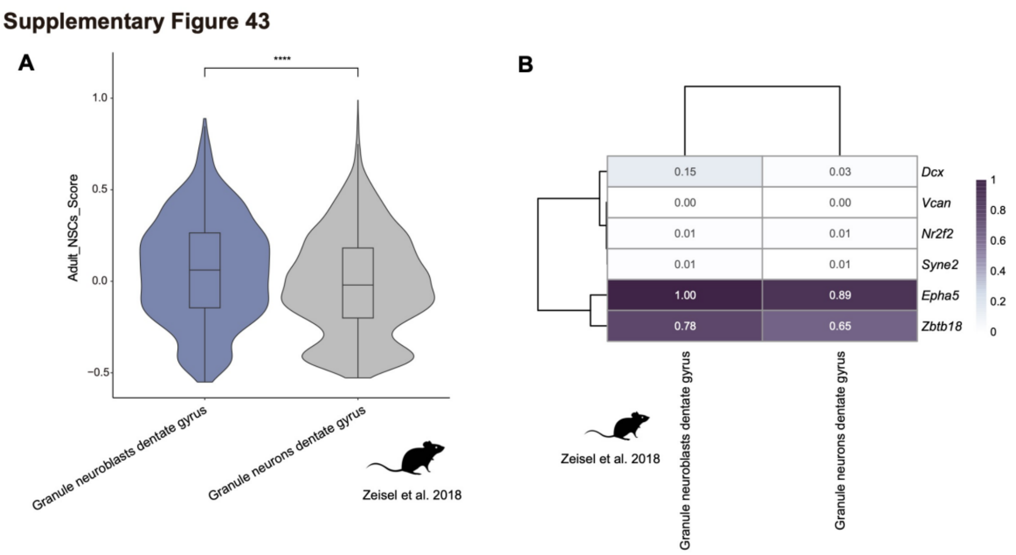


**Fig. S43. Avian NSC signatures predict neuroblast identity in the mouse dentate gyrus.**
(**A**) Violin plot showing significantly higher adult NSC scores in granule neuroblasts versus mature granule neurons in the mouse dentate gyrus (**** $p < 0.0001$). (**B**) Heatmap of representative avian NSC marker gene expression (e.g., *Epha5*, *Zbtb18*) in mouse neuroblast and neuron clusters (data from (*28*)).

**Supplementary Figure 44**

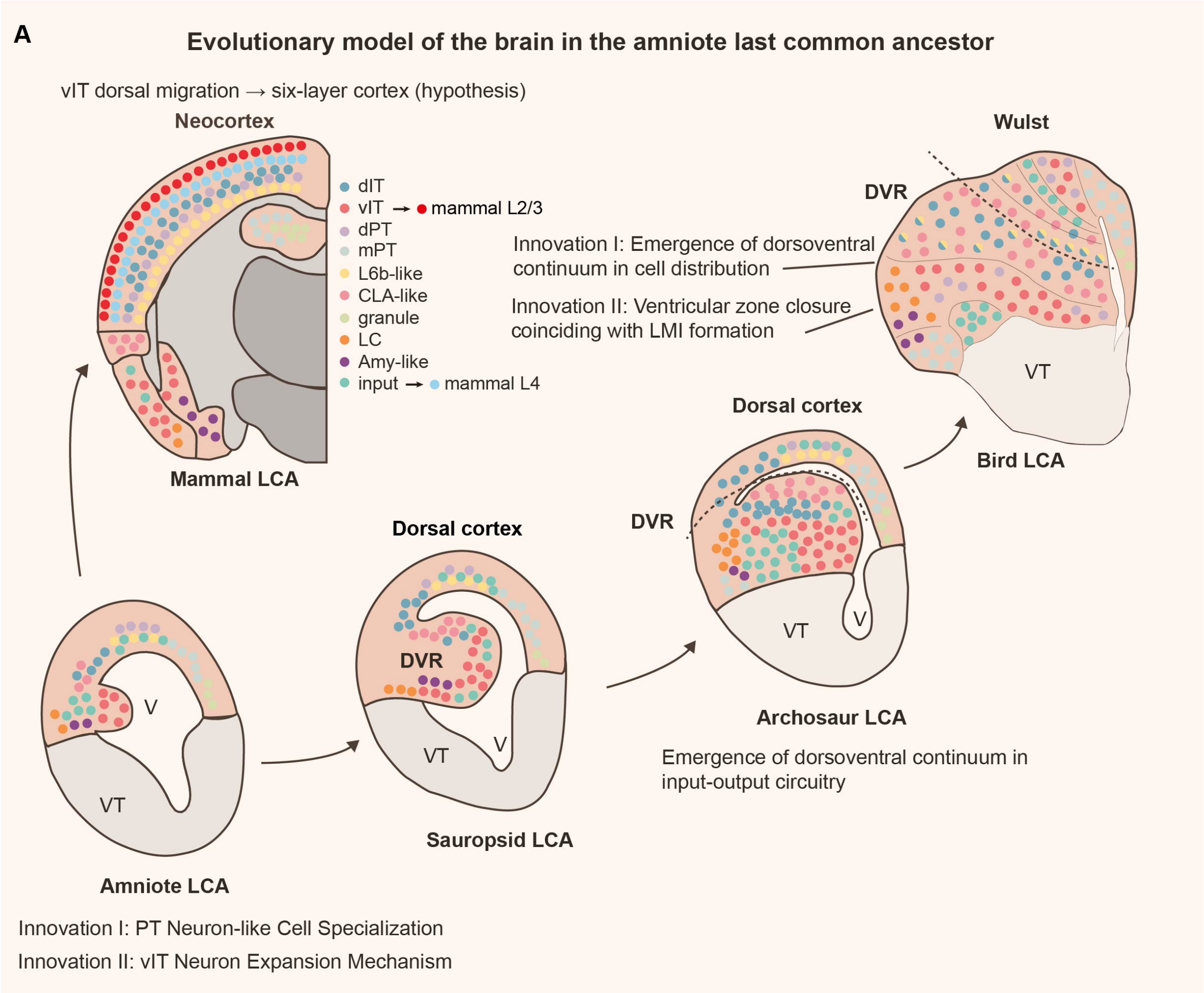


**Fig. S44. Evolutionary model of dorsoventral circuitry integration in the amniote pallium.** Schematic presents a unified framework for pallial evolution from the amniote last common ancestor (LCA) to modern lineages, depicting two fundamental innovations in the LCA: (i) specialization of PT neuron-like projection cells and (ii) expansion of ventral-derived intratelencephalic (vIT) neurons. In mammals, dorsal migration of vIT populations contributed to six-layered neocortex formation (*29*), ultimately differentiating into layer 2/3 IT neurons. The archosaur lineage instead developed a dorsoventral continuum circuitry (*30*), with birds exhibiting two derived features: (i) a graded cell distribution along the dorsoventral axis and (ii) ventricular zone closure coinciding with LMI formation (*31, 32*). The model establishes an evolutionary continuum linking mammalian cortical laminar organization with avian DVR circuits through conserved neuronal classes, suggesting shared developmental mechanisms despite divergent morphological outcomes.

#### Supplementary Figure 45

A

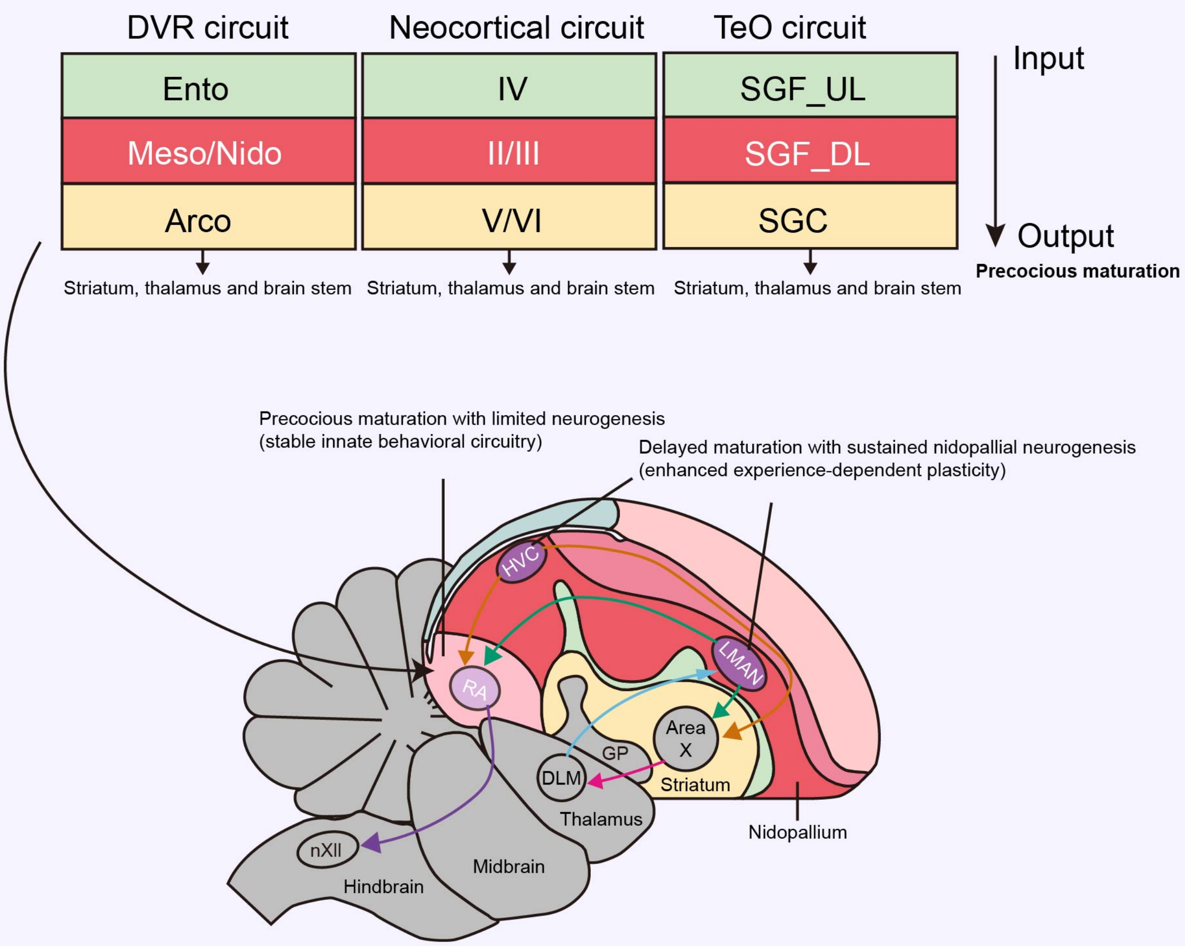


B

| Brain subdivision | Auditory parrot | Auditory songbird | Vocal parrot | Vocal songbird |
|---|---|---|---|---|
| Hyperpallium | CMHV | CMHV | HVo | HVo |
| | | | 1AHV (?) | Av (?) |
| Mesopallium/Ni-dopallium | NCM, L1, L3 | NCM, L1, L3 | NLC | HVC |
| | NIDL | HVC shelf | NAO | MAN |
| Arcopallium | ACM | RA CuP | AAC | RA |
| Striatum | PC | PC | LPOm | Area X |
| Thalamus | Ov(-) | Ov (-) | DMm | DLM (+/-) |
| Midbrain | MLd | MLd | DM | DM |

**Fig. S45. Pioneer and reserve hypothesis.**
(**A**) Schematic showing a comparison of DVR, neocortical, and TeO circuit architectures between birds and mammals. Avian input–output circuits mirror neocortical layering: the entopallium or SGF_UL corresponds to input layer IV, mesopallial and nidopallial regions correspond to superficial layers (II/III), and the arcopallium or SGC corresponds to deeper layers (V/VI) (*33, 34*). Notably, output regions consistently exhibit precocious maturation. The avian song system exemplifies the developmental and neurogenic dynamics of pallial excitatory neurons: the robust nucleus of the arcopallium (RA) matures early with minimal adult neurogenesis, whereas delayed maturation accompanied by sustained neurogenesis in the hyperpallium, nidopallium, and mesopallium likely supports enhanced plasticity. (**B**) Comparative mapping of auditory and vocal control brain regions across parrots and songbirds. Although terminology differs from that of Jarvis *et al.* (*35*), homologous functions are distributed across similar pallial subdivisions.

**Data S1.**
Sample information

**Data S2.**
DNA probe sequences used for HCR RNA-FISH

**Data S3.**
Adult budgerigar neuronal supertype marker genes

**Data S4.**
Developmental neuronal subtype marker genes

**Data S5.**
Differentially expressed genes in adult NSCs
